# Advancing Ligand Binding Affinity Prediction with Cartesian Tensor-Based Deep Learning

**DOI:** 10.1101/2025.06.04.657800

**Authors:** Jie Yu, Xia Sheng, Zhehuan Fan, Zhaokun Wang, Duanhua Cao, Yongxin Hao, Yingying Zhang, Panpan Shao, Huicong Ma, Tian Cao, JingXin Rao, Mingan Chen, Kaixian Chen, Xutong Li, Dan Teng, Xiaomin Luo, Mingliang Wang, Sulin Zhang, Mingyue Zheng

**Author notes:** **Corresponding Author** (Mingliang Wang), (Sulin Zhang), (Mingyue Zheng). J.Y., X.S., Z.H.F., Z.K.W., and D.H.C. contributed equally to this work. **Author Contributions** M.Y.Z. and J.Y. designed the research study. J.Y. and X.S. developed the method and wrote the code. M.L.W., H.C.M, and T.C. conducted chemical synthesis. All authors contributed to the analysis of the results. J.Y., M.Y.Z. and X.S. wrote the paper. All authors read and approved the manuscript. **Note** The authors declare no competing financial interest.

## Abstract

We present PBCNet2.0, a cartesian tensor-based Siamese Neural Network for protein-ligand relative binding affinity prediction. Trained on 8.6 million protein-ligand complex structure pairs, PBCNet2.0 achieves zero-shot performance comparable to computationally intensive physics-based simulations. Our prioritization experiments show that PBCNet2.0 speeds up binding affinity optimization by 718% while reducing resource use by 41%. Through extensive retrospective experiments, we demonstrate that PBCNet2.0 intrinsically comprehends protein-ligand interactions, showing high sensitivity to intermolecular interactions and exceptional perception of spatial geometric information. Strikingly, PBCNet2.0 exhibits an emergent capability to predict affinity changes induced by binding residue variations, highlighting its potential for identifying resistance mutation. We prospectively validated these capabilities on two targets ENPP1 and ALDH1B1, where PBCNet2.0 successfully identified affinity shifts arising from subtle molecular interactions and conformational differences, and pinpointed critical binding residues with an 83% hit rate. This combination of computational efficiency, spatial geometric perception of binding site, and generalizable affinity prediction establishes PBCNet2.0 as a transformative tool for developing pharmacological probes for all human proteins.

## 1. Introduction

The human genome harbors a vast number of proteins that remain underexplored despite their potential therapeutic value^1,2^. Many of these proteins are poorly understood and have not been adequately studied, yet they hold significant promise for drug development. A key factor contributing to this underutilization is the lack of high-quality, well-characterized molecular probes^1^. Recognizing this gap, the global initiative Target 2035 has been launched with the ambitious goal of identifying pharmacological probes for all human proteins^3,4^.

Within this context, protein-ligand binding affinity prediction^5–7^ plays crucial roles in accelerating the development of molecular probes. While physics-based methods such as free energy perturbation (FEP, e.g., Schrödinger’s FEP+^8^) have achieved chemical accuracy (root-mean-square error ~1.1 kcal · mol^−1^) in predicting relative binding affinities, their substantial computational cost, complexity in system setup, and dependency on expert intervention limit their broad applicability^9^. Consequently, artificial intelligence (AI), especially deep learning, presents a compelling opportunity for high-throughput and accurate affinity predictions by learning directly from structural data.

Recent advances in deep learning models that incorporate three-dimensional (3D) structural information have demonstrated promising results in protein-ligand binding prediction^5,10^. Our group developed EquiScore, an equivariant graph neural network that integrates physical principles of ligand-protein interactions, showing robust performance in virtual screening tasks^5^. In parallel, Moon et al. developed PIGNet2.0, a physics-informed graph neural network that explicitly models various protein-ligand interactions to predict binding free energies, proving effective in both virtual screening and lead optimization^11^. We previously introduced the Pairwise Binding Comparison Network (PBCNet), which employs a Siamese Neural Network (SNN) architecture enhanced with physics-informed graph attention mechanisms to rank binding affinities among structurally similar ligands^6^. PBCNet processes pairs of protein-ligand complexes to predict relative binding affinities, with affinity labels derived from identical assays to minimize systematic errors common in large-scale affinity databases^12^. While PBCNet showed significant advantages over existing high-throughput methods and achieved comparable performance to Schrödinger’s FEP+ after fine-tuning, there remains a performance gap in zero-shot predictions.

We identified two primary limitations. First, PBCNet’s training dataset, though carefully curated, encompasses only ~20,000 unique small molecules and 960 protein targets, representing a limited chemical and biological space. Recent research on scaling laws^13,14^, validated across multiple domains including protein structure prediction^15^ and large language models^16^, indicates that model performance correlates positively with data volume and model complexity. The restricted scale of PBCNet’s training set may therefore limit its generalization across diverse chemical structures and protein families. Second, PBCNet’s geometric encoding approach introduces inherent constraints. The model utilizes atom pairwise statistical potentials^17^ (APSP) and angular encoding modules to guide attention weights during message passing. While these components implement physical principles such as distance-dependent interactions and directional bonding, they rely on predetermined geometric features from known atom pairs and fixed angle bins. This methodology potentially limits the model’s ability to learn context-specific interactions from structural data and may be suboptimal for novel atom types not included in the predefined APSP set^18^. These geometric constraints could impede the detection of unusual binding patterns, particularly in systems with non-standard ligand chemistry^19^.

To address these limitations and advance the field of protein-ligand binding affinity prediction, we present PBCNet2.0, a next-generation model leveraging equivariant graph neural networks (E-GNNs)^20^. Our approach is guided by two key insights from recent advances in artificial intelligence. First, inspired by scaling laws, we expanded our training dataset to encompass 8.6 million protein-ligand complex pairs from BindingDB^21^, representing a 14-fold increase over PBCNet’s original dataset. This expanded dataset covers a broader chemical and biological landscape, with 0.28 million unique small molecules and 1,122 distinct protein targets, enabling more comprehensive learning of diverse interaction patterns. Second, we fundamentally redesigned our model architecture by implementing a Cartesian-based E-GNN^22^ framework for message passing. This approach naturally encodes three-dimensional structural relationships through equivariant transformations, moving beyond PBCNet’s reliance on predefined potential functions to learn geometric constraints directly from data. This study presents our investigation into these innovations through several key aspects:

First, we evaluate PBCNet2.0’s performance in lead optimization tasks, comparing it with existing computational methods including physics-based approaches and AI models. We then analyze the model’s ability to capture and interpret protein-ligand interactions through detailed ablation studies and geometric perturbation experiments. The discussion extends to an unexpected emergent capability: the prediction of binding affinity changes induced by pocket mutations, which has notable implications for understanding drug resistance^23–26^. Finally, we validate our findings through prospective experiments on two drug targets, ectonucleotide pyrophosphatase/phosphodiesterase 1 (ENPP1)^27^ and aldehyde dehydrogenase 1B1 (ALDH1B1)^28^, demonstrating the practical utility of our approach in real-world molecular probe development and lead optimization scenarios.

## 2. Results

### 2.1 PBCNet2.0: a structure-based ΔpAct predictor

Building upon the Siamese Neural Network (SNN) architecture of PBCNet, PBCNet2.0 introduces an equivariant message passing module to analyze protein-ligand binding interfaces and predict relative binding affinity changes. Here, we describe the model’s training methodology, architectural design, and intended applications. Throughout our analysis, we use ΔpAct to denote the negative logarithm of relative binding affinity, applicable across multiple experimental measurements (IC_50_, EC_50_, K_i_, and K_d_).

Recent studies of scaling laws in machine learning have demonstrated that model performance consistently improves with increased training data volume and model complexity, providing empirical guidance for AI system optimization. Based on this principle, we assembled a comprehensive training dataset of 8.6 million protein-ligand complex pairs (Fig. 1a), a 14-fold increase from PBCNet’s original dataset (0.6 million). The ***Training set*** section details the data processing methodology.

**Figure 1.**
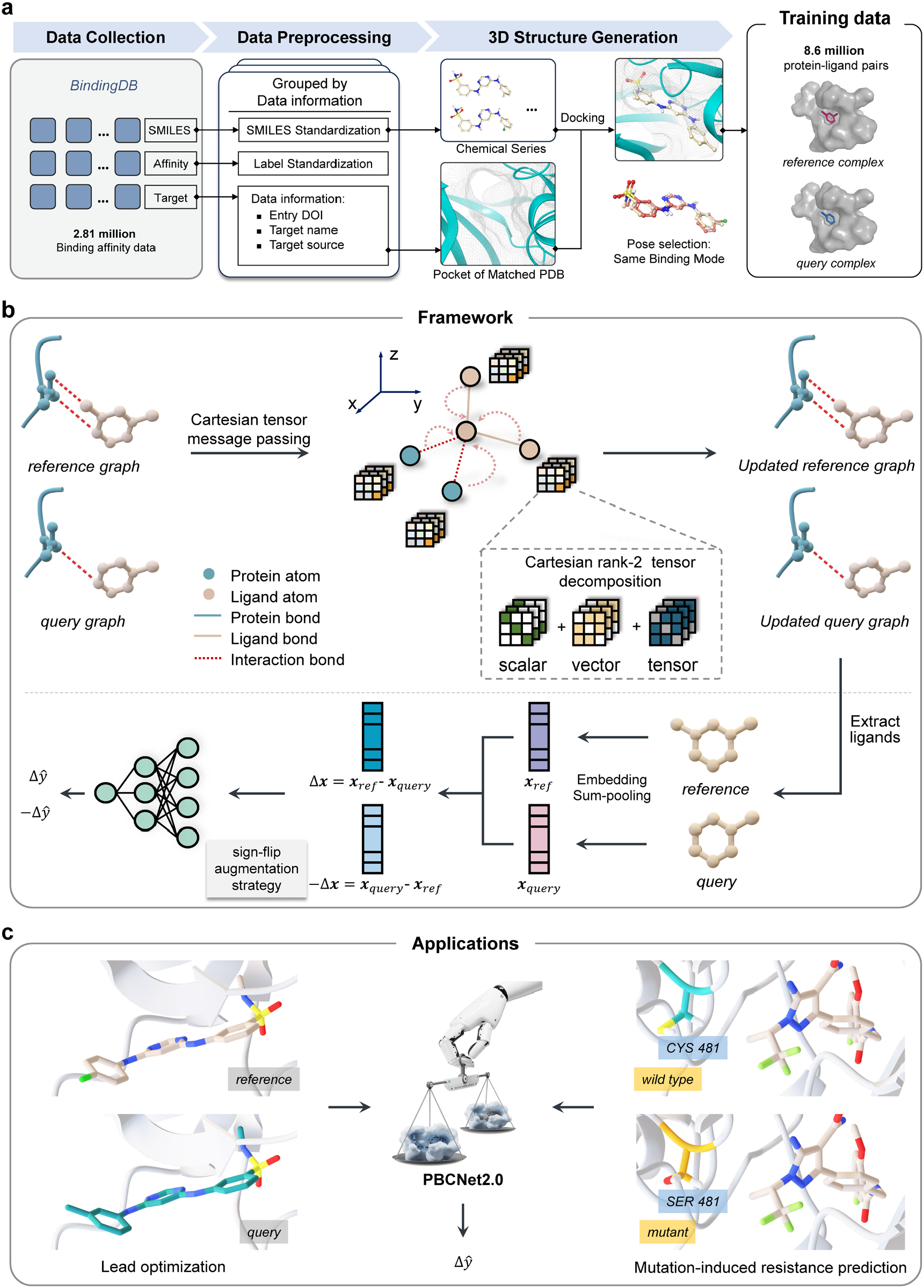
a. Processing of training data for PBCNet2.0. The raw data was collected from BindingDB (2023.12 version), and input binding poses were generated through docking and rule-based filtering. PBCNet2.0 is trained on 8.6 million protein-ligand pairs. The training pairs share the same protein pocket but differ in ligand structures. **b. The framework of PBCNet2.0**. The complexes are processed through a Cartesian rank-2 Tensor-based message-passing module, designed to facilitate mutual information exchange between the ligands and the protein pocket. This module yields node-level representations. In this process, each hidden dimension of each node-level representation is represented as a 3 × 3 matrix, which can be decomposed into three matrices with degrees of freedom 1, 3, and 5, respectively, using the irreducible representation of tensors. This decomposition enables efficient equivariant information propagation through matrix multiplication. In the Readout module, molecular representations (graph-level) are obtained by averaging the scalar node-level representations of the ligand atoms. The difference between the representations of the reference and query molecules (***x***^*ref*^ and ***x***^*que*^) serves as the final pair-wise representation (Δ***x***), which is then fed into the Prediction module to output the predicted relative binding affinity Δ*y*. A computationally efficient sign-flip augmentation strategy is also implemented here. **c. The application of PBCNet2.0**. During inference, the PBCNet2.0 framework requires only the binding poses of two structurally similar small molecules bound to the same protein to predict their relative binding affinity. Additionally, when given structures with identical ligand molecules bound to protein pockets containing specific residue mutations, the model can predict the resulting changes in binding affinity.

The PBCNet2.0 architecture consists of three primary components: (1) a message-passing module, (2) a readout module, and (3) a prediction module, as illustrated in Figure 1b. The message-passing module facilitates information transfer across protein-ligand interaction graphs to generate node-level representations. The key advancement in PBCNet2.0 lies in the integration of an equivariant graph neural network^20^ with Cartesian tensors^22^, enabling comprehensive analysis of distance-dependent and angular geometric information in protein-ligand interactions and molecular conformations.

In the implementation, each node’s hidden representation is encoded as a Cartesian rank-2 tensor, represented by a 3 × 3 matrix. This matrix is decomposed into three independent 3 × 3 matrices with 1, 3, and 5 degrees of freedom (detailed in Equation 1, ***Methods***), corresponding to scalar, vector, and tensor features. These features undergo parallel updates through 3 × 3 matrix operations, maintaining model equivariance efficiently. During message passing, protein pocket and ligand atom representations are updated through their interactions, after which pocket atoms are removed while ligand atoms are retained.

The readout module generates graph- and pair-level representations. Graph-level representations (***x***_*ref*_ and ***x***_*query*_) are computed by averaging the invariant node representations of reference and query ligands. The pair-level representation is defined as Δ***x*** = ***x***_*ref*_ − ***x***_*query*_, which differs from PBCNet’s concatenation of ***x***_*ref*_, ***x***_*query*_, and Δ***x***. This modification enhances the model’s focus on protein-ligand interactions, as structural similarity between analogous ligands would otherwise result in Δ***x*** approaching zero, creating computational challenges. The pair-level representation is processed through a three-layer feedforward neural network in the prediction module to estimate relative binding affinity (Δ *ŷ*). To ensure anti-symmetry (*f*(Δ***x***) = −*f*(−Δ***x***)), we incorporate −Δ***x*** = ***x***_*query*_ − ***x***_*ref*_ during training and compute the combined loss. Further technical details are provided in the ***Methods*** section.

For inference, PBCNet2.0 requires only the binding poses of two structurally similar molecules bound to the same protein target to predict their relative binding affinity (Fig. 1c). This enables systematic ranking of compound series based on predicted binding affinities, facilitating rational lead optimization and probe development. Notably, PBCNet2.0 exhibits an additional capability: when presented with structures of an identical ligand bound to protein pockets containing specific amino acid mutations, the model can predict the resulting changes in binding affinity (Fig. 1c). This capability will be thoroughly examined in subsequent sections.

### 2.2 PBCNet2.0 Achieves Substantial Improvement in Lead Optimization Tasks with Perception of Protein - Ligand Interaction Information

Following our previous work on PBCNet^6^, we first systematically evaluated the performance of PBCNet2.0 in lead optimization tasks through three experimental paradigms: (1) **Zero-shot prediction**, which assess guidance capability during early-stage optimization when no structure-activity relationship (SAR) data is available; (2) **Few-shot prediction**, which evaluates performance in mid-to-late-stage optimization where a modest amount of SAR data enables model fine-tuning; and (3) **Prioritization experiment**, which validates the model’s ability to identify high-activity compounds in real-world lead optimization scenarios. All metrics and methodologies matched our previous work^6^, as summarized them in the Supplementary Section 1.

The zero-shot prediction results, presented in Figure 2a and Supplementary Figure 1, show the average performance of various methods across 16 chemical series referred to as the FEP test set (combining FEP1 and FEP2 sets). We assessed ranking performance using Pearson’s correlation coefficient (R) and Spearman’s rank correlation coefficient (ρ) (refer to ***Evaluation metrics*** section). We compared against twelve baseline methods (detailed in ***Baseline models*** section): a deep learning based co-folding model (Boltz-2^29^), three traditional structure-based approaches (Glide SP^30^, MM-GB/SA^31^, and Schrödinger’s FEP+^8^), three sequence-based AI models (PSICHIC^7^, BIND^32^, and PLAPT^33^), and five structure-based AI models (PIGNet2^11^, RTMScore^34^, GenScore^35^, OnionNet-2^36^, and PBCNet^6^). The detailed prediction results of each method on each chemical series are summarized in the Supplementary Data 1.

**Figure 2.**
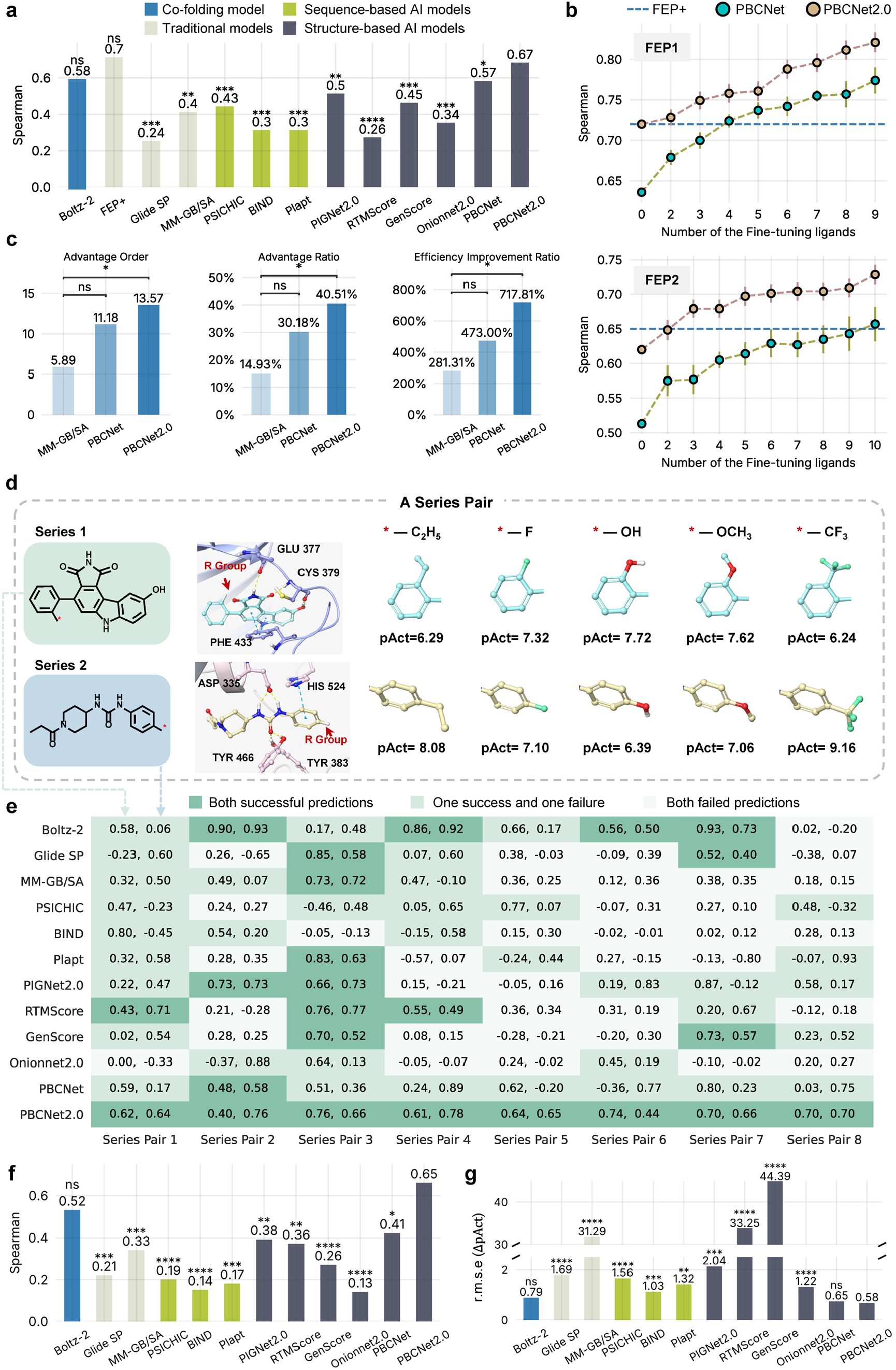
a. Direct prediction results. The x-axis represents various binding affinity prediction methods, while the y-axis denotes their average of their ranking capability (Spearman) on the FEP set. A two-sided Wilcoxon signed-rank test compared each model against PBCNet2.0, with significant levels: *p ≤ 0.05, **p ≤ 0.01, ***p ≤ 0.001, ****p ≤ 0.0001, and ns (not significant) for p > 0.05. Different model types are shown in different colors. **b. Finetuned prediction results**. The x-axis represents the number of compounds used for fine-tuning, while the y-axis shows the ranking capability metrics. Cyan dots represent PBCNet performance, pink dots show PBCNet2.0 performance, and a blue dashed line indicates the Schrödinger’s FEP+ baseline. **c. Selection experiment results**. The x-axis represents different methods, and the y-axis shows performance across eight systems. Three panels display distinct evaluation metrics. A one-sided Wilcoxon signed-rank test evaluated whether PBCNet2.0 and PBCNet significantly outperformed MM-GB/SA. **d. Illustration of the partial modification process for a chemical series pair from the SAR-Diff Set**. A schematic representation of structural changes within a chemical series pair. **e. The predictive performance of the models on the SAR-Diff set**. The x-axis lists 8 series pairs, and the y-axis represents 11 prediction methods. Each cell in the heatmap displays the Spearman correlation coefficient for a model on a given series pair. Dark green cells indicate models successfully predicted both series within a pair, light green cells show models with predictive ability for only one series, and pale cells represent models without predictive ability for either series. **f-g. Average performance of methods on SAR-Diff set**. The x-axis represents various binding affinity prediction methods, while the y-axis denotes the average of their ranking capability (Spearman) and prediction accuracy (r.m.s.e with ΔpAct) on the SAR-Diff set. The one-sided Wilcoxon signed-rank test was used to analyze whether the ranking performance of PBCNet2.0 was significantly superior to others and whether its prediction error was significantly lower. *p ⩽ 0.05, **p ⩽ 0.01, ***p ⩽ 0.001, ****p ⩽ 0.0001, and ns (not significant) for p > 0.5. Different types of models are distinguished using bars of different colors.

Benchmarking results establish PBCNet2.0’s superior predictive capability, achieving ρ = 0.67 and R = 0.66 on the FEP set, an 18% enhancement over its predecessor PBCNet (ρ = 0.57, R = 0.56). Notably, PBCNet2.0 approaches the performance of the gold-standard Schrödinger FEP+ (ρ = 0.70 and R = 0.70, Δρ = 0.03 and ΔR = 0.04), with statistically indistinguishable ranking accuracy (p = 0.39 > 0.05). This result makes PBCNet2.0 the first AI approach to achieve parity with industrial-strength FEP simulations in critical evaluation metrics.

To visualize PBCNet2.0’s improvements, we compared predicted ΔpAct values from both PBCNet2.0 and PBCNet for test systems from the FEP set (Supplementary Fig. 2). PBCNet2.0’s predictions exhibit a stronger correlation with experimental values and cover a broader value range. For example, in the pfkfb3 system, PBCNet2.0’s predictions span [−2, 2], closely matching the experimental value range, while PBCNet’s predictions remain limited to [−1, 1].

Figure 2b illustrates the few-shot prediction results, with detailed data summarized in Supplementary Data 2. The x-axis indicates the number of compounds used for fine-tuning, while the y-axis shows the ranking capability metrics. Cyan dots represent PBCNet’s performance, red dots show PBCNet2.0’s performance, and a blue dashed line indicates the Schrödinger’s FEP+ baseline. Fine-tuning proves highly effective for PBCNet2.0: its performance improves steadily as more fine-tuning ligands are added, consistently outperforming both PBCNet and Schrödinger’s FEP+. This data-efficient refinement capability makes PBCNet2.0 valuable when limited SAR data is available for a target of interest.

The real-world utility of PBCNet2.0 is demonstrated through retrospective prioritization experiments across 8 pharmaceutically relevant targets (Fig. 2c). The detailed data and evaluation process are summarized in Supplementary Data 3 and the Supplementary Section 1. Benchmarking results show that PBCNet2.0 outperforms both PBCNet and MM-GB/SA on all metrics. Specifically, PBCNet2.0 achieved an **advantage order** of 13.57, while compared to PBCNet’s 11.18 and MM-GB/SA’s 5.89. The **advantage ratio**, which measures theoretical resource savings in model-guided lead optimization, shows PBCNet2.0 reduced resource investment by 40.51%, surpassing PBCNet (30.18%) and MM-GB/SA (14.93%). The **efficiency improvement ratio**, reflecting enhanced efficiency in compound optimization, shows PBCNet2.0 accelerated lead-optimization projects by 717.81%, surpassing both PBCNet (473.00%) and MM-GB/SA (281.31%). Statistical analysis confirms that the performance of PBCNet2.0 in the prioritization experiment is significantly (p ≤ 0.05) superior to MM-GB/SA.

These comprehensive results demonstrate that PBCNet2.0 exhibits significant advances in relative binding affinity prediction, with notable improvements in both accuracy and reliability. This enhanced predictive capability positions PBCNet2.0 as a valuable tool for molecular probe development and lead optimization. We next examine the underlying mechanisms contributing to its predictive performance.

Mastropietro et al. reported that many machine learning-based scoring models for protein-ligand binding suffer from a bias where predictions are predominantly driven by ligand memorization rather than understanding protein-ligand interactions^37^. To rigorously test whether PBCNet2.0 overcomes this bias. First, we conducted a strategic ablation study using the FEP dataset. We systematically removed all protein-ligand interaction edges from the original molecular graphs (**Complete graphs**), generating the **Incomplete graphs** (Supplementary Fig. 3a). We then monitored the performance dynamics of PBCNet2.0 on both graphs over the course of training (Supplementary Fig. 3b). During the initial phase of training (approximately up to 12.5K steps), both curves exhibit a similar upward trend, suggesting that the model initially mainly relies on molecular structure information for predictions. However, as training progresses beyond 12.5K steps, a marked divergence emerges. The model performance on the **Incomplete graphs** declines steadily, whereas that on the **Complete graphs** continues to improve. This pattern indicates that protein-ligand interaction information has emerged as the primary determinant of PBCNet2.0’s predictions, implying that PBCNet2.0 may progressively develop an understanding of the interaction information.

To further verify that PBCNet2.0 learns protein-ligand interaction patterns rather than memorizing molecular features or structural variations, we evaluated its performance on the SAR-Diff set, which was specifically designed to decouple structural modifications from binding affinity changes (refer to ***SAR-Diff held-out*** test set section). Specifically, the SAR-Diff set comprises eight pairs of chemical series (16 series total) where each pair share identical R-group modification but exhibit distinct structure-activity relationships^38–40^. As illustrated in Figure 2d with an example from the first series pair, and further demonstrated in Supplementary Figure 3c showing the absence of activity correlation between series, this dataset design necessitates models to identify interaction-dependent activity variations instead of relying on structural memorization. Therefore, successful predictions on this benchmark require accurate modeling of the interactions between modified chemical groups and their corresponding protein binding sites.

Figure 2e presents the comprehensive evaluation results (Supplementary Data 4), with heatmap organized to show performance of different methods (y-axis) across series pairs (x-axis). Spearman correlation coefficients (ρ) quantify prediction quality. The two numbers in each heatmap cell stand for the ρ of a model on a given series pair. Each column indicates the ρ of different models on the same chemical series, as illustrated by the arrows between Figure 2d and Figure 2e. When the ρ ≥ 0.4, the corresponding model is considered to have predictive power on the series. The darkest green cells denote that models successfully predicted both series within a pair, lighter green indicate models only own predictive ability for one series, and pale cells represent models have no predictive ability for either series. PBCNet2.0 exhibited consistent predictive capability across all series pairs, distinguishing itself from other methods, including Boltz-2 which succeeded in only four pairs. Notably, both PBCNet2.0 and PBCNet were unique in avoiding complete prediction failures for any series pair.

Figure 2f provides a detailed comparison of average model performance on these 16 series (ranking performance measured by ρ and prediction accuracy measured by root-mean-square error, r.m.s.e.). PBCNet2.0 demonstrated superior performance in both metrics. These results highlight that PBCNet2.0 successfully masters the interactions between modified groups and protein pockets, with protein-ligand interaction patterns serving as key determinants of prediction accuracy. A comparison between Figure 2f and 2a reveals that the SAR-Diff set presented a more challenging evaluation scenario than the FEP set for most models (with the exception of RTMScore), further validating PBCNet2.0’s robust performance under demanding conditions.

These collective results demonstrate that PBCNet2.0 serves as an effective computational tool for lead optimization and molecular probe development. The model shows measurable improvements over its predecessor PBCNet across multiple performance metrics, including ranking accuracy and prediction capability. The mechanistic analysis reveals that PBCNet2.0 effectively incorporates protein-ligand interaction information to enable accurate relative binding affinity predictions.

### 2.3 PBCNet2.0 Captures Subtle Intermolecular Interactions with Strict Geometric Constraints

Following our demonstration that protein-ligand interaction information plays a crucial role in PBCNet2.0’s predictions, we sought to evaluate the model’s capability in identifying specific protein-ligand interactions and its utilization of geometric information. While our previous analysis of PBCNet focused primarily on hydrogen bond (H-bond) interaction recognition^41^, we extended this investigation to PBCNet2.0. Through examination of the same cases analyzed in the PBCNet study, we found that PBCNet2.0 maintains comparable effectiveness in H-bond interaction identification, as evidenced in Supplementary Figure 4.

Beyond H-bonds, we investigated fluorine orthogonal multipolar interactions^42^, an important but frequently overlooked protein-ligand interaction type^43–45^. Traditional empirical expressions for intermolecular interactions, including many general-purpose force fields, often inadequately represent this interaction^43,46^. We investigated whether PBCNet2.0, utilizing a data-driven approach, could learn and identify this interaction from comprehensive structural data. As illustrated in Figure 3a, fluorine’s high electronegativity creates a nucleophilic center, while the carbonyl oxygen’s electron-withdrawing effect makes the carbonyl carbon electrophilic. This electronic complementarity enables orthogonal multipolar interactions between fluorine and carbonyl carbon. These interactions are characterized by specific geometric requirements: the fluorine-carbon distance must be under 3.8Å, with the F ⋯ C = O angle approaching 90°. Accurate detection of such interactions requires precise modeling of both 3D spatial relationships and angular dependencies.

**Figure 3.**
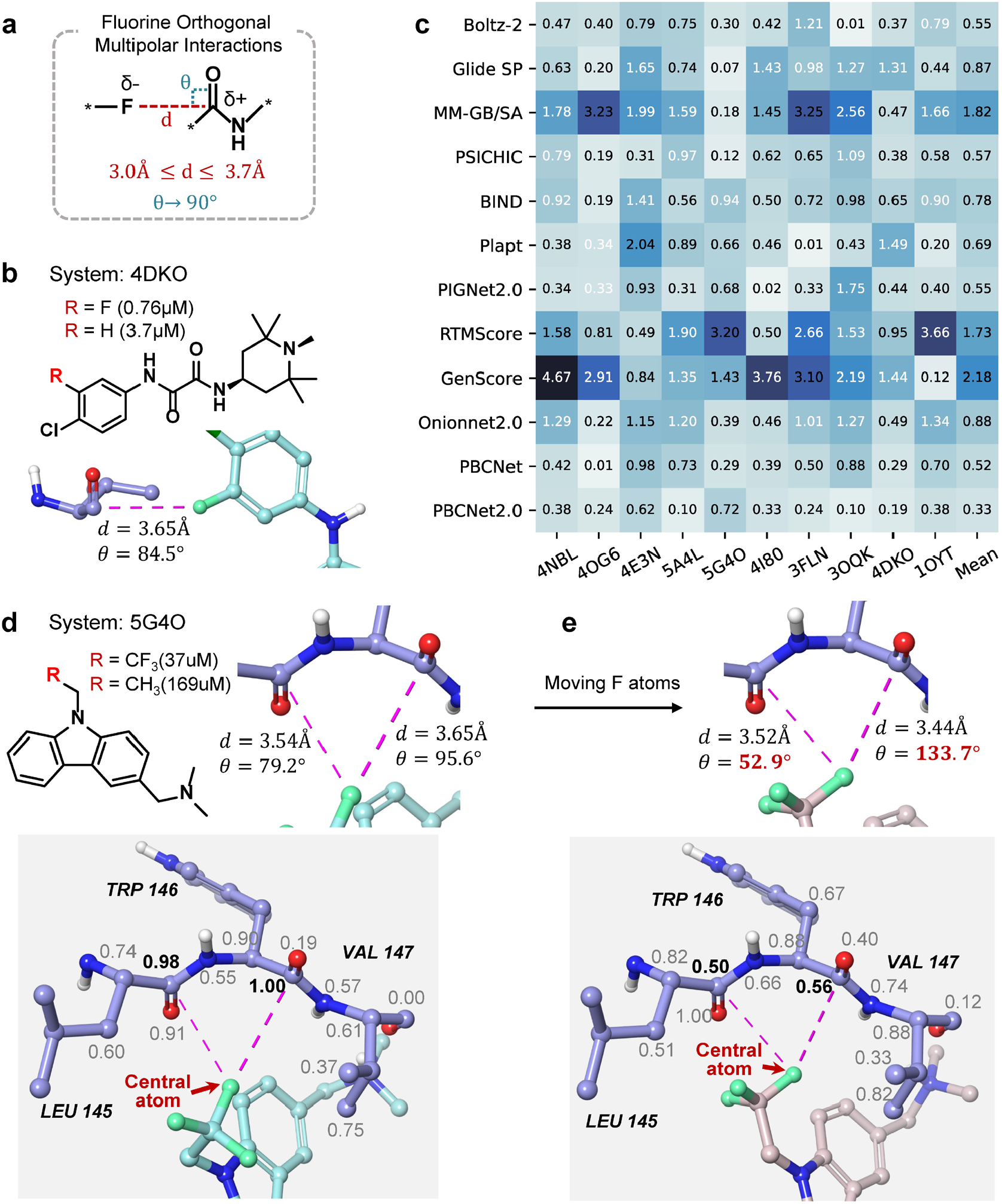
a. Illustration of fluorine orthogonal multipolar interaction. A schematic demonstrating its formation mechanism and geometric constraints. **b. Examples from the F-Opt Set**. Representative cases showcasing fluorine orthogonal multipolar interactions, including key distances and angles. **c. Absolute prediction errors of models on each compound pair:** The y-axis represents various methods, and the x-axis corresponds to compound pairs (denoted by PDB IDs). The numbers in the table indicate absolute errors, with white numbers highlighting incorrect predictions. **d. Interpretability analysis of the 5G4O system**. For a molecule with a trifluoromethyl R-group, two fluorine orthogonal multipolar interactions with the protein are identified. The binding mode and geometric constraints are illustrated, with fluorine orthogonal multipolar interactions represented by purple dashed lines. Protein atoms within 5 Å of the central fluorine atoms are annotated with weights, and bold numbers denote the weights of carbon atoms involved in the interactions. **e. Disruption of geometric constraints in fluorine orthogonal multipolar interaction**. The angles of fluorine orthogonal multipolar interactions were disrupted, changing one angle from 79.2° to 52.9° and another from 95.6° to 133.7° (both being unreasonable angles). Consequently, the weight distribution of surrounding atoms was altered.

To evaluate this capability systematically, we developed the F-Opt benchmark test set (see ***F-Opt test set*** section), comprising 10 selected compound pairs where fluorine substitution creates orthogonal multipolar interactions that enhance binding affinity (Fig. 3b, Supplementary Fig. 5). We assessed various binding affinity prediction methods using this benchmark. Figure 3c (Supplementary Data 5) displays the absolute prediction errors for each model across compound pairs (identified by PDB IDs), with darker shading indicating larger errors. White numbers denote cases where predicted and experimental ΔpAct values show opposite signs (Sign(Predicted ΔpAct) ≠ Sign(Experimenta*l* ΔpAct)). PBCNet2.0 demonstrated the highest accuracy with the lowest mean absolute error (m.a.e. = 0.33) among all methods. PBCNet achieved the second-best performance (m.a.e. = 0.52). Notably, only PBCNet2.0 and PBCNet achieved 100% sign concordance, while other methods showed error rates between 20% and 50%, highlighting the reliability of both approaches.

To gain mechanistic insights, we conducted interpretability analyses on two systems from the F-Opt benchmark (5G4O and 4E3N, detailed in ***Interpretability analysis*** section) to understand how fluorine orthogonal multipolar interactions are recognized. As shown in Figure 3d and Supplementary Figure 6, these interactions are represented by purple dashed lines. We annotated protein atoms within 5Å of the fluorine atoms (central atoms) with weight values (0.00 − 1.00), with boldface numbers indicating the interaction weights of participating carbon atoms. The consistently high weights (0.98 − 1.00) assigned to these carbon atoms in both systems demonstrate PBCNet2.0’s ability to identify geometrically constrained fluorine interactions. To evaluate PBCNet2.0’s sensitivity to angular parameters, we conducted a perturbation study on the 5G4O system. Using Maestro, we modified two key interaction angles by moving the central fluorine atom: reducing one angle from79.2° to 52.9° and increasing another from 95.6° to 133.7° (both angles falling outside the acceptable range). Throughout this process, the ligand pose was adjusted to maintain structural integrity. Figure 3e shows that the weight values for the interacting carbon atoms decreased substantially (from 0.98 to 0.50 and from 1.00 to 0.56), demonstrating that the equivariant framework enables PBCNet2.0 to recognize angular constraints. This geometric sensitivity is essential for accurately modeling subtle intermolecular interactions.

Despite PBCNet’s strong predictive performance on the F-Opt test set, its interpretability analysis reveals limitations in mechanistic understanding (Supplementary Fig. 7a). Analysis of PBCNet’s statistical atom-pair potential function shows that this empirical approach assigns unexpectedly low interaction probabilities to fluorine-carbonyl carbon pairs at relevant distances (Supplementary Fig. 7c), indicating inadequate modeling of fluorine orthogonal multipolar interactions. Additionally, PBCNet shows limited sensitivity to angular parameters (Supplementary Fig. 7b). These limitations stem from its relatively coarse 3D modeling approach, which constrains its ability to capture precise physicochemical interactions. In comparison, PBCNet2.0’s geometry-aware equivariant architecture, combined with extensive training data, enables more sophisticated modeling of intermolecular interactions. This architectural advancement not only improves predictive performance but also establishes a robust framework for learning complex interaction patterns through data-driven approaches. The clear correlation between predictions and underlying physicochemical principles enhances PBCNet2.0’s utility for both probe development and lead optimization.

### 2.4 PBCNet2.0 Demonstrates Emergent Abilities for Identifying Drug-Resistant Mutations

Residue mutations can directly impact protein structure, stability, function, and molecular interactions^24–26^ (Fig. 4a). Residue mutations may alter binding site properties and drug binding affinities, potentially leading to drug resistance^23^. For instance, in non-small cell lung cancer (NSCLC) treated with first-generation EGFR inhibitors (e.g., Gefitinib^47^), the acquired T790M mutation in EGFR kinase domain emerges in > 50% of resistant cases within 9-14 months^48,49^, sterically hindering drug binding and causing therapeutic failure. This rapid resistance crisis necessitated urgent development of next-generation inhibitors, exemplified by Osimertinib^50^—a third-generation EGFR inhibitor specifically engineered to overcome T790M-mediated resistance. Critically, the stronger the drug selection pressure, the more rapidly resistance develops because drug target has been proved to be very plastic^51^. Hence, an early prediction and identification of mutations that may potentially impact ligand binding is important for lead optimization and probe development. Although PBCNet2.0 was not explicitly trained on data involving amino acid mutations (the training set only contained data for different ligands binding to the same protein site), we were curious whether PBCNet2.0 could have automatically learned to predict mutation-induced binding affinity changes. This hypothesis was motivated by the shared nature of both tasks— modeling changes in protein-ligand interactions—and by PBCNet2.0’s ability to capture subtle molecular interactions as demonstrated above. To rigorously test this hypothesis, we constructed a Mutation Benchmark Set (see ***Mutation test set*** section) containing 8 pharmaceutically relevant targets with clinically observed binding pocket mutations. As shown in Figure 4b (Supplementary Data 6), PBCNet2.0 demonstrated relative solid predictive performance, achieving an average Spearman correlation of 0.53 (*ρ* = 0.53). This is a stark contrast to methods like Boltz-2, Glide SP, BIND, PLAPT, RTMScore, and GenScore, which even exhibited negative correlations. The failure of Boltz-2 on this task may be attributable to the co-folding method’s difficulty in accurately predicting the structural changes caused by mutations in the amino acid sequence, thereby causing its affinity prediction module to completely fail.^52^ A deeper target-level look into PBCNet2.0’s predictions exposed varying performances across different targets. While PBCNet2.0 exhibited strong ranking performance for tACE^53^ (*ρ* = 0.76) and HIV-1 PR^54^ (*ρ* = 0.75) targets (Supplementary Fig. 8a), it showed less satisfactory results for AKR1B1^55^ (*ρ* = 0.28) and PfDFHR-TS^56^ (*ρ* = 0.26) targets (Supplementary Fig. 8b).

**Figure 4.**
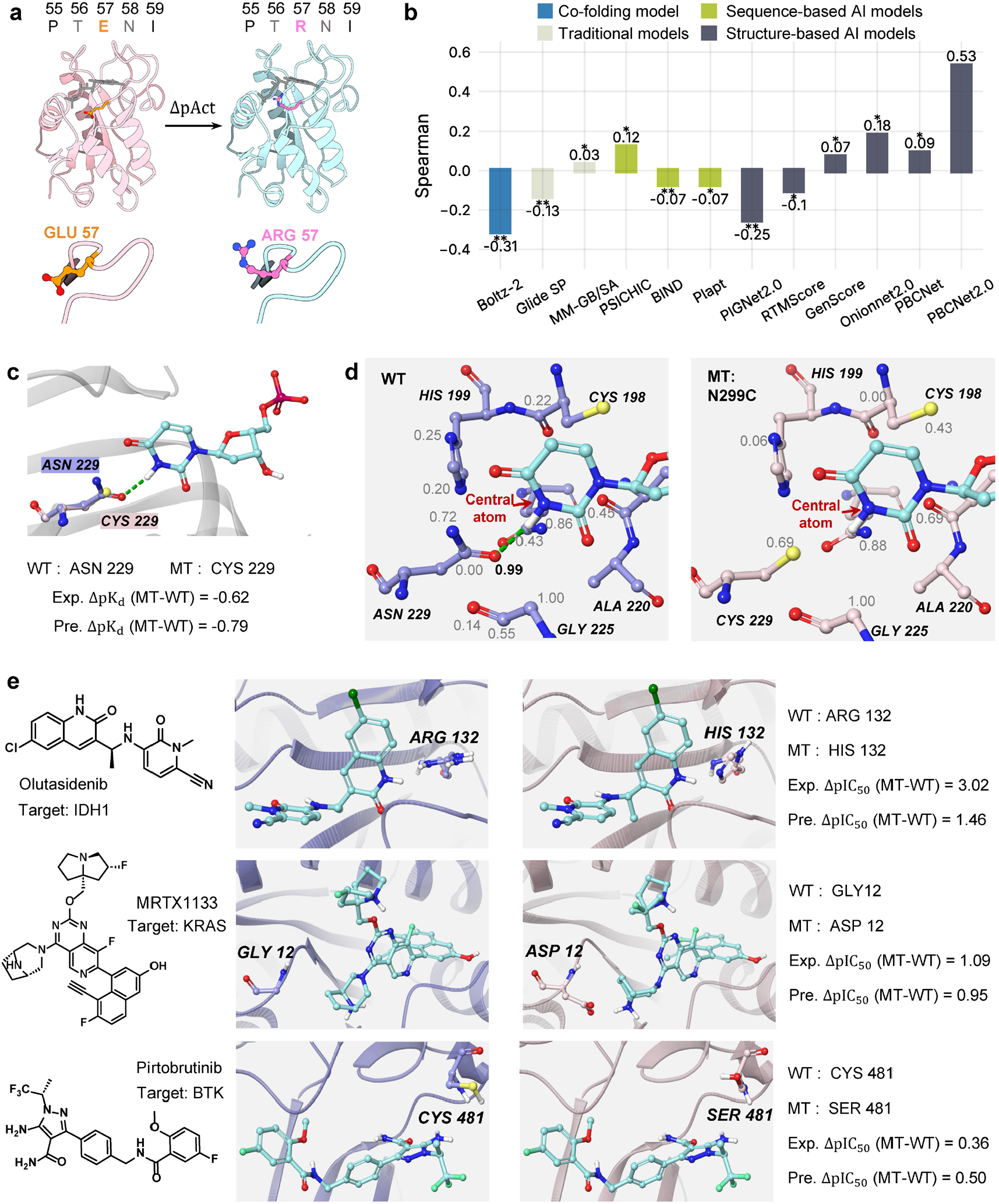
a. Illustration of amino acid mutation. A schematic representation of a mutation at position 113 in the protein pocket, where THR is mutated to TYR. This alters the protein pocket environment, affecting small molecule binding and resulting in changes to binding affinity. **b. Average performance of methods on the Mutation set**. The x-axis represents various binding affinity prediction methods, while the y-axis denotes the average of their ranking capability (Spearmen) on the Mutation set. The one-sided Wilcoxon signed-rank test was used to analyze whether the ranking ability of PBCNet2.0 was significantly superior to others. *p ≤ 0.05 and **p ≤ 0.01. Different types of models are distinguished using bars of different colors. **c. Binding Mode of the Mutation Case**. The mutation of ASN at position 229 to CYS causes the protein to lose a hydrogen bond with the small molecule, leading to a decrease in pK_d_ by 0.62. PBCNet2.0 predicts a decrease of 0.79. **d. Interpretability analysis**. The hydrogen bond is depicted as green dashed lines, and protein atoms within 5 Å of the central nitrogen atom were annotated with weights, where the bold number indicates the weights associated with the oxygen atom involved in the hydrogen bond. **e. Retrospective analysis of mutant-selective inhibitors**. Three mutant-selective inhibitors that are either in clinical trials or already in the market are shown: Olutasidenib (target: IDH1), MRTX1133 (target: KRAS), and Pirtobrutinib (target: BTK). For each compound, the figure shows chemical structures on the left, binding modes in wild-type and mutant proteins in the middle panels, and quantitative data on the right. The right section presents wild-type and mutant residues alongside experimental and predicted ΔpIC_50_ values for each protein-ligand pair.

To further investigate the rationale behind PBCNet2.0’s predictions, we conducted an interpretability analysis of **Lactobacillus casei Thymidylate Synthase** (LTS)^57^ target. The N229C mutation abolished a key hydrogen bond between the ASN229 amide oxygen and ligand pyridine nitrogen (Fig. 4c). This mutation decreased the binding affinity (ΔpK_d_ = −0.62)^57^, closely matching PBCNet2.0’s prediction (ΔpK_d_ = −0.79). Additionally, we visualized the weights (details in ***Interpretability analysis*** section) assigned to neighboring atoms of the pyridine nitrogen during the model’s message-passing phase (Fig. 4d). In Figure 4d, the hydrogen bond is depicted as green dashed lines, and protein atoms within 5 Å of the central nitrogen atom were annotated with weights, where boldface numeral indicates the weight of the oxygen atom involved in the interaction. We found that the involved oxygen atom was assigned a relative high weight (0.99) in wild-type simulations, with complete signal loss post-mutation. This phenomenon demonstrates the validity of PBCNet2.0’s predictions. Through its acquired ability to capture protein-ligand interactions, the model has successfully generalized to mutation prediction tasks.

Thereafter, we retrospectively analyzed three cases of mutant-selective inhibitors that are either in clinical trials or already in the market: (1) Olutasidenib, approved by the Food and Drug Administration (FDA) in December 2022, targets the mutant **Isocitrate Dehydrogenase 1** (IDH1) (with >1000-fold selectivity) to treat acute myeloid leukemia (AML) ^58^; (2) MRTX1133, a potent, noncovalent, and selective **Kirsten Rats Arcomaviral Oncogene Homolog** (KRAS) G12D inhibitor, entered clinical trials in March 2023 to address manifold tumors harboring KRAS^G12D^ mutations^59^; and (3) Pirtobrutinib, the first-in-class noncovalent **Bruton’s tyrosine kinase** (BTK) inhibitor approved by the FDA^60^. The structures, binding modes, and experimental and predicted relative binding affinities of these inhibitors are depicted in Figure 4e. As shown in Figure 4e, PBCNet2.0 accurately predicted relative binding affinities across these structurally diverse inhibitors, despite never being explicitly trained on mutation-specific data. This emergent capability^61,62^ likely stems from the model’s generalized understanding of protein-ligand recognition patterns, developed through training on a large-scale dataset containing 8.6 million entries.

The above experimental results demonstrate that PBCNet2.0 can identify critical mutations, positioning it as a versatile tool for both probe development and molecular optimization.

### 2.5 Experimental Validation

Building upon our retrospective evaluation and analysis of PBCNet2.0, we operationalized it in a prospective experimental validation study to verify its capacity to capture subtle intermolecular interactions and predict resistance-associated mutations that govern ligand binding.

#### 2.5.1 Experimental Validation of Subtle Molecular Interaction Capture in ENPP1 Inhibitors

In our previous work, we identified compound **A1** (Fig. 5a) as a promising ENPP1 inhibitor^27^, and the binding mode analysis suggested it might form a fluorine orthogonal multipolar interaction with the target protein. Here, we further investigate **A1** with detailed analysis. First, we conducted an interpretability analysis on the fluorine of **A1** (Fig. 5b). Results showed a relative high weight (0.94) for the potential dipole interaction between this fluorine and the target protein’s carbonyl oxygen. Then, based on **A1**, we designed an analog **A2** (Fig. 5a), replacing the fluorine with hydrogen to eliminate the potential dipole effect while minimizing other structural changes. Using PBCNet2.0, we predicted **A2**’s binding affinity for ENPP1 with **A1** as the reference. PBCNet2.0 predicted a substantial difference in binding affinity between **A1** and **A2**, specifically forecasting a 40-fold (ΔpAct = 1.61) reduction in binding affinity upon fluorine-to-hydrogen substitution. This marked disparity in predicted binding affinities provides strong computational evidence supporting the existence of the fluorine orthogonal multipolar interaction.

**Figure 5.**
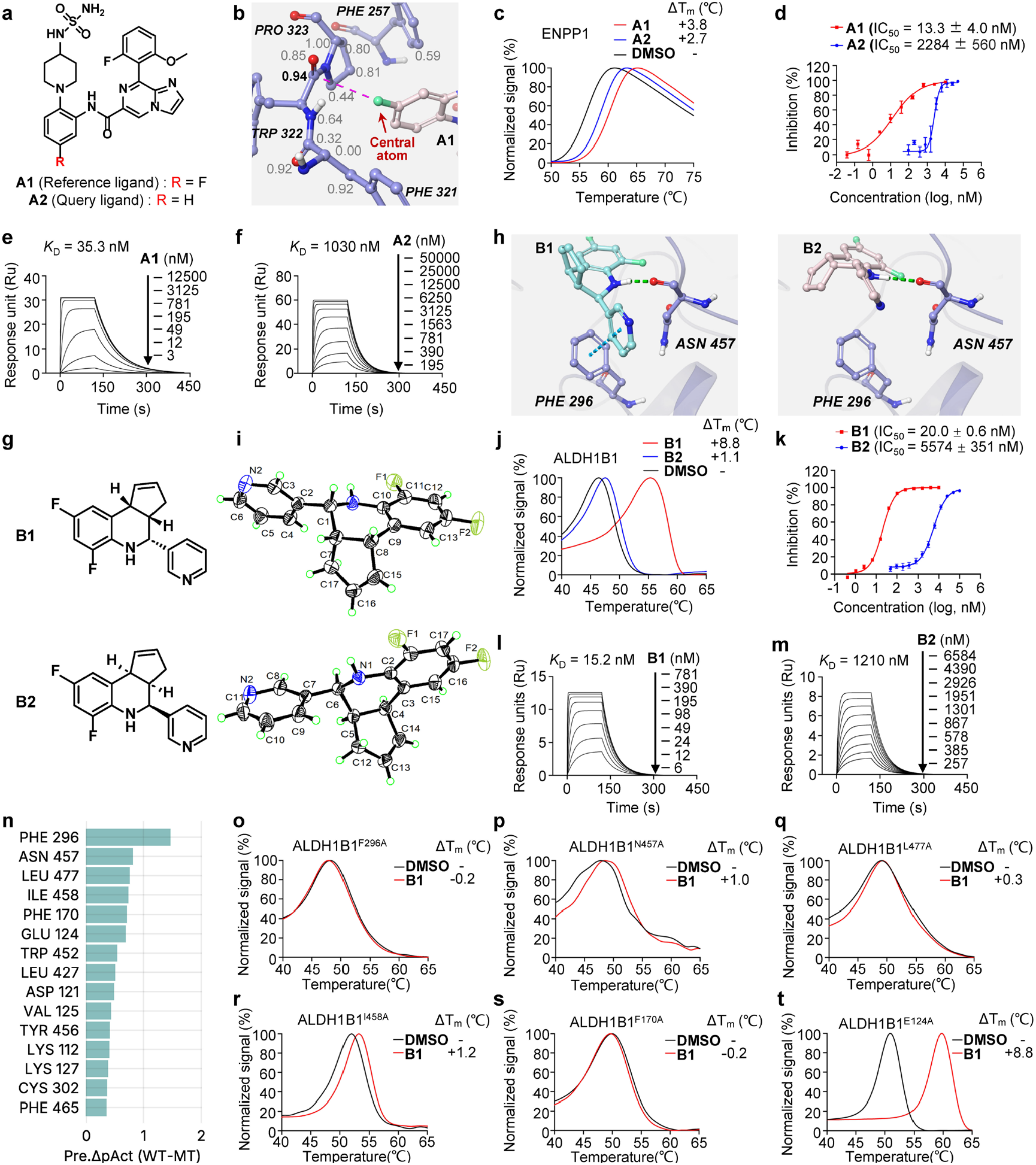
a. Chemical structures of ENPP1 inhibitors. Reference ligand **A1** (R = F) and query ligand **A2** (R = H) where the fluorine atom was replaced with hydrogen. **b**. Interpretability analysis. The fluorine orthogonal multipolar interaction between **A1** and TRP322 of ENPP1, with interaction distance (d = 3.22Å) and angle (θ = 109.2°). Fluorine orthogonal multipolar interaction represented by purple dashed lines. Protein atoms within 5 Å of the central fluorine atoms are annotated with weights, and bold numbers denote the weights of carbon atoms involved in the interactions. **c. Protein thermal shift assay**. Impact of **A1** (50 μM) and **A2** (50 μM) on the thermal stability of ENPP1 protein (2 μM) as determined by PTS assay. Temperature (°C) on the x-axis and normalized signal (%) on the y-axis. **d. *In vitro* enzyme activity assay**. Dose-dependent inhibition of **A1** and **A2** against ENPP1. The substrate for the ENPP1 enzymatic reaction is thymidine 5’-monophosphate p-nitrophenyl ester (TMP). Error bars represent the mean ± SEM of three independent experiments. **e-f. Surface plasmon resonance binding assay**. The binding kinetics measurement of **A1** and **A2** to ENPP1 protein were using SPR assay. Graphs depicting equilibrium response units versus **A1** and **A2** concentrations were plotted. **g. Chemical structures of ALDH1B1 inhibitors**. Chiral centers of Reference ligand **B1** and query ligand **B2** were different. **h. Binding modes of B1 and B2**. While **B1** and **B2** adopt largely similar binding modes, a critical difference emerged due to their distinct stereochemistry: the pyridine ring of **B2** is unable to form the favorable π − π stacking interaction with PHE296 that is observed for **B1. i. X-Ray Structural Data**. Absolute configurations for **B1** and **B2. j. Protein thermal shift assay**. Impact of **B1** (50μM) and **B2** (50μM) on the thermal stability of ALDH1B1 protein (5μM) Temperature (°C) on the x-axis and normalized signal (%) on the y-axis. **k. *In vitro* enzyme activity assay**. Dose-dependent inhibition of **B1** and **B2** against ALDH1B1. Error bars represent the mean ± SEM of three independent experiments. **l-m. Surface plasmon resonance binding assay**. The binding kinetics measurement of **B1** and **B2** to ALDH1B1 protein were measured using SPR assay. Graphs depicting equilibrium response units versus **B1** and **B2** concentrations were plotted. **n. Mutation prediction results of PBCNet2.0**. The x-axis represents the value of the decrease in binding affinity predicted by the model after the mutation, and the y-axis represents the mutated residue. The 15residues prioritized by the model (out of 43) are shown. **o-t. Protein thermal shift assay**. Impact of **B1** on the thermal stability of various ALDH1B1 mutants.

To validate the accuracy of these computational results, we synthesized **A1** and **A2** (Supplementary Section 3) and conducted corresponding experimental validations (Supplementary Section 2). The protein thermal shift (PTS) assay revealed that both **A1** and **A2** increased the thermal stability of ENPP1 protein (Fig. 5c), suggesting direct binding of these compounds to ENPP1. As expected, **A1** indeed exhibited a stronger thermostabilizing effect on ENPP1 (ΔT_m_ = 3.8°C) than **A2** (ΔT_m_ = 2.7°C). *In vitro* enzyme activity assays showed that **A1** inhibited ENPP1 enzymatic activity (IC_50_ = 13.3nM, Fig. 5d) obviously more potently than **A2** (IC_50_ = 2284nM, Fig. 5d). These data suggested that **A1** possessed stronger binding affinity for ENPP1 protein. This was confirmed by subsequent surface plasmon resonance (SPR) studies (Fig. 5e-f), which showed *K*_D_ values of 35.3nM and 1030nM for **A1** and **A2** binding to ENPP1, respectively, indicating a substantially higher binding affinity of **A1**. The Δp*K*_D_ measured by SPR assay was 1.46, closely aligning with PBCNet2.0’s prediction of 1.61, with a minor error of 0.15. These experimental observations collectively demonstrate the establishment of a robust fluorine-mediated orthogonal multipolar interaction between compound **A1** and ENPP1.

This analysis provides compelling validation of PBCNet2.0’s ability to accurately predict the impact of subtle intermolecular interactions on protein-ligand binding affinity. The strong correlation between computational predictions and experimental measurements supports the utility of PBCNet2.0 as a reliable tool for rational structure-based probe development and lead optimization.

#### 2.5.2 Experimental Validation of Subtle Molecular Interaction Capture and Drug-Resistant Mutation Prediction in ALDH1B1 Inhibitors

In our previous investigation, we identified a racemic compound exhibiting inhibitory activity against ALDH1B1. Two critical questions remained to be addressed: (1) The potential differential biological activities between stereoisomers, and (2) The identification of key amino acid residues that could potentially confer drug resistance. Here, we employed PBCNet2.0 to systematically investigate these aspects.

To investigate the first question, we performed computational analysis of the enantiomeric pair (**B1** and **B2**, Fig. 5g). Molecular docking revealed that despite overall similar binding modes, a key distinction arose from their stereochemical differences: **B2**’s pyridine ring could not establish the favorable π − π stacking interaction^63,64^ with PHE296 that was observed with **B1** (Fig. 5h). PBCNet2.0 predicted a significant difference in binding affinity, suggesting that **B1** would exhibit approximately 10-fold higher potency compared to **B2** (predicted ΔpAct = 1.07). Model interpretability analysis supported this docking observation by highlighting the significance of the π − π stacking interaction between **B1**’s pyridine ring and PHE296 (Supplementary Fig. 9).

To validate these computational predictions experimentally, we synthesized the racemate and separated the enantiomers **B1** and **B2** through chiral chromatography. Their absolute configurations were determined by X-ray crystallography (Fig. 5i, Supplementary Section 4). PTS assays (Fig. 5j) demonstrated that both compounds stabilized ALDH1B1, with **B1** showing notably higher thermostabilization (ΔT_m_ = 8.8°C) compared to **B2** (ΔT_m_ = 1.1°C). Enzymatic assays (Fig. 5k) revealed that **B1** inhibited ALDH1B1 (IC_50_ = 20nM) substantially more potently than **B2** (IC_50_ = 5574nM). SPR measurements (Fig. 5l-m) further confirmed this differential binding, yielding *K*_D_ values of 15.2nM and 1210nM for **B1** and **B2**, respectively. The experimental binding affinity differences aligned with PBCNet2.0’s predictions, demonstrating the model’s capability to identify stereochemistry-dependent activity variations and capture subtle three-dimensional interaction features.

To address the second question, we performed a systematic pocket-directed *in silico* alanine scanning^65,66^ by computationally mutating each binding pocket residue to alanine. PBCNet2.0 was then employed to predict changes in binding affinity between each mutated ALDH1B1-**B1** complex and the wild-type complex. Based on the predictions shown in Figure 5n and Supplementary Figure 10, six residues (PHE296, ASN457, LEU477, ILE458, PHE170, and GLU124) were selected for experimental validation through site-directed mutagenesis and binding assays (Supplementary Section 2). PTS assays and SPR analyses demonstrated that alanine substitutions at five of these positions (all except GLU124) significantly impaired **B1** binding (Fig. 5o-t and Supplementary Fig. 11). This high success rate (5/6) in identifying functionally important residues from the selected candidates demonstrates PBCNet2.0’s potential utility in predicting resistance-conferring mutations. For comparison, we also analyzed these residues using the conventional MM-GB/SA method, which ranked PHE296 and PHE170 among its top predictions, while placing ASN457, ILE458, and LEU477 at lower positions (Supplementary Fig. 10b).

The application of PBCNet2.0 led to three key findings: (1) Accurate binding mode prediction through the integration of molecular docking and interpretable AI analysis; (2) Identification of a stereo-chemically pure ALDH1B1 inhibitor, **B1**, with improved potency compared to its racemic predecessor; (3) Successful prediction of potential resistance mutations via computational alanine scanning. These results demonstrate PBCNet2.0’s utility in providing structural and pharmacophoric insights to guide rational probe development and structure-based lead optimization.

## 3. Discussion

We present PBCNet2.0, an advanced deep learning framework designed to predict protein-ligand relative binding affinities. The model leverages an extensive training dataset of 8.6 million protein-ligand pairs and employs a Cartesian tensor-based equivariant network architecture. Our benchmarking demonstrates that PBCNet2.0 achieves comparable accuracy to Schrödinger’s FEP+ simulations in zero-shot prediction scenarios, while maintaining computational efficiency. Through systematic evaluation, we show that PBCNet2.0 accurately captures protein-ligand interaction patterns and geometric features at the atomic level. Notably, the model exhibits emergent ability to predict binding affinity changes resulting from pocket residue mutations, despite not being specifically trained for this task. Both retrospective and prospective validation studies support PBCNet2.0’s utility in expediting lead optimization and probe development processes.

While demonstrating significant capabilities, PBCNet2.0 faces certain methodological limitations common to contemporary structure-based deep learning approaches. A primary constraint lies in its reliance on static protein-ligand binding conformations, which precludes explicit consideration of entropic effects and conformational dynamics during the binding process. Given recent advances in protein-ligand co-folding methods^15,67^, incorporating protein flexibility through integrated structure prediction approaches represents a promising avenue for future development.

Additionally, PBCNet2.0 shows limitations in predicting complete activity loss cases where R-group modifications lead to altered binding modes. This challenge arises from two main factors: First, the scarcity of experimentally validated negative data in public databases, especially regarding binding mode verification and target engagement confirmation. Second, the technical difficulties in accurately measuring binding affinities for compounds with very low activity. Since PBCNet2.0 is designed to compare binding affinity differences between compounds sharing similar binding modes, preliminary conformational analysis and pose validation can help filter out compounds with potentially disrupted binding modes, thereby avoiding this type of inaccurate predictions. Several promising research directions warrant further investigation. First, incorporating high-quality protein mutation datasets into PBCNet2.0’s training could enhance the model’s ability to capture specific protein-ligand interactions. This integration may improve the model’s generalization capabilities across different protein targets and binding scenarios. Second, the current training framework of PBCNet2.0, which focuses on learning generalizable protein-ligand interaction patterns, could serve as an effective pretraining strategy. The pretrained model could then be fine-tuned for specific downstream applications such as virtual screening, potentially addressing the common challenge of ligand structure memorization in traditional training approaches. Third, rigorous validation through real-world drug optimization projects would be essential to assess PBCNet2.0’s practical utility. Such applications would not only evaluate the model’s performance but also identify specific areas requiring improvement, thereby guiding its refinement for drug discovery applications.

## 4. Methods and Data

### 4.1 Datasets

#### 4.1.1 Training set

The training dataset of PBCNet2.0 consists of 8.6 million protein-ligand complex pairs, processed through a systematic pipeline (Supplementary Fig. 12) comprising three sequential stages: (1) Data Collection, (2) Data Preprocessing, and (3) Three-Dimensional Structure Generation. In the initial data collection phase, we extracted approximately 2.81 million protein-ligand binding affinity measurements (IC_50_, EC_50_, K_i_, and K_d_) from the BindingDB Database (2023.12 version)^21^.

In the data preprocessing phase, we systematically filtered and refined the dataset. Molecules with invalid SMILES^68^ structures during RDKit^69^ preparation were excluded. We removed affinity data entries marked as NaN, 0, or containing inequality symbols (‘<‘ or ‘>‘). The data were then organized into groups based on three key identifiers: ‘Entry DOI’, ‘Target name’, and ‘Target source’, ensuring that affinity measurements within each group corresponded to the same target and publication source. Within each group, molecules were clustered using a Tanimoto similarity threshold of 0.5 to establish chemical series of structural analogs. For molecules with multiple reported affinity values, we implemented a quality control protocol: when the maximum-to-minimum affinity ratio was below 3-fold, we used the arithmetic mean; otherwise, the molecule was excluded. For each compound series, we gathered relevant PDB entries based on target information. To select appropriate PDB structures for docking, we analyzed structural similarities between co-crystallized ligands and compounds in each series by calculating Tanimoto coefficients. The PDB structure containing the co-crystallized ligand with the highest similarity was selected for docking. To maintain data quality, chemical series without co-crystallized ligands showing similarity above 0.5 were excluded from further analysis.

Following the confirmation of ligands and PDB structures, we employed Glide^30^ for generating 3D docking conformations. The ligands underwent preprocessing via the Schrödinger LigPrep module using default parameters. The protein structures were prepared using the Protein Preparation Wizard in Schrödinger suite according to standard protocols. Water molecules forming more than three hydrogen bonds with either ligand or receptor atoms were preserved, and receptor grids were generated centered on the co-crystallized ligand. Subsequently, docking was performed using the Glide module with default parameters, generating up to 200 poses per ligand.

Given that structurally similar ligands from the same chemical series typically exhibit comparable binding poses^6^, a fundamental assumption in our modeling approach, pose selection was carefully executed. For each series, we first identified the maximum common substructure (MCS) between individual ligands and their corresponding co-crystallized reference. The root-mean-square deviation (r.m.s.d.) was calculated between each docked pose and the experimentally determined reference pose within the MCS region. Poses with r.m.s.d. ≤ 2.0 Å were deemed acceptable, indicating preservation of the reference binding mode. For ligands with multiple acceptable poses, the one with the highest Glide score was selected. Ligands failing to yield acceptable poses were excluded to maintain data integrity. All operations were conducted using Schrödinger version 2022-3 via the Schrödinger Python API.

Following preprocessing, we systematically paired molecules within each chemical series, implementing a maximum threshold of 15,000 pairs per series. To ensure comprehensive coverage, priority was assigned to previously unpaired molecules during the pairing process. Analysis of the resulting label distribution revealed a normal distribution pattern, with a notable concentration of values within the [−1, 1] range (Supplementary Fig. 13a, left). This concentration presented a potential risk of model overfitting, as the model could achieve artificially low training error by defaulting to predictions near the mean value (Supplementary Fig. 13b). To mitigate this risk, we implemented a data balancing strategy, employing under-sampling in regions of high data density and over-sampling in regions of low density. The resulting balanced dataset exhibited a more uniform label distribution, as illustrated in Supplementary Figure 13a (right).

To ensure data independence, we excluded protein-ligand pairs that overlapped with the test sets. A comprehensive comparison between the training sets of PBCNet2.0 and its predecessor reveals substantial improvements in both target coverage and chemical diversity, as illustrated in Supplementary Figure 13c.

We also addressed potential modification bias, where systematic changes in binding affinity might correlate with specific R-group modifications (e.g., methyl to ethyl conversions). Such bias could lead to model overfitting through simple structural pattern recognition rather than learning meaningful protein-ligand interactions. Our analysis of binding affinity changes across pairs with identical R-group modifications, presented in Supplementary Figure 13d, demonstrates normally distributed activity changes centered around zero, confirming the absence of systematic modification bias.

#### 4.1.2 FEP held-out test set

For model evaluation, we utilized two established FEP benchmark datasets. The first dataset (FEP1) from Wang et al.^8^ contains eight chemical series with experimentally determined binding free energies (ΔG) and corresponding FEP+ predictions. The binding free energies were converted to pIC_50_ values under non-competitive binding assumptions using the equation:

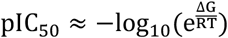

where R represents the gas constant (1.987 × 10^−3^ kcal · K^−1^ · mol^−1^), T is the thermodynamic temperature at 297 K, and e denotes Euler’s number (2.718). The second dataset (FEP2) from Schindler et al.^70^ comprises eight congeneric series targeting pharmaceutically relevant proteins, with experimental IC_50_ measurements. FEP2 exhibits enhanced molecular diversity compared to FEP1, featuring variations in charge properties, ring systems, and core structures. Both datasets underwent logarithmic transformation of binding metrics and standardized ligand pairing protocols consistent with our training methodology. These datasets were combined to form the comprehensive FEP set, with detailed information available in Supplementary Table 1.

#### 4.1.3 SAR-Diff held-out test set

The SAR-Diff test set comprises eight pairs of chemical series, where each pair contains two distinct series. While both series within a pair undergo identical R-group modifications, they exhibit different structure-activity relationships and core scaffolds. This test set specifically evaluates the model’s ability to recognize and interpret protein-ligand interactions, as accurate predictions require analysis of binding pocket interactions rather than simple molecular structure pattern recognition.

The dataset was curated from the ChEMBL database^71^, with series selected from identical assays (as identified by their ChEMBL assay ID) to minimize potential ranking errors within each series. Following data acquisition and ligand structure extraction, we identified appropriate PDB structures and performed docking procedures consistent with our training set protocol to generate 3D structural data. The complete dataset specifications and metadata are provided in Supplementary Table 2.

#### 4.1.4 Mutation test set

To evaluate the model’s capacity for capturing protein-ligand interactions, we assembled a mutation dataset consisting of eight targets from the mutation-induced drug resistance database (MdrDB)^72^. The dataset was filtered to include only mutations within the protein-ligand binding pocket that preserved the original interaction patterns (hydrogen bonds, hydrophobic interactions, salt bridges, π-stacking, π-cation interactions, and halogen bonds). For each target system, we examined a series of mutations while maintaining an identical binding ligand. Each series incorporated a wild-type (WT) complex and multiple mutant (MT) complexes, with corresponding relative binding energy measurements (ΔΔG). Protein and ligand structures were prepared using Schrödinger Maestro, followed by docking simulations to generate binding conformations. The most probable conformations were selected based on their binding modes to establish the mutation test set. The final dataset encompassed mutation data from 8 distinct targets, with comprehensive details provided in Supplementary Table 3.

#### 4.1.5 F-Opt test set

To evaluate the model’s interpretability, we analyzed its performance using protein-ligand fluorine orthogonal multipolar interactions as a model system. We curated a fluorine-focused optimization (F-Opt) dataset from published studies, comprising 10 pairs of protein-ligand complexes. In these pairs, ligand modifications from -H or -CH3 to -F or -CF3 groups led to enhanced binding affinities through the formation of fluorine orthogonal multipolar interactions. Two key criteria guided the dataset selection: (1) explicit literature documentation of fluorine orthogonal multipolar interactions as the primary mechanism for improved activity, and (2) availability of co-crystal structures for structural validation. Complete details of the protein-ligand systems are provided in Supplementary Table 4.

### 4.2 PBCNet2.0 implement

PBCNet2.0 predicts relative binding affinities between pairs of protein-ligand complexes. At its core, the model employs an equivariant graph neural network architecture with Cartesian tensor representations, building upon TensorNet^22^. The model represents atomic interactions in Cartesian coordinates using rank-2 tensors (3 × 3 matrices) as atomic descriptors. These Cartesian rank-2 tensors can be efficiently decomposed into irreducible components under rotational transformations (Equation 1). This decomposition enables straightforward 3 × 3 matrix operations during the message-passing phase, enhancing the computational efficiency of the equivariant network. The following sections detail the irreducible decomposition of rank-2 Cartesian tensors and the architectural components of PBCNet2.0.

#### 4.2.1 Irreducible Cartesian rank-2 tensor decomposition

Here, *X* denotes a rank-2 tensor (3 × 3 matrix) defined in Cartesian coordinates, with *X*_*ij*_ representing the value in the *i* − th row and *j* − th column. Any tensor *X* can be decomposed in the following manner:

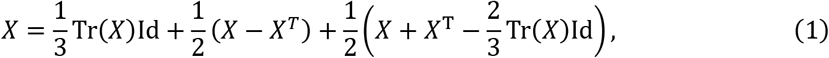

where Tr(*X*) = ∑_*i*_ *X*_*ii*_ is the trace operator, *X*^*T*^ denotes the transpose of *X* and Id is the identity matrix. The first term is proportional to the identity matrix, possessing only one degree of freedom. This term is invariant under rotations and represents a scalar. The second term is a skew-symmetric matrix and contains three independent components that transform as a vector under rotations. The third term is a symmetric and traceless, with five independent components that represent a higher-order tensor. From a representation theory perspective, Equation 1 shows that the 9-dimensional Cartesian rank-2 tensor (*X*) can be decomposed into components of dimensions 1 (scalar), 3 (vector), and 5 (tensor), respectively. We refer to *X* as a “full tensor”, while its components are referred to as scalar (*I*), vector (*A*), and tensor (*S*) features.

The tensor representation can be constructed from relative position vectors in neural networks. For example, given a vector ***r*** = (*r*_*x*_, *r*_*y*_, *r*_*z*_) (e.g., the difference between the coordinates of two nodes in a graph), we can build a well-behaved tensor *X* = *I* + *A* + *S* as follows:

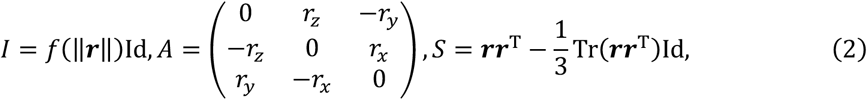

where ‖***r***‖ is the length of ***r***, *f*(·) is mapping function, and ***rr***^T^ denotes the outer product of ***r*** with itself. We can also modify the full tensor by multiplying invariant quantities to each component, *X*^′^ = *f*_*I*_ *I* + *f*_*A*_*A* + *f*_*S*_*S*, where*f*_*I*_, *f*_*A*_, and *f*_*S*_ can be constants or learnable invariant functions (e.g., functions of distances). These modifications preserve the transformation properties of the tensor under rotations.

#### 4.2.2 Graph preprocessing and feature initialization

In our protein-ligand interaction graph construction, we defined the protein pocket as residues within 8Å of the ligand. The graph representation treats each heavy atom in the pocket-ligand complex as a node and each covalent bond as an edge. Additional distance edges were established between protein-ligand and ligand-ligand node pairs with spatial distances ≤ 5.0Å. For protein-protein node pairs, we implemented a dual-threshold approach: if either protein atom was connected to a ligand atom via a distance edge, a distance edge was created when their spatial distance was ≤ 5Å; otherwise, a more stringent threshold of ≤ 3Å was applied. This hierarchical graph construction methodology optimizes the representation of protein-ligand interactions while efficiently reducing computational complexity by limiting distant protein-protein connections.

For feature initialization, while the original TensorNet architecture was limited to atom types and edge lengths as node and edge initial features, we expanded the feature set to include comprehensive chemical descriptors. The enhanced model incorporates nine distinct atomic features (detailed in Supplementary Table 5) and various bond types (listed in Supplementary Table 6), providing a more complete representation of atoms and their local chemical environments.

#### 4.2.3 Initial tensor representations

PBCNet2.0 learns a Cartesian full tensor representation 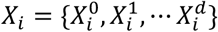 (*d* represents the number of hidden dimensions) for an atom *a*_*i*_. In the following, we describe the computational process using *X*_*i*_ as an example. Atomic representation *X*_*i*_ can be decomposed into scalar, vector and tensor contributions *I*_*i*_, *A*_*i*_, *S*_*i*_ via Equation 1. Conversely, these components can be merged to form *X*_*i*_.

We treat *a*_*i*_ as the central atom, 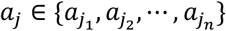 (*n* is the number of neighbor atoms of *a*_*i*_) represents a neighbor atom of *a*_*i*_, and 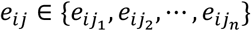 is an edge between *a*_*i*_ and *a*_*j*_. The coordinates of *a*_*i*_ and *a*_*i*_ are presented as ***r***_*i*_ and ***r***_*i*_, the edge vector ***r***_*ij*_ = ***r***_*i*_ − ***r***_*i*_ can be obtained, and ‖***r***_*ij*_‖ presents the length of *e*_*ij*_. We initialize each edge scalar features by the identity matrix 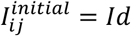, and each vector 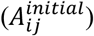 and tensor 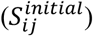 feature are initialized using the normalized edge vectors 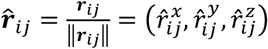 through Equation 2. Thus, the initial full tensor representation of *e*_*ij*_ is 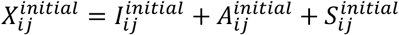. Next, we aim to encode bond- and atom-invariant information, including edge distances, edge and atom chemical features (Supplementary Table 5 and 6), into the initial full tensor 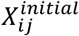. This information is encoded using linear layers, with the distance information extended by a radial basis function (RBF) before encoding:

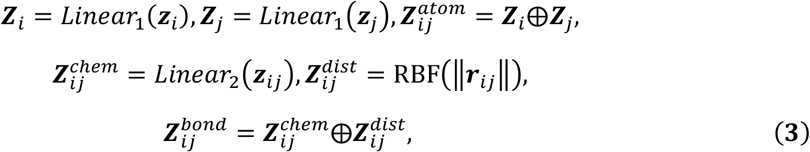

where ***z***_*i*_, ***z***_*j*_ and ***z***_*ij*_ are chemical features of *a*_*i*_, *a*_*j*_ and *e*_*ij*_, respectively. ‖***r***_*ij*_ ‖is the length of *e*_*ij*_. RBF(⋅) and ⨁ are the radial basis function and concatenate operation. ***Z***_*i*_, ***Z***_*j*_ and 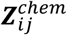 are chemical embeddings of *a*_*i*_, *a*_*j*_ and *e*_*ij*_, while 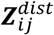 is distance embedding of *e*_*ij*_. 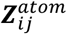 and 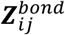 are atom pair-wise and bond-invariant representation of *e*_*ij*_. Then 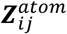 and 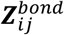 are combined with 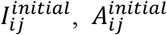 and 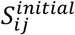 according to:

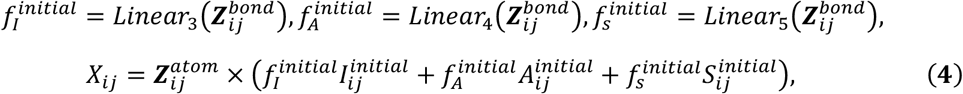

where *X*_*ij*_ is the invariant information-merged initial tensor representation of *e*_*ij*_. It should be noted that the original TensorNet^22^ includes an additional distance decay function in Equation 4. The assumption of distance decay function is that atoms that are farther apart have less influence on each other, which obviously affects the identification of long-range interaction. Moreover, this function also inevitably introduces a bias, where the correlation weights between atom pairs become larger as their distance decreases, and vice versa. In PBCNet2.0, we have eliminated this function. We then get atom-wise initial tensor representation of *a*_*i*_ by aggregating all neighboring edge tensor representations:

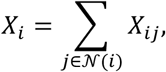

where 𝒩(*i*) means the neighbor set of the central atom *a*^*i*^. At this point, Frobenius norm (an invariant norm) ‖*X*‖_F_ = Tr(*X*^*T*^*X*) of *X*_*i*_ is computed and fed to a series of layers:

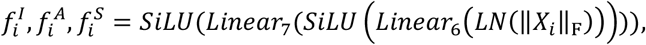

where *LN*(⋅) and *SiLU*(⋅) are layer norm and SiLU activation. After the decomposition of this atom-wise tensor representation *X*_*i*_ into its irreducible representations via Equation 1, 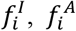 and 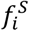 are then used to modify component-wise linear combinations to obtain the final initial atomic tensor representations 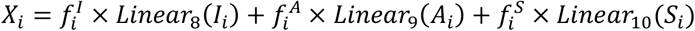.

#### 4.2.4 Tensor representations updating

PBCNet2.0 consists of three message passing layers. We denote the input of the first layer as 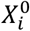 (i.e., the initial atomic tensor representation), the output of the first layer (which becomes the input of the second layer) as 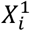, and so on. To introduce the message passing process, we focus on the transition from 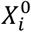 to 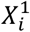 as an example. Firstly, 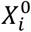 is normalized by

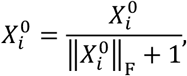

Then 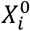 is decomposed into scalar, vector and tensor features 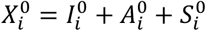. The obtained features are transformed by independent linear combinations to get an updated tensor embedding 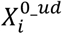 according to

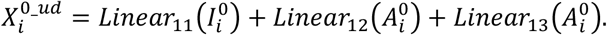

We then aim to merge the information from neighboring atoms used to further update 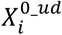. Firstly, the bond-invariant embeddings 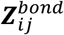 are concatenated with the *O*(3)-invariant scalar embedding of *a*_*i*_ and *a*_*j*_ (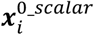 and 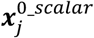, see *Scalar embedding extracting* section for more details), and then are transformed into three invariant terms as follows:

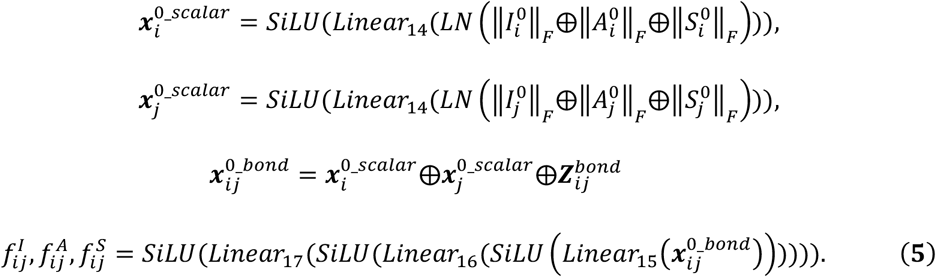

Note that we also eliminated the application of distance decay function compared to the original TensorNet here. Next, the messages *M*_*ij*_ sent from neighbors *a*_*j*_ to atom *a*_*i*_ are computed as:

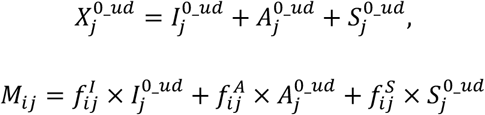

These messages are then aggregated:

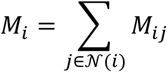

where 𝒩(*i*) means the neighbor set of the central atom *a*^*i*^. Then updated tensor embedding 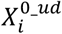 is further updated by *M*_*i*_ with a parity-preserved form:

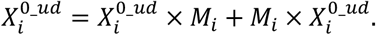

Next, 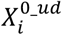 is further decomposed into scalar, vector, and tensor features (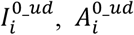, and 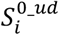) with Equation 1. These features are individually normalized according to:

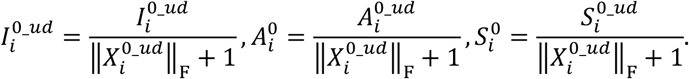

These normalized features are then used to compute independent linear combinations:

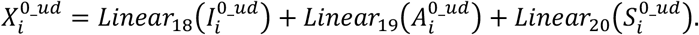

Finally, a residual message 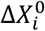 is computed in a parity-preserving manner and used to update 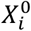 to get 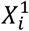:

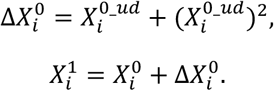

Repeating the above process, we can get 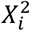 and 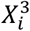 for the subsequent layers.

#### 4.2.5 Scalar embedding extracting

Since the protein-ligand binding affinity is a *O*(3)-invariant property, the Frobenius norms of each component of 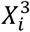, which is also *O*(3)-invariant, are used to get final atomic scalar embedding 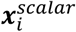,

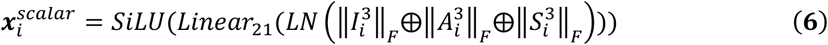

#### 4.2.6 Readout

After the above process, we obtain the final scalar embedding 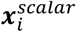 of each atom. We then summed the scalar embeddings from the ligand atoms as the final complex embedding ***x***^*scalar*^,

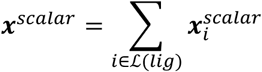

where ℒ(*lig*) is the ligand atom set of a pocket-ligand complex. Finally, we utilize the embedding of the reference complex 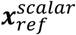 to subtract that of the query complex 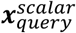, thereby obtaining the difference embedding 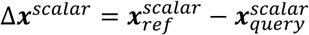, which then be fed into a 3-layer perceptron to obtain the final prediction Δŷ. If the binding affinity of the reference complex *y*_*ref*_ is known, we can get the predicted binding affinity of tested complex ŷ_*query*_ = *y*_*ref*_ − Δ*y*. In the training process, to ensure, to the extent possible, that *MMdell*(Δ***x***^*scalar*^) = −*MMdell*(−Δ***x***^*scalar*^), we input 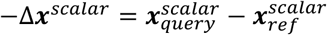 into the fully connected layer as well and calculate the loss together as a data augmentation strategy with a low computational overhead.

### 4.3 Baseline models

To thoroughly evaluate our method, we selected a range of representative baseline models. For traditional approaches, we chose Schrödinger’s **Glide-SP**^30^, **MM-GBSA**^31^, and Schrödinger’s **FEP+**^8^. The Glide-SP and MM-GBSA modules were used with Schrödinger’s default parameters, while the performance of FEP+ is based on the reported results^8,70^.

**Boltz-2**^29^ is a recently published deep learning-based co-folding model. This model takes sequence representations of both the protein and the ligand as input and is capable of predicting the three-dimensional structure of the protein-ligand complex along with its binding affinity. The weight parameters and source code for Boltz-2 were obtained from its original publication. Boltz-2 reports the binding affinity value as lo*g*(IC_50_), derived from an IC_50_ measurement in micromolar units (µM). Therefore, we adopt ‘6 − predi*c*ted va*l*ues’ as the final prediction results.

For deep learning scoring methods, we considered both sequence-based and structure-based approaches. The sequence-based methods include:

- **PSICHIC**^7^. It directly decodes protein-ligand interaction fingerprints from sequence data by incorporating physicochemical constraints. The weights and code of PSICHIC is from its original publication. The binding affinity values output by the model are pK_d_ or pK_i_, and we directly evaluate the model using its outputs.
- **BIND**^32^. It is a graph neural network that uses Evolutionary Scale Modeling 2^73^ (ESM-2) embeddings for proteins and graph-based representations for ligands to perform virtual screening. We ran the model with the default parameters provided in the paper and evaluated it using the IC_50_ values from the output.
- **PLAPT**^33^. It utilizes transfer learning from pretrained transformers like ProtBERT^74^, requiring only 1D protein and ligand string data for predictions. The model’s weights are set to their default values, and the predicted IC_50_ values are directly used for performance calculation.

The structure-based methods differ in terms of input forms (ranging from 2D contact features to 3D graph structures), learning objectives (from direct binding affinity prediction to fine-grained interaction modeling), and model architectures, providing a comprehensive evaluation benchmark. These models comprise:

- **RTMScore**^34^. It predicts binding affinity by fitting a protein-ligand distance likelihood potential, using a Mixture Density Network (MDN) framework. In the pretrained models provided in the paper, we selected the best-performing model, rtmscore_model3. The paper defines a custom affinity score, where higher values indicate stronger binding affinity.
- **GenScore**^35^. Building on RTMScore, the model adds a ranking loss term to the training loss and incorporates a fine-tuning strategy. We evaluated the model using GatedGCN_ft_1.0, which showed the strongest ranking ability according to the results. The output affinity values are also custom-defined.
- **PIGNet2**^11^. The model focuses on modeling the interactions between protein-ligand pairs, refining the types of interactions into hydrogen bonds, metal bonds, van der Waals forces, and hydrophobic interactions. During replication of the paper’s results, we found that the model pda_3 had the performance most closely aligned with the reported results on the FEP test set, so we selected this model for comparison. The predicted binding free energy (ΔG) is converted into pIC_50_.
- **OnionNet-2**^36^. It is a scoring model based on convolutional neural network to predict the binding affinity pKd, where the protein-ligand interactions are characterized by the number of contacts between protein residues and ligand atoms in multiple distance shells. We used the default parameter weights for the model.

### 4.4 Evaluation metrics

Model performance was evaluated using four metrics: the Pearson correlation coefficient (r), Spearman’s rank correlation coefficient (ρ), root-mean-square error (r. m. s. e.), and mean-absolute error (m. a. e.). The correlation coefficients range from 0 to 1, where 1 indicates perfect correlation. The Pearson correlation measures the linear relationship between predicted and experimental binding affinities, while Spearman’s correlation assesses the monotonic relationship between predicted and experimental rankings. Root-mean-square error measures the average magnitude of prediction errors, where lower values indicate better prediction accuracy. Mean absolute error represents the average absolute difference between predicted and experimental values, where lower values indicate better performance.

### 4.5 Interpretability analysis

In the message passing module of PBCNet2.0, the pocket-ligand complex is deemed a whole graph and information on atoms is transmitted across the molecule graph. This module provide atom–atom pairwise interpretation of key inter-molecular interactions, which is helpful for guiding the design a drug for a specific target. The invariant terms 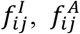, and 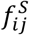 obtained from the Equation 5 can be seen as a measure of importance between atoms *a*_*i*_ and *a*_*j*_. We extracted these invariant terms from the first layer of the message-passing model and computed their average as relevance weights *f*_*ij*_ using the formula:

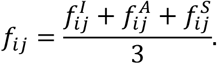

Next, we performed min-max normalization on the weights of all neighboring atoms to obtain the final weights *w*_*ij*_ used in visualization:

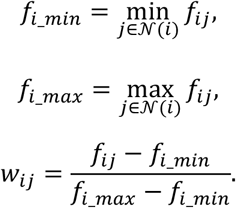

The normalized weights *w*_*ij*_ have a minimum value of 0 and a maximum value of 1, facilitating their use in visualization. An ideal model will tend to assign a higher weight to the pair of atoms which may form a key inter-molecular interaction because they usually have a larger influence on each other’s chemical environment and on ligand binding affinity.

Notably, because the positions of the hydrogen atoms depended heavily on the corresponding program used to add hydrogens, and due to the sheer number of hydrogen atoms which would substantially increase the computational load, we did not take hydrogen atoms into account in the modeling process. Here, for hydrogen-bond donors, the heavy atoms covalently linked with hydrogen atoms were chosen for further analysis, consistent with previous practices in PBCNet.

## 5. Data availability

The training data is from BindingDB source and can be found at Zendo (https://zenodo.org/records/15656365).

The test data are available under an MIT License via GitHub (https://github.com/YuJie-0202/PBCNet2.0).

## 6. Coda availability

The code of this work is available under an MIT License via GitHub at (https://github.com/YuJie-0202/PBCNet2.0).

## 7. Acknowledgments

We gratefully acknowledge financial support from Strategic Priority Research Program of the Chinese Academy of sciences (XDB0830200), National Natural Science Foundation of China (T2225002, 82273855, and 82474143), National Key Research and Development Program of China (2022YFC3400504 and 2023YFC2305904), LingangLaboratory (LGL-8888), Youth Innovation Promotion Association CAS (2023296), and Young Elite Scientists Sponsorship Program by CAST (2023ONRC001). We thank the staff members of the Large-scale Protein Preparation System at the National Facility for Protein Science in Shanghai (NFPS), Shanghai Advanced Research Institute, Chinese Academy of Science, China for providing technical support and assistance in data collection and analysis.

## Notes

### Competing Interest Statement

The authors have declared no competing interest.

### Summary of Updates

We have added the relevant evaluation results of Boltz-2.

