## Supplementary material for "Advancing Ligand Binding Affinity Prediction with Cartesian Tensor-Based Deep Learning": SI

**Author Contributions**

[○]J.Y., X.S., Z.H.F., Z.K.W., and D.H.C. contributed equally to this work. M.Y.Z. and J.Y. designed the research study. J.Y. and X.S. developed the method and wrote the code. M.L.W., H.C.M, and T.C. conducted chemical synthesis. All authors contributed to the analysis of the results. J.Y., M.Y.Z. and X.S. wrote the paper. All authors read and approved the manuscript.

**Note**

The authors declare no competing financial interest.

### 1. Supplementary Sections

#### Supplementary Section 1. Lead optimization tasks

Zero-shot prediction

Zero-shot learning, a subfield of transfer learning, directly applies knowledge learned from training samples to prediction tasks on test samples without any fine-tuning. In our application, it leverages training knowledge to predict binding affinities between unseen protein-ligand pairs, simulating the early stages of lead optimization with limited activity data. By evaluating PBCNet2.0's performance on this task, we can assess its ability to make accurate predictions during the early stages of lead compound optimization, thereby providing effective guidance for subsequent experiments.

The FEP set was used as the test set. For each series of compounds in the test set, we randomly selected one ligand molecule as a reference to infer the absolute binding affinities of the remaining ligand molecules, repeating this process ten times to mitigate randomness in reference molecule selection. Pearson's correlation coefficient (R) and Spearman's rank correlation coefficient (ρ) were used to evaluate the model's ranking capability for each system. During precision assessment, no reference molecule was needed. Instead, we directly paired compounds within each series and calculated the pairwise root-mean-square error (r.m.s.e.$_{pw}$) after model prediction for evaluation.

r.m.s.e.$_{pw}$ is defined using the following:

$$\mathrm{r.m.s.e.}_{\mathrm{pw}} = \sqrt{\frac{1}{N}\sum_{u=1}^{N}(\tilde{y}^{(i,u)} - \hat{y}^{(i.u)})^2}$$

where $(i, u)$ corresponds to a paired sample composed of a query complex and a reference complex (from the same congeneric series); $\tilde{y}^{(i,u)}$ and $\hat{y}^{(i,u)}$ are the true label and prediction result of the paired test sample respectively.

Few-shot prediction

As lead compound optimization progresses, additional activity data becomes available. These structure-activity relationship (SAR) data can be utilized to fine-tune PBCNet2.0, thereby enhancing its predictive performance for specific systems—a process referred to as "few-shot learning." This scenario enables assessment of PBCNet2.0's performance and practical value during the intermediate stages of lead compound optimization.

The FEP set was again used as the dataset for this task. For each test series, 0–10 ligands with known binding affinities were randomly selected as fine-tuning ligands, which also served as reference ligands during inference. The remaining ligands were treated as test ligands. This process was repeated ten times to reduce the impact of randomness. We evaluated the model's ranking capability using the average Spearman and Pearson correlation coefficients obtained on the FEP set during the fine-tuning process.

##### Prioritization experiment

To evaluate the capability of PBCNet2.0 in accelerating lead optimization, we conducted a simulated experiment to determine whether the model could prioritize the selection of compounds with the highest activity ahead of the experimental discovery steps, following the work of Jiménez-Luna and others[1]. We termed this model capability as "selection ability."

This experiment utilized an uncertainty-guided active learning (AL) strategy to strategically prioritize sample acquisition[2]. Data acquisition was simulated as iterative selections within each chemical series, with PBCNet2.0 serving as the active learner. In each series, compounds exhibiting the highest activity were designated as target ligands for identification. When multiple compounds shared the same highest activity, priority was given to the earliest synthesized compound as the target ligand. In the first iteration, the earliest synthesized compound from each chemical series was selected as the reference ligand, and the activities of remaining compounds were evaluated. Subsequently, three ligands with the highest predicted values were selected. If the target ligand was not among these three, they became new reference ligands for the next iteration. In the second iteration, the four existing reference ligands were paired to form a fine-tuning set for improving PBCNet2.0. Fine-tuning guided PBCNet2.0's evaluation of predicted activity values and uncertainties for the remaining ligands. This evaluation directed the prioritization of three ligands according to the predefined sampling method. The iteration continued until successful identification of the target ligand. Here, we employed a model-oriented AL strategy.

Identical to that used in the work of PBCNet, with three evaluation metrics: Advantage order, Advantage ratio, and Efficiency improvement ratio. The calculation formulas are as follows:

$$\text{Advantage order} = \text{Experimental order} - \text{Model selection order}$$

$$\text{Advantage ratio} = \frac{\text{Experimental order} - \text{Model selection order}}{\text{Number of ligands}} \times 100\%$$

$$\text{Efficiency improvement ratio} = \frac{\text{Experimental order} - \text{Model selection order}}{\text{Model selection order}} \times 100\%$$

Here, the 'advantage ratio' represents the theoretical percentage of resources saved when using model-guided lead compound optimization compared to optimization without model guidance. The 'efficiency improvement ratio' indicates the enhancement in efficiency between conducting compound optimization projects with and without the model, assuming the project concludes upon obtaining the most active compound.

#### Supplementary Section 2. Experimental methods

##### Protein Expression and Purification of ENPP1

To produce soluble ENPP1 protein, we engineered the gene encoding the extracellular domain of human ENPP1 (residues 110-926). This gene was fused with the N-terminal secretion signal sequence (residues 1-59) from mouse ENPP2, and a Flag tag was inserted into the C-terminal segment to facilitate purification. The resulting chimeric gene was subsequently cloned into the pcDNA3.1 vector. For protein expression, Expi293F GnTi$^{-/-}$ cells (Thermofisher, A39240) were employed. Five days after transfection, the medium supernatant was collected for protein purification. The purification process commenced with enrichment using Anti-Flag affinity resin, followed by elution of the target protein using Flag peptides. The ENPP1 protein was then further purified using a Superdex 200 Increase column. The purified ENPP1 protein was validated through SDS-PAGE analysis. Finally, the protein was stored in a buffer containing 20 mM HEPES (pH 7.4) and 150 mM NaCl at -80°C.

##### Protein Expression and Purification of ALDH1B1

A truncated form of human ALDH1B1 (residues 20-517) or mutants (F296A, N457A, L477A, I458A, F170A, E124A) was subcloned into the pET-15b vector, incorporating an N-terminal 6×His tag. The recombinant plasmid was transformed into BL21(DE3) Chaperone competent cells (WeiDibio, EM1002S) and grown in lysogeny broth (LB) containing 100 µg/mL ampicillin, 35 µg/mL chloramphenicol, and 1×chaperone inducer. Cultures were incubated at 37°C until reaching an optical density at 600 nm (OD600) of 0.6-0.8, at which point expression was induced with 0.5 mM isopropyl β-D-1-thiogalactopyranoside (IPTG). Cells were subsequently incubated at 16°C for 20 h to allow protein production. Harvested cells were pelleted by centrifugation at 1,600 g for 30 min at 4°C and lysed in buffer (Beyotime, P0013Q) supplemented with protease inhibitor cocktail (Beyotime, P1031) and nuclease (Beyotime, D7121-25KU). Cell disruption was performed by mechanical agitation at room temperature for 25 min. Following clarification by centrifugation at 45,937 g at 4°C for 40 min, the supernatant was subjected to affinity purification using HisTrap columns (Cytiva, 17524801) on an AKTA Pure system (Cytiva). Bound protein was eluted with buffer containing 20 mM HEPES (pH 7.4), 500 mM NaCl, 1 mM TCEP, 500 mM imidazole, and 5% glycerol. Eluted fractions were desalted into storage buffer (20 mM HEPES, pH 7.4, 150 mM NaCl) using HiTrap desalting columns (Cytiva, 29048684) and stored at -80°C until use.

##### In Vitro Enzymatic Activity of ENPP1

The enzymatic activity assay was conducted at 37°C utilizing a transparent 96-well plate, with a total reaction volume of 100 μL. In all assays, p-nitrophenyl-5'-thymidine monophosphate (p-Nph-5'-TMP) served as the substrate. Recombinant human ENPP1 protein was employed at a concentration of 3 nM, alongside a series of diluted compounds, within a reaction buffer consisting of 50 mM Tris-HCl (pH 8.5), 130 mM NaCl, 1 mM $CaCl_2$, and 5 mM KCl. The enzymatic reaction was initiated by introducing the substrate to achieve a final concentration of 100 μM. The production of the reaction product, p-nitrophenol, was continuously monitored using a Spark multifunctional microplate reader from Tecan, by measuring the change in OD405. The $IC_{50}$ values were determined by fitting the data to a logistic curve using GraphPad Prism 9 software.

##### In Vitro Enzymatic Activity of ALDH1B1

The enzymatic activity assay was conducted at room temperature utilizing a 384-well white Optiplate (PerkinElmer, 6007290), with a total reaction volume of 50 μL. Reaction mixtures consisting of assay buffer (100 mM sodium phosphate, 1 mM $MgCl_2$ and 1 mM TCEP, pH 8.0) and 5% DMSO (or varying concentrations of compounds dissolved in DMSO), 1mM $NAD^+$, 100 nM His-ALDH1B1$^{20-517}$ protein, and 1 mM acetaldehyde was added at the end to initiate the reaction. The byproduct of the reaction, NADH, was continuously monitored using a Spark multifunctional microplate reader from Tecan, with an excitation wavelength of 340 nm and an emission wavelength of 460 nm. The $IC_{50}$ values were determined by fitting the data to a logistic curve using GraphPad Prism 9 software.

##### Surface plasmon resonance assay

The surface plasmon resonance (SPR) experiments were performed using a Biacore 8K or a Biacore 1K instrument (Cytiva) at 25°C. Flag-tagged ENPP1 or His-tagged ALDH1B1 (wild type and mutants) protein was covalently immobilized onto a CM5 sensor chip (Cytiva) by a standard amine-coupling procedure in 10 mM sodium acetate of different pH (pH 4.0 or pH 4.5). The running buffer protein contained 10 mM HEPES, pH 7.4, 150 mM NaCl. Compounds were serially diluted into the running buffer and injected onto the sensor chip at a flow rate of 30 μL/min for 120 s (contact phase), followed by 300 s or 200 s of buffer flow (dissociation phase). The equilibrium dissociation constant ($K_D$) value was derived using Biacore Insight Evaluation software (Cytiva).

Protein thermal shift assay

The protein thermal shift assay was conducted using the QuantStudio 5 (Applied Biosystems) to evaluate the compound-induced changes in protein thermal stability. Flag-tagged ENPP1 or His-tagged ALDH1B1 (wild type and mutants) protein was incubated with compounds (50 μM) and SYPRO Orange dye (Sigma, S5692) (5× for ENPP1 and 10× for ALDH1B1). The mixtures were then transferred into 384-well plates (Monad, MQ50701S) with a final volume of 10 μL. The fluorescence signal was recorded as the temperature was gradually raised from 25°C to 95°C. The data were analyzed using the Protein Thermal Shift software v1.4 to determine the $T_m$ value.

**Supplementary Section 3. Compound information of A1 and A2**

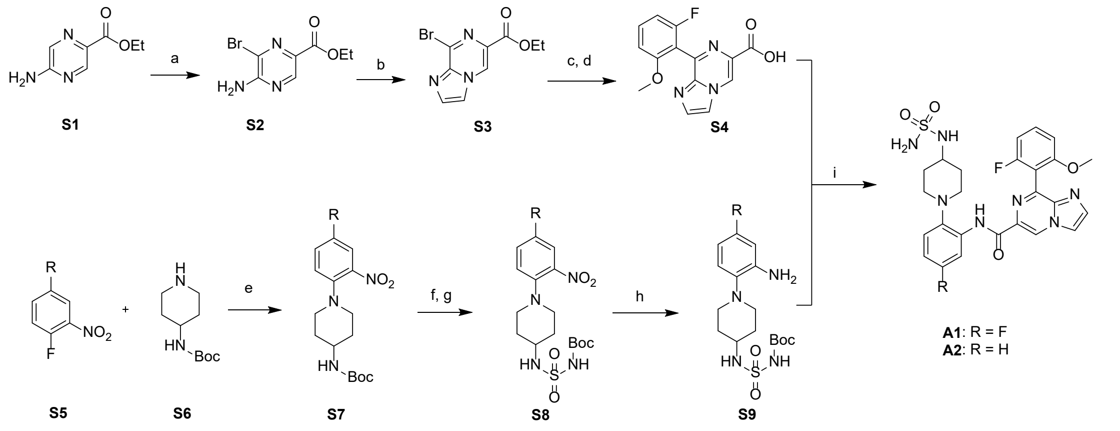

**Scheme S1**. Synthesis of Compounds **A1** and **A2**. Reagents and conditions: (a) NBS, $CH_3CN$, rt, overnight; (b) chloroacetaldehyde, *i*-PrOH, 100°C; (c) aryl boronic acid, Pd(t-$Bu_3P)_2$, DIPEA, dioxane/$H_2O$ (20:1, v/v), 100°C; (d) NaOH, EtOH/$H_2O$ (1:1, v/v), rt; (e) $K_2CO_3$, MeCN, 80°C; (f) HCl, MeOH, 50°C; (g) chlorosulfonyl isocyanate/*t*-BuOH, or sulfonyl chloride, TEA, DCM, 0°C~rt; (h) Fe powder, $NH_4Cl$, MeOH/THF/$H_2O$ (1:1:1, v/v/v), 60°C; (i) EDCI, HOAT, DMF, rt, overnight.

Synthesis of Compounds **A1** and **A2**.

Ethyl 5-aminopyrazine-2-carboxylate **S1** was treated with N-Bromosuccinimide (NBS) in MeCN to generate **S2**. The cyclization of **S2** with chloroacetaldehyde afforded ethyl 8-bromoimidazo[1,2-a]pyrazine-6-carboxylate **S3**, which was followed by Suzuki coupling and hydrolysis reaction to give carboxylic acid **S4**. Compound **S7** was synthesized by heating **S5** and amine **S6** in the presence of $K_2CO_3$ in $CH_3CN$. **S9** was obtained by sulfonylation and reduction. The condensation reaction of **S4** and **S9** in the presence of EDCI and HOAT provided the final product.

Compound **A1** was prepared in a reported procedure,[12] and compound **A2** was prepared in a similar procedure as compound **A1**.

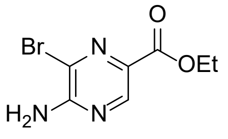

*ethyl 5-amino-6-bromopyrazine-2-carboxylate* **(S2):** yellow solid (90% yield), **$^1$H NMR (500 MHz, Chloroform-*d*)** δ 8.69 (s, 1H), 5.63 (s, 2H), 4.43 (q, *J* = 7.1 Hz, 2H), 1.40 (t, *J* = 7.1 Hz, 3H). LC-MS (ESI) m/z: $[M+H]^+$: Calcd for $C_7H_9BrN_3O_2$ 247.07, found : 247.05.

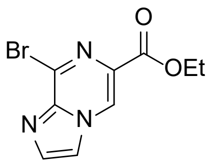

*ethyl 8-bromoimidazo[1,2-a]pyrazine-6-carboxylate* **(S3):** light yellow solid (60% yield), **$^1$H NMR**

**(500 MHz, Chloroform-*d*)** δ 8.94 (d, *J* = 17.2 Hz, 1H), 7.95 – 7.87 (m, 2H), 4.50 (q, *J* = 7.1 Hz, 2H), 1.45 – 1.41 (m, 3H). LC-MS (ESI) m/z: $[M+H]^+$ Calcd for: $C_9H_9BrN_3O_2$ 271.09, found : 226.07.

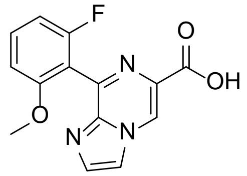

*8-(2-fluoro-6-methoxyphenyl)imidazo[1,2-a]pyrazine-6-carboxylic acid* **(S4):** white solid (50% yield), **$^1$H NMR (500 MHz, DMSO-*d*$_6$)** δ 9.17 (s, 1H), 8.32 (s, 1H), 7.81 (s, 1H), 7.47 – 7.20 (m, 1H), 6.98 – 6.52 (m, 2H). LC-MS (ESI) m/z: $[M+H]^+$ Calcd for: $C_{14}H_{11}FN_3O_3$ 288.07, found : 288.08.

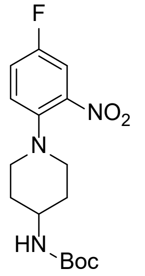

*tert-butyl (1-(4-fluoro-2-nitrophenyl)piperidin-4-yl)carbamate* **(S7-A1):** yellow solid (96% yield ), **$^1$H NMR (500 MHz, DMSO-*d*$_6$)** δ 7.78 (dd, *J* = 8.4, 3.0 Hz, 1H), 7.53 – 7.46 (m, 1H), 7.43 (dd, *J* = 9.2, 5.1 Hz, 1H), 6.88 (d, *J* = 7.9 Hz, 1H), 3.38 (s, 1H), 3.11 – 3.00 (m, 2H), 2.84 – 2.73 (m, 2H), 1.81 – 1.75 (m, 2H), 1.53 – 1.43 (m, 2H), 1.38 (s, 9H). LC-MS (ESI) m/z: $[M+H]^+$ Calcd for: $C_{16}H_{23}FN_3O_4$ 340.37, found: 340.33.

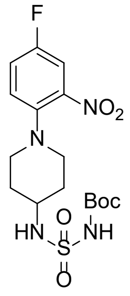

*tert-butyl (N-(1-(4-fluoro-2-nitrophenyl)piperidin-4-yl)sulfamoyl)carbamate* **(S8-A1):** yellow solid (75% yield), **$^1$H NMR (500 MHz, Chloroform-*d*)** δ 7.52 (dd, *J* = 8.0, 2.9 Hz, 1H), 7.26 – 7.20 (m, 1H), 7.22 – 7.15 (m, 2H), 5.17 (d, *J* = 7.1 Hz, 1H), 3.53 – 3.42 (m, 1H), 3.22 – 3.15 (m, 1H), 2.92 – 2.84 (m, 2H), 2.10 – 2.05 (m, 2H), 1.83 – 1.72 (m, 1H), 1.50 (s, 9H). LC-MS (ESI) m/z: $[M+Na]^+$ Calcd for: $C_{16}H_{23}FN_4O_6SNa$ 441.43, found: 441.30.

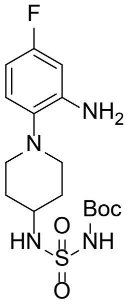

*tert-butyl (N-(1-(2-amino-4-fluorophenyl)piperidin-4-yl)sulfamoyl)carbamate* **(S9-A1):** white solid (66 % yield), **$^1$H NMR (500 MHz, DMSO-$d_6$)** δ 10.82 (s, 1H), 7.75 (d, *J* = 7.4 Hz, 1H), 6.88 (dd, *J* = 8.7, 6.1 Hz, 1H), 6.43 (dd, *J* = 11.1, 3.0 Hz, 1H), 6.30 – 6.22 (m, 1H), 5.01 (s, 2H), 3.20 (s, 1H), 2.93 (dd, *J* = 9.8, 6.3 Hz, 2H), 2.54 (s, 2H), 1.85 (d, *J* = 12.0 Hz, 2H), 1.71 – 1.59 (m, 2H), 1.43 (s, 9H). LC-MS (ESI) m/z: $[M+H]^+$ Calcd for: $C_{16}H_{26}FN_4O_4S$ 389.46, found: 389.20.

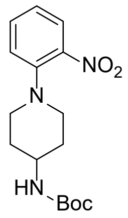

*tert-butyl (1-(2-nitrophenyl)piperidin-4-yl)carbamate* **(S7-A2):** yellow solid (96 % yield), **$^1$H NMR (500 MHz, Chloroform-*d*)** δ 7.76 (dd, *J* = 8.1, 1.6 Hz, 1H), 7.48 – 7.42 (m, 1H), 7.13 (dd, *J* = 8.4, 1.2 Hz, 1H), 7.06 – 6.97 (m, 1H), 4.61 – 4.47 (m, 1H), 3.61 (s, 1H), 3.31 – 3.16 (m, 2H), 3.07 – 2.80 (m, 2H), 2.08 – 1.91 (m, 2H), 1.65 – 1.55 (m, 2H), 1.45 (s, 9H). LC-MS (ESI) m/z: $[M+H]^+$ Calcd for: $C_{16}H_{24}N_3O_4$ 322.17, found: 322.21.

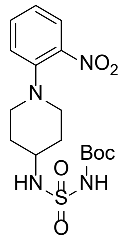

*tert-butyl (N-(1-(2-nitrophenyl)piperidin-4-yl)sulfamoyl)carbamate* (**S8-A2**)*:* yellow solid (84% yield), **$^1$H NMR (500 MHz, Chloroform-*d*)** δ 7.78 (dd, *J* = 8.1, 1.6 Hz, 1H), 7.52 – 7.44 (m, 1H), 7.24 (s, 1H), 7.14 (dd, *J* = 8.2, 1.2 Hz, 1H), 7.10 – 7.01 (m, 1H), 5.20 (d, *J* = 7.2 Hz, 1H), 3.53 – 3.39 (m, 1H), 3.32 – 3.18 (m, 2H), 2.99 – 2.81 (m, 2H), 2.12 – 2.05 (m, 2H), 1.85 – 1.72 (m, 2H), 1.51 (s, 9H). LC-MS (ESI) m/z: $[M+Na]^+$ Calcd for: $C_{16}H_{22}N_4O_6SNa$ 423.14, found: 423.22.

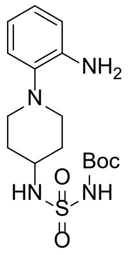

*tert-butyl (N-(1-(2-aminophenyl)piperidin-4-yl)sulfamoyl)carbamate* (**S9-A2**)*:* yellow solid (64% yield), **$^1$H NMR (500 MHz, Chloroform-*d*)** δ 7.05 – 6.85 (m, 2H), 6.80 – 6.64 (m, 2H), 5.34 (d, *J* = 7.3 Hz, 1H), 3.45 (d, *J* = 10.9 Hz, 1H), 3.27 – 3.04 (m, 2H), 2.84 – 2.62 (m, 2H), 2.21 – 1.95 (m, 2H), 1.80 – 1.66 (m, 2H), 1.50 (s, 9H).LC-MS (ESI) m/z: $[M+H]^+$ Calcd for: $C_{16}H_{27}N_4O_4S$ 371.17, found: 371.21.

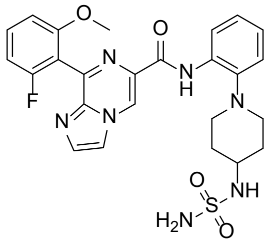

*8-(2-fluoro-6-methoxyphenyl)-N-(2-(4-(sulfamoylamino)piperidin-1-yl)phenyl)imidazo[1,2-a]pyrazine-6-carboxamide* **(A2).** Eluting with dichloromethane/methanol = 20/1, 75% yield. **$^1$H NMR (500 MHz, Methanol-*d₄*)** *δ* 9.35 (s, 1H), 8.47 (d, *J* = 6.5 Hz, 1H), 8.32 (s, 1H), 7.96 – 7.89 (m, 1H), 7.65 (d, *J* = 6.8 Hz, 1H), 7.23 (d, *J* = 6.3 Hz, 1H), 7.16 – 7.05 (m, 3H), 7.01 (t, *J* = 8.6 Hz, 1H), 3.80 (s, 3H), 3.25 – 3.15 (m, 1H), 3.01 – 2.88 (m, 2H), 2.78 – 2.69 (m, 2H), 1.95 – 1.88 (m, 2H), 1.57 – 1.42 (m, 2H). **$^{13}$C NMR (150 MHz, Methanol-$d_4$)** *δ* 163.2, 162.1, 161.6, 160.7, 160.6, 145.1, 144.1, 141.0, 136.1, 134.5, 134.1, 133.7 (d, *J* = 10.3 Hz), 126.2, 125.6, 122.8, 122.0, 119.9, 118.4, 114.2 (d, *J* = 18.2 Hz), 109.3 (d, *J* = 21.5 Hz), 108.8, 56.9, 52.2, 34.4. HRMS (ESI) *m/z* calculated for $C_{25}H_{26}FN_7O_4S$ $[M+H]^+$ 540.1829, found: 540.1833. HPLC purity: 98.8%.

Copies of $^{1}H$ and $^{13}C$ NMR spectra

$^{1}H$ NMR spectra *for ethyl 5-amino-6-bromopyrazine-2-carboxylate* ***(S2):***

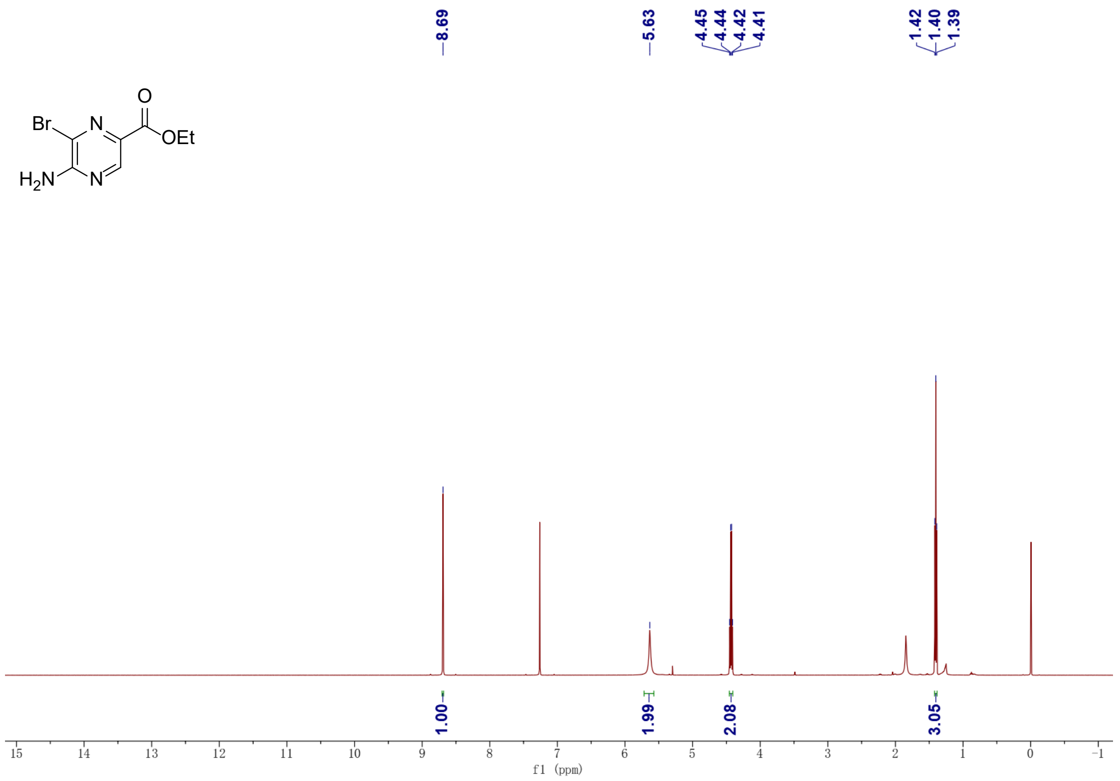

$^{1}H$ NMR spectra *for ethyl 8-bromoimidazo[1,2-a]pyrazine-6-carboxylate* ***(S3):***

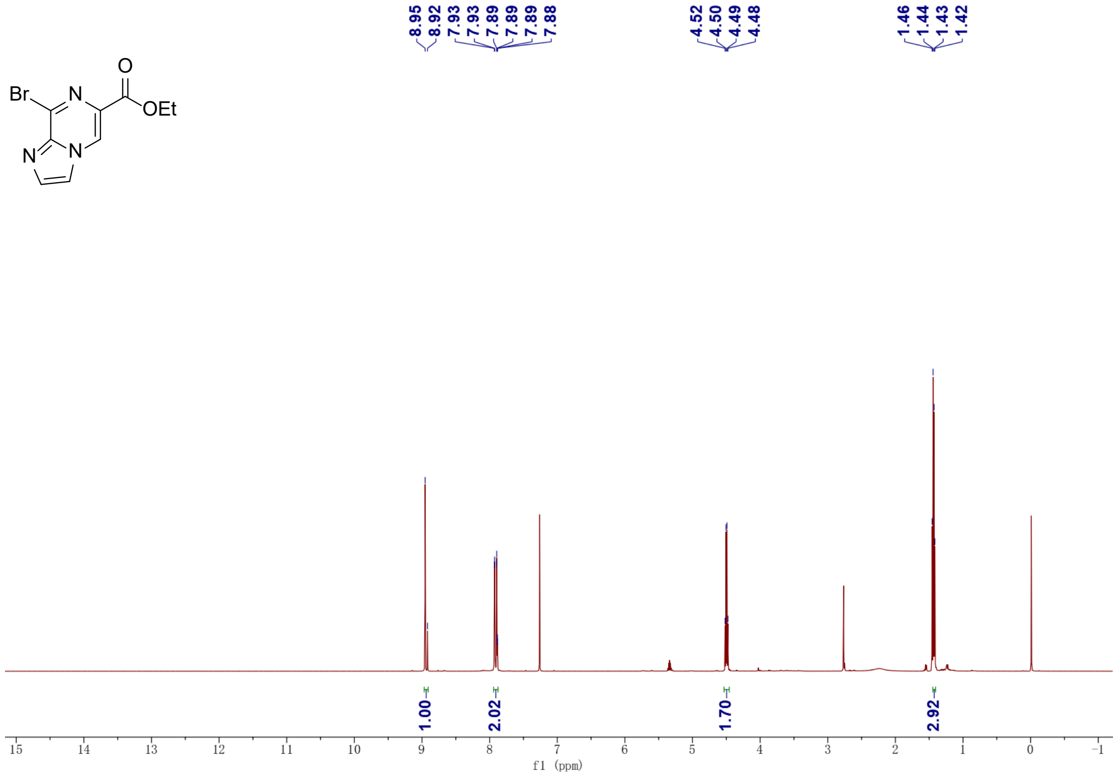

[1]H NMR spectra *for 8-(2-fluoro-6-methoxyphenyl)imidazo[1,2-a]pyrazine-6-carboxylic acid* ***(S4):***

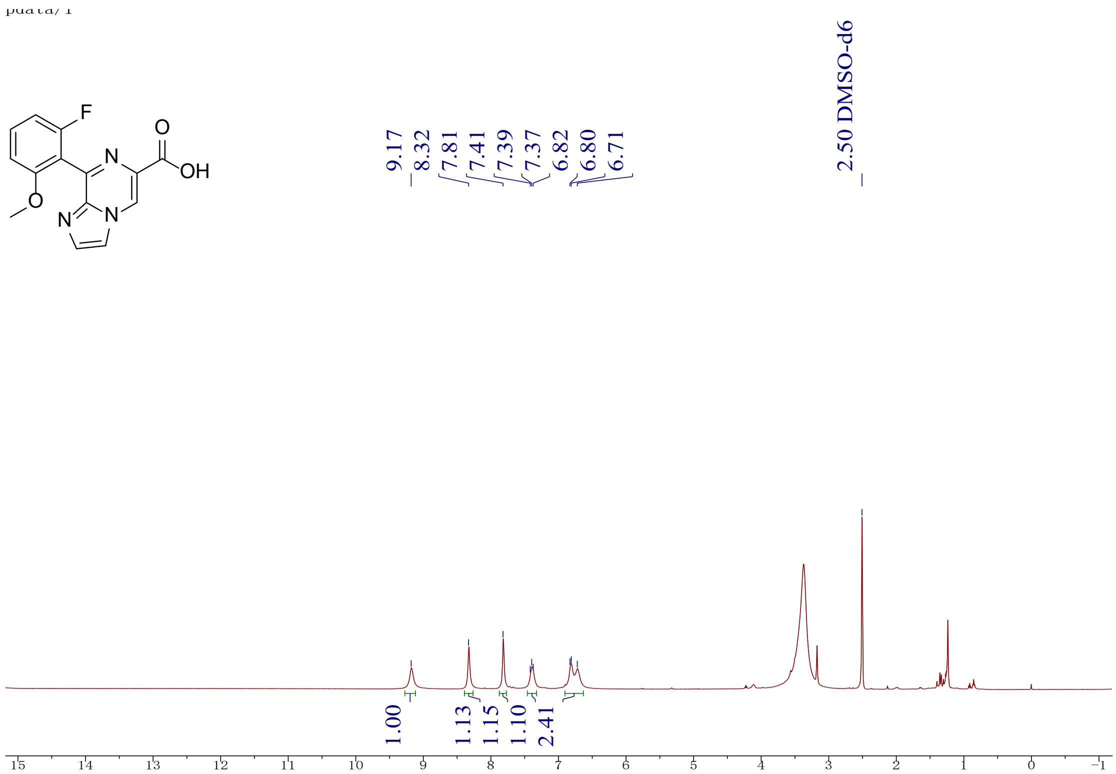

[1]H NMR spectra *for tert-butyl (1-(4-fluoro-2-nitrophenyl)piperidin-4-yl)carbamate* ***(S7-A1):***

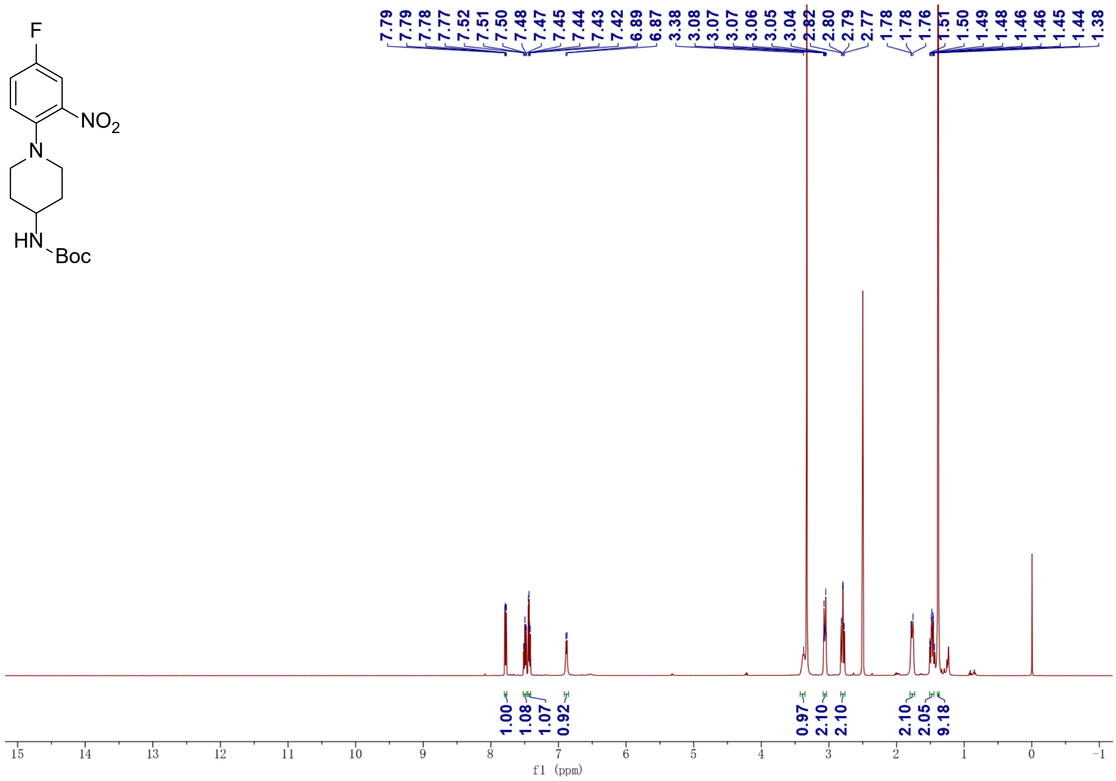

[1]H NMR spectra *for tert-butyl (N-(1-(4-fluoro-2-nitrophenyl)piperidin-4-yl)sulfamoyl)carbamate* ***(S8-A1):***

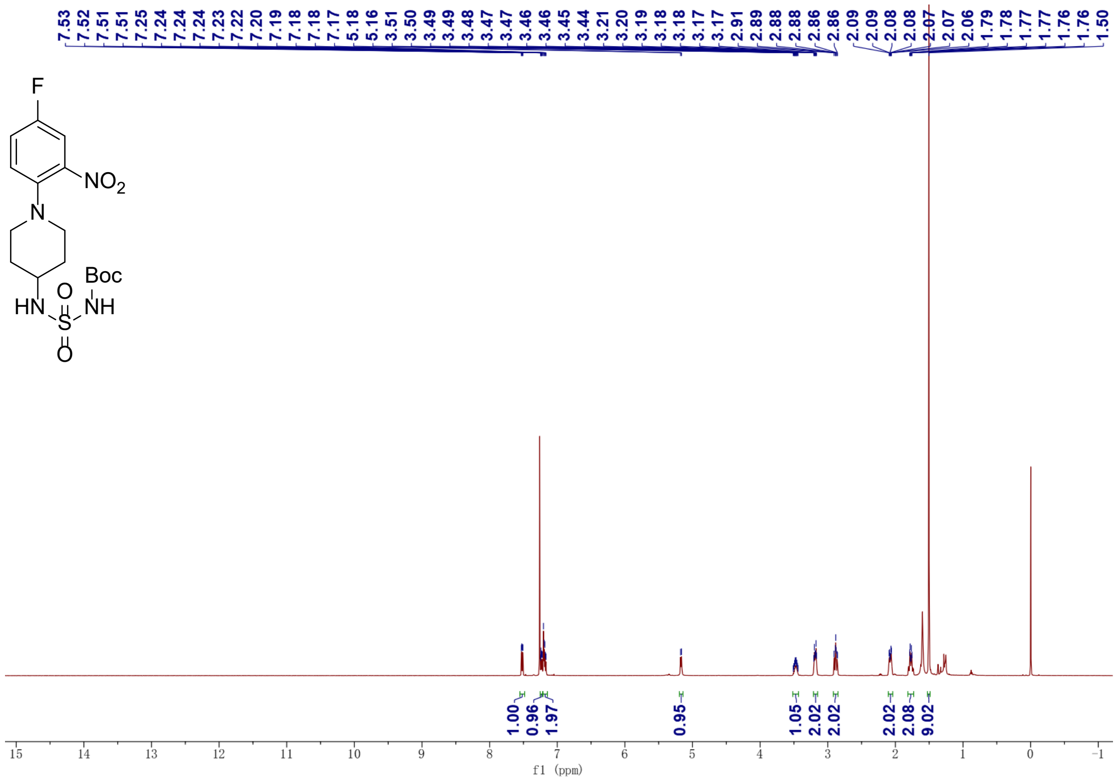

[1]H NMR spectra *for tert-butyl (N-(1-(2-amino-4-fluorophenyl)piperidin-4-yl)sulfamoyl)carbamate* ***(S9-A1):***

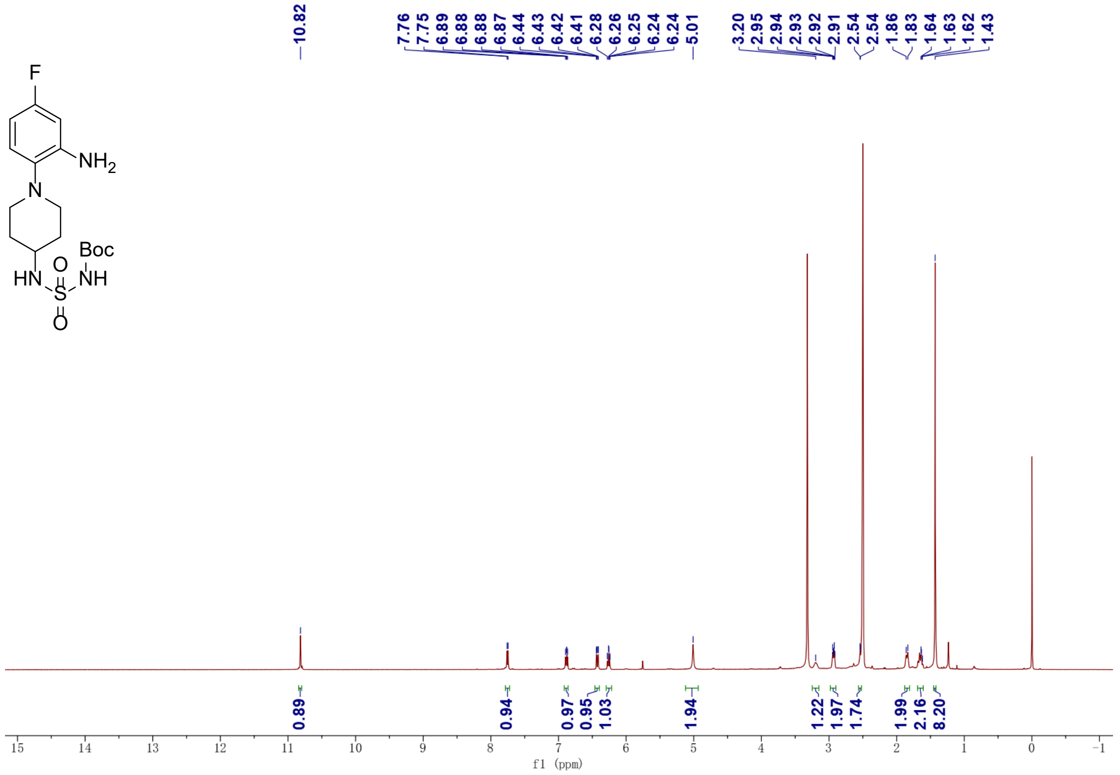

$^{1}$H NMR spectra *for tert-butyl (1-(2-nitrophenyl)piperidin-4-yl)carbamate* ***(S7-A2):***

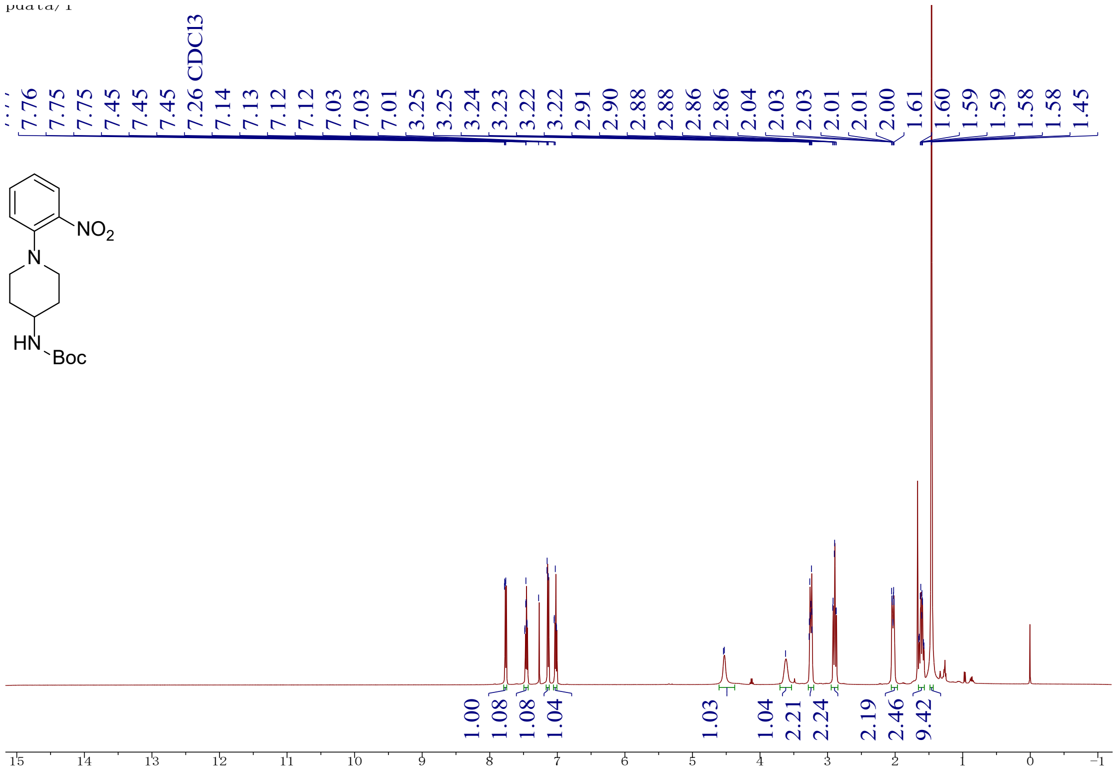

$^{1}$H NMR spectra *for tert-butyl (N-(1-(2-nitrophenyl)piperidin-4-yl)sulfamoyl)carbamate* ***(S8-A2):***

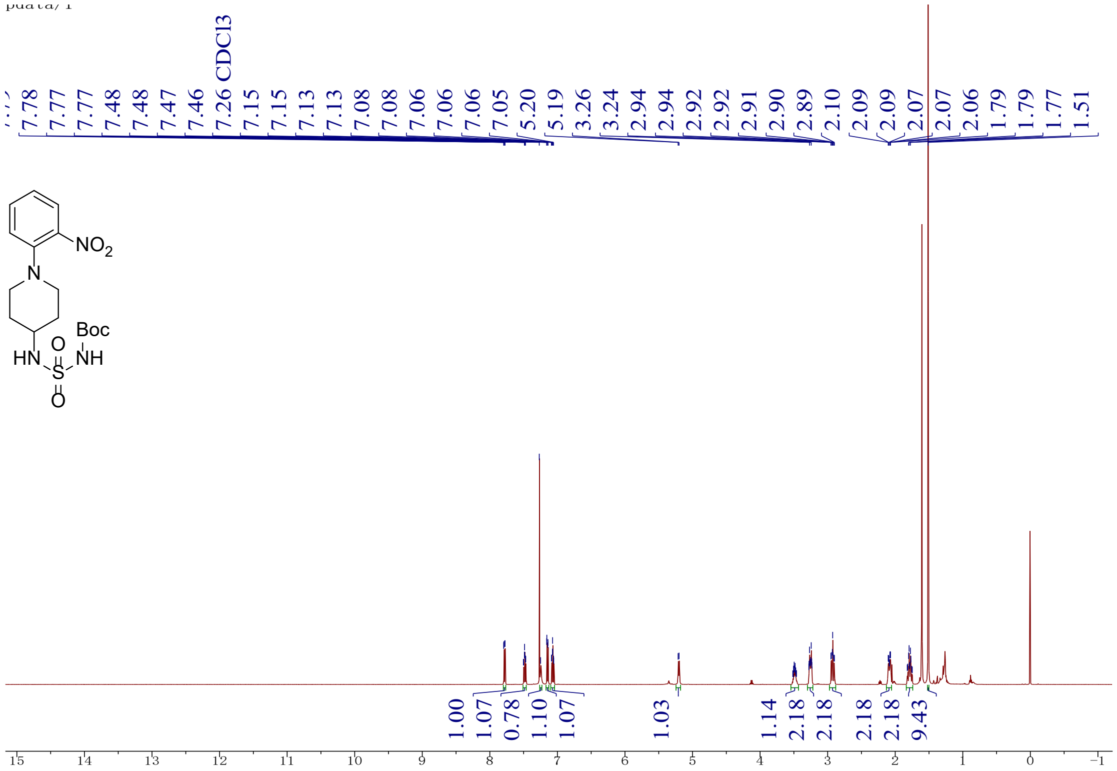

$^1$H NMR spectra *for tert-butyl (N-(1-(2-aminophenyl)piperidin-4-yl)sulfamoyl)carbamate* ***(S9-A2):***

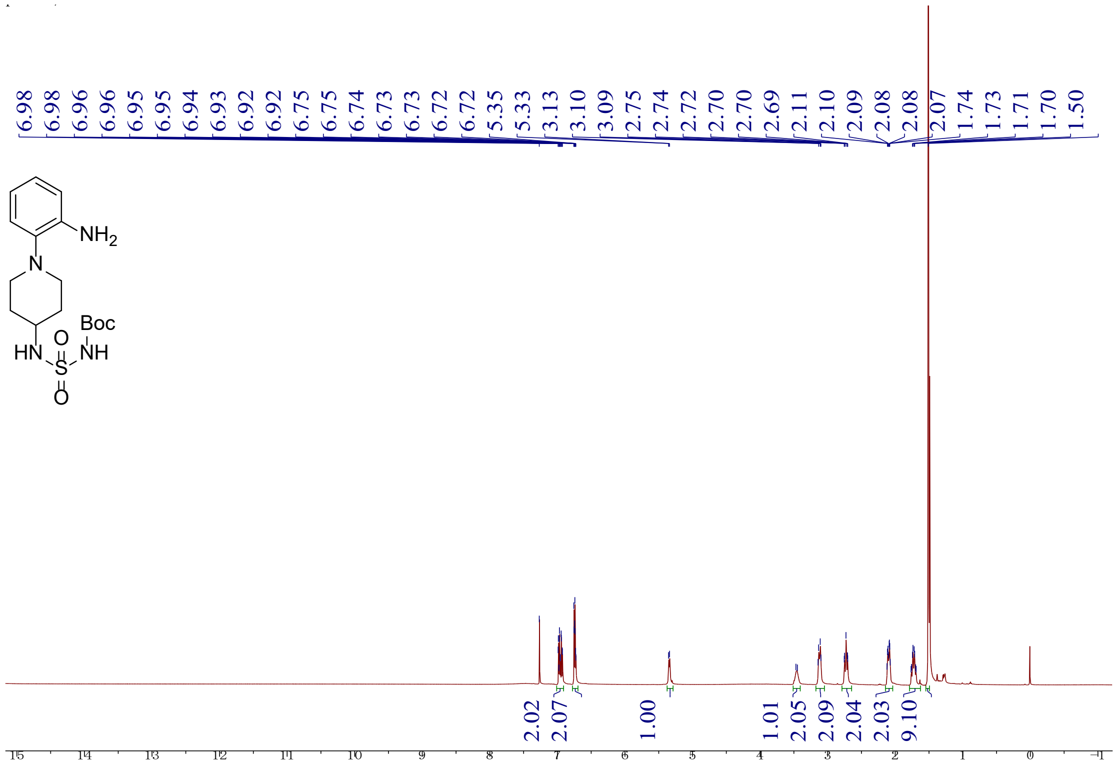

$^1$H and $^{13}$C NMR spectra *for 8-(2-fluoro-6-methoxyphenyl)-N-(2-(4-(sulfamoylamino)piperidin-1-yl)phenyl)imidazo[1,2-a]pyrazine-6-carboxamide* (**A2**)

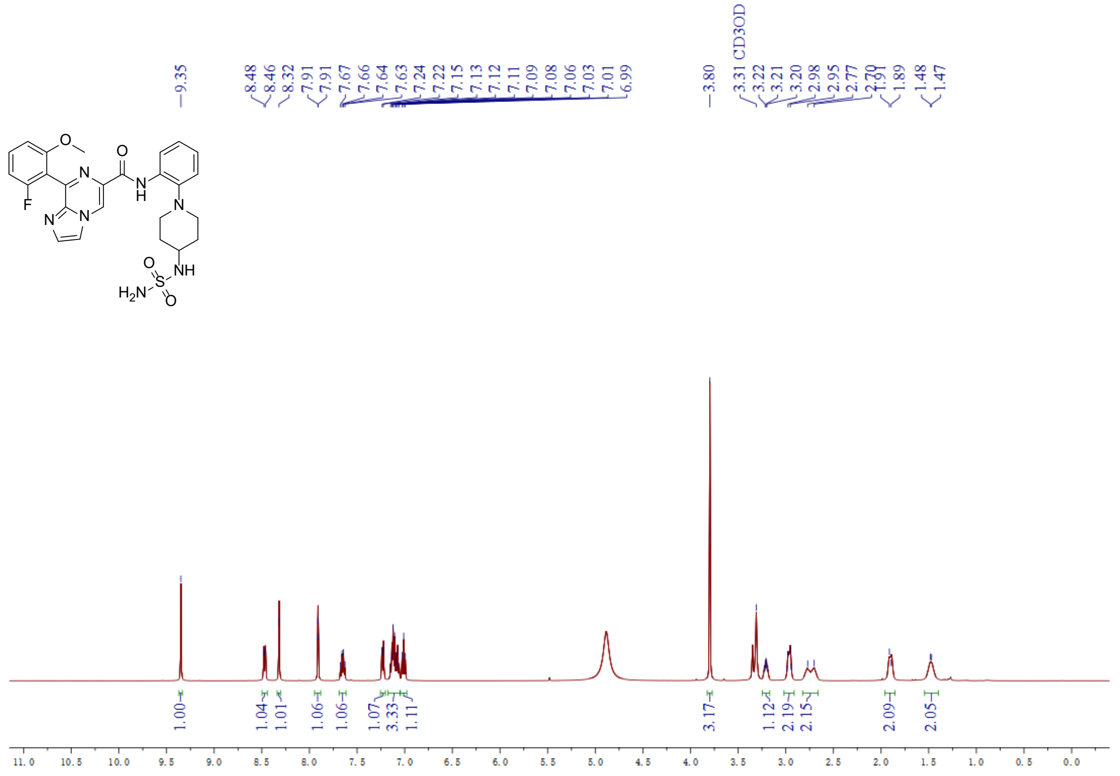

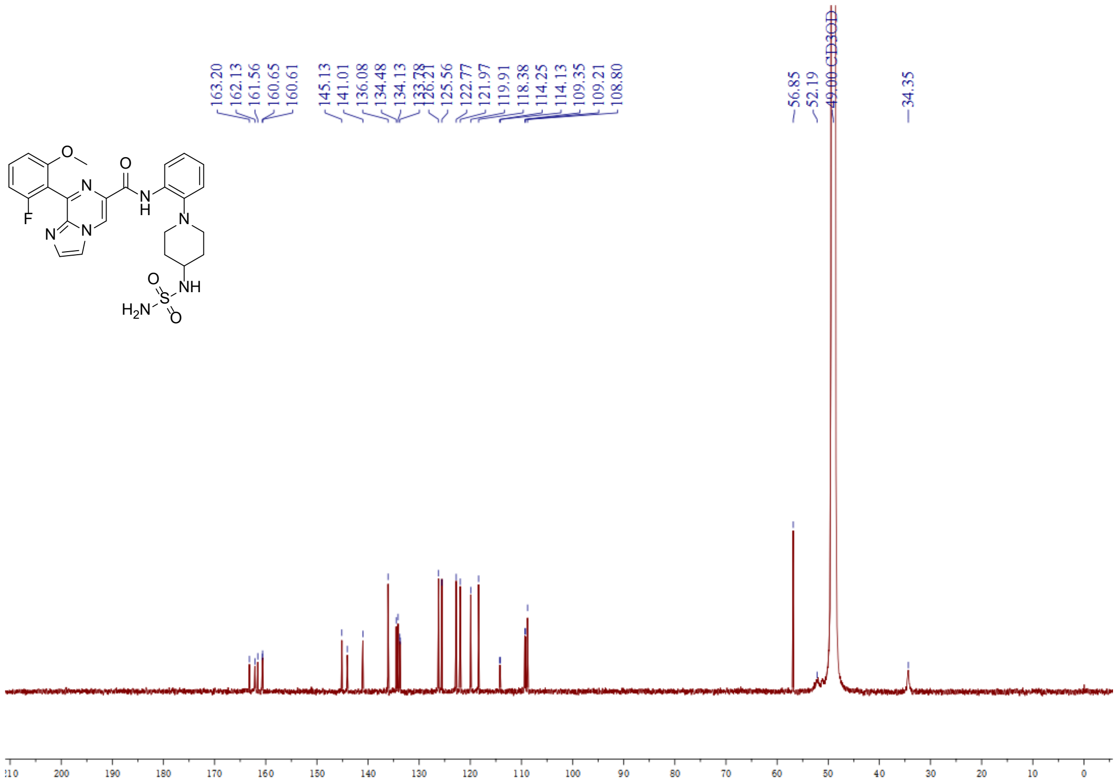

HPLC spectra for compound **A2**

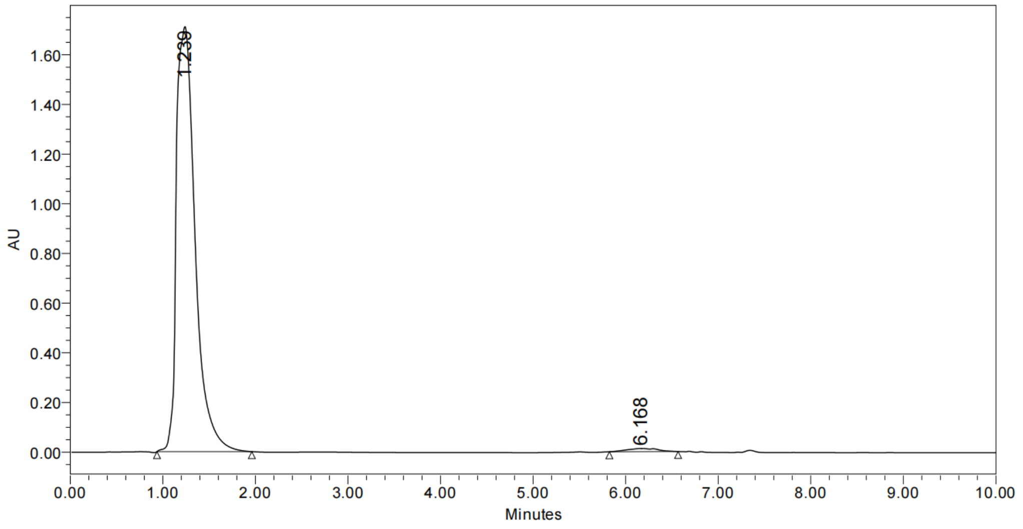

| | RT | Area | % Area | Height |
|---|---|---|---|---|
| 1 | 1.239 | 24021326 | 98.81 | 1710484 |
| 2 | 6.168 | 290438 | 1.19 | 12518 |

ESI HR-MS spectra for compound **A2**

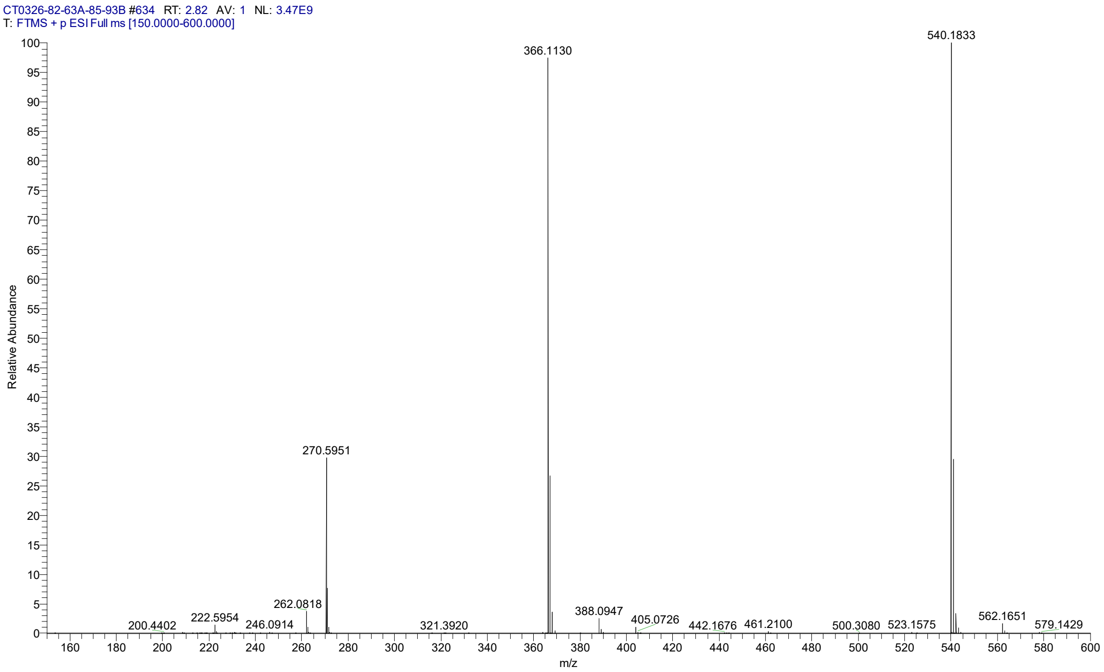

#### Supplementary Section 4. Compound information of B1 and B2

Synthesis of derivatives **B1** and **B2** via Povarov Cyclization

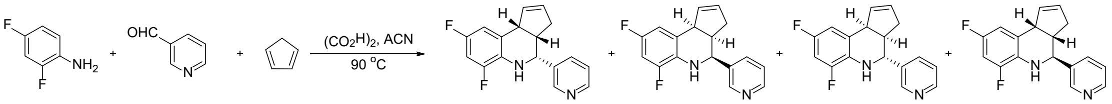

1 2 3 B1 B2 B3 B4

A 20 mL screw-top vial was charged with 2,4-difluoroaniline (**1**, 2.0 mmol, 258 mg, 1.0 equiv.), 3-formylpyridine (**2**, 2.0 mmol, 214 mg, 1.0 equiv.), freshly cracked cyclopentadiene (**3**, 8.0 mmol, 603 µL, 4.0 equiv) and acetonitrile (10.0 mL). To this mixture was added oxalic acid (2.2 mmol, 198 mg, 1.1 equiv.), and the vial was sealed with a Teflon-lined cap and heated at 90 °C for 8 hours. The mixture was cooled to ambient temperature and diluted with dichloromethane, then it was sequentially washed with 1 M aqueous HCl, saturated $NaHSO_3$, and brine. The organic layer was dried over $Na_2SO_4$ and concentrated. The residue was purified via silica gel chromatography, followed by slurry treatment with petroleum ether and ethyl acetate to obtain a mixture of **B1** and **B2** (248 mg, 44% yield), which was then separated by SFC to afford the single isomers of **B1** and **B2**. The compounds **B3** and **B4** (10 mg) were less than 5% yield with about 82% purity.

HPLC trace of **B1**and **B2**

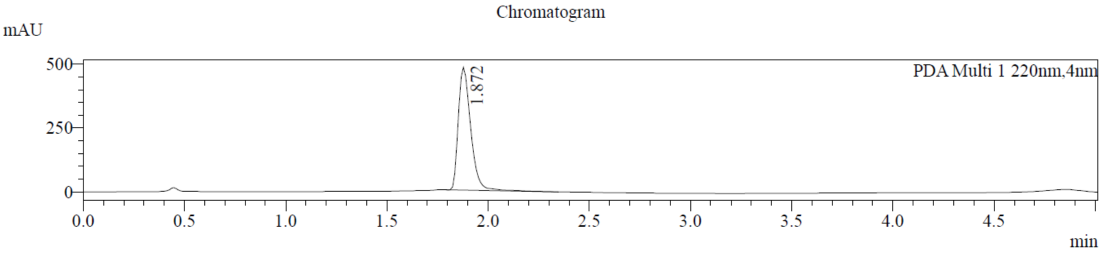

SFC analyst of **B1**and **B2**

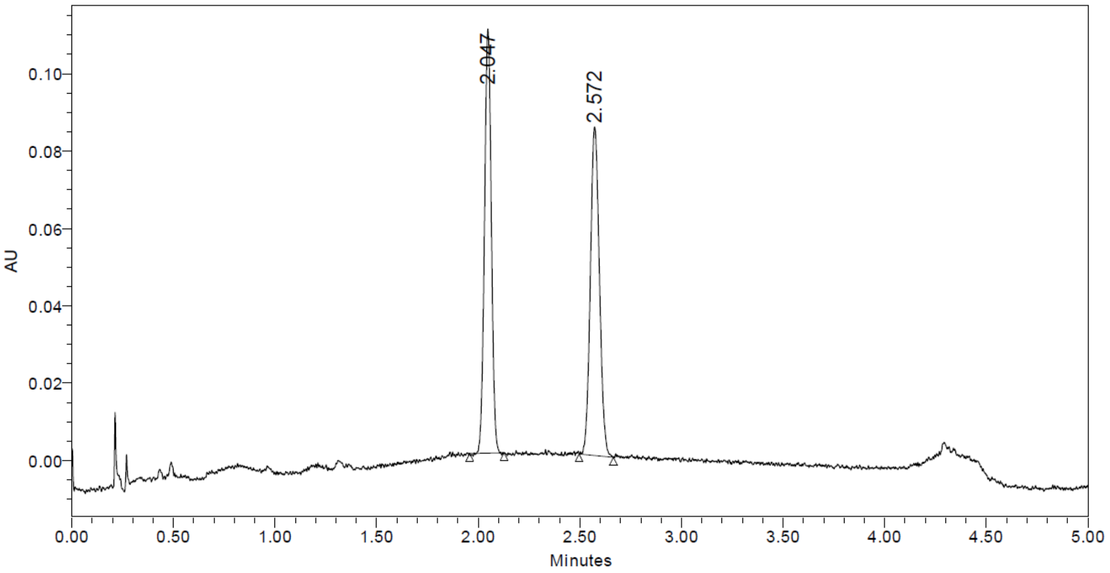

| | RT | Area | % Area |
|---|---|---|---|
| 1 | 2.047 | 257514 | 49.41 |
| 2 | 2.572 | 263658 | 50.59 |

Preparative separation method

**Instrument:** Waters 150 preparative SFC(SFC-26); **Column:** ChiralCel OD, 250×30mm I.D., 10µm; **Mobile phase:** A for $CO_2$ and B for Methanol; **Gradient:** B 30 %; **Flow rate:** 120 mL /min; **Back pressure:** 100 bar; **Column temperature:** 38°C; **Wavelength:** 220nm; **Cycle time**: ~5 min; **Sample preparation:** Compound was dissolved in ~100 mL Methanol/DCM; **Injection:** 4 mL per injection. **Work up:** After separation, the fractions were dried off via rotary evaporator at bath temperature 40°C to get the desired isomers.

Data for **B1**

1. Chemical Structural for **B1**

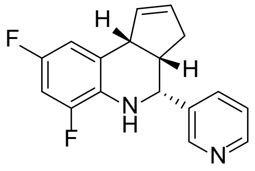

**B1**

2. HPLC trace of **B1**

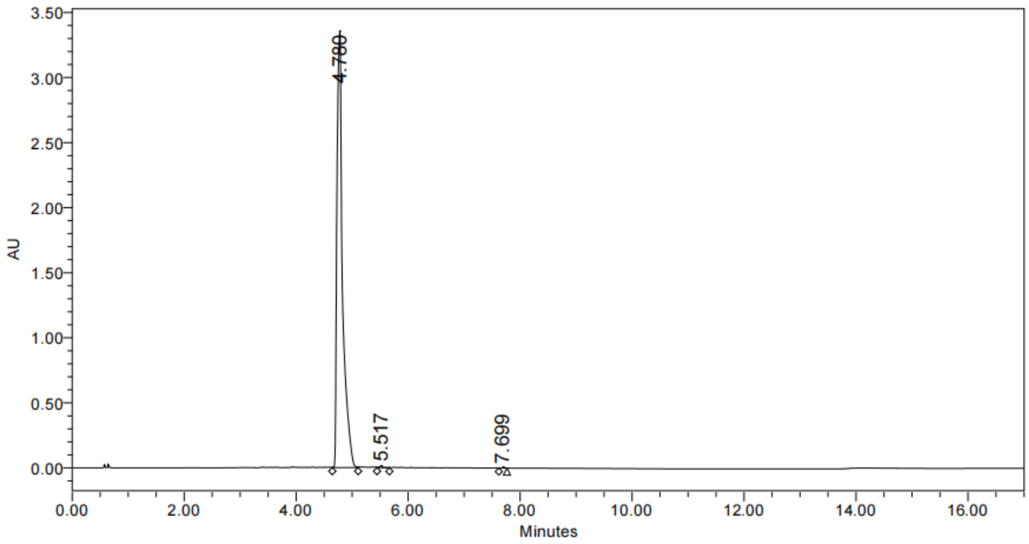

Peak Results

| | RT | Area | Height | % Area |
|---|---|---|---|---|
| 1 | 4.780 | 23582982 | 3357120 | 99.66 |
| 2 | 5.517 | 60602 | 16512 | 0.26 |
| 3 | 7.699 | 19126 | 7318 | 0.08 |

3. SFC analyst of **B1**

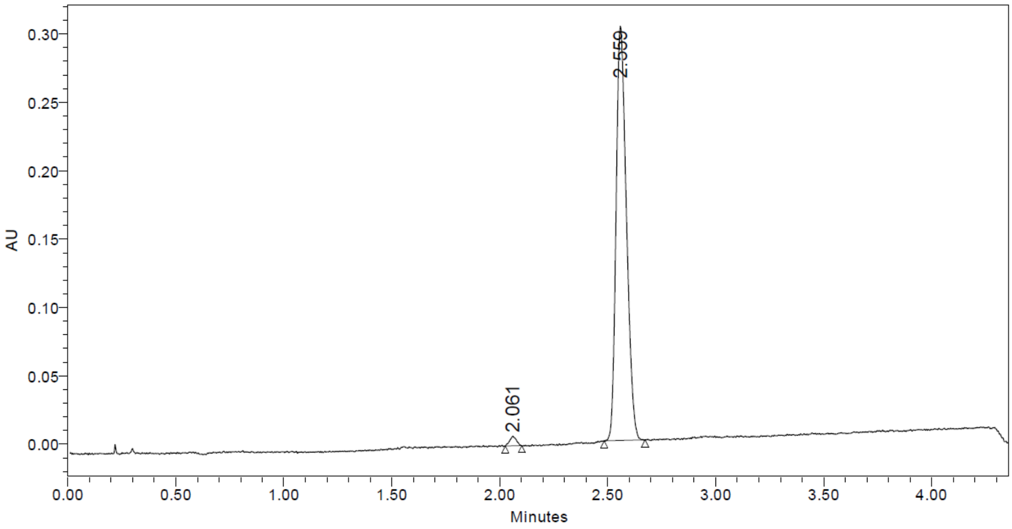

| | RT | Area | % Area |
|---|---|---|---|
| 1 | 2.061 | 14834 | 1.45 |
| 2 | 2.559 | 1006054 | 98.55 |

4. HRMS of **B1**

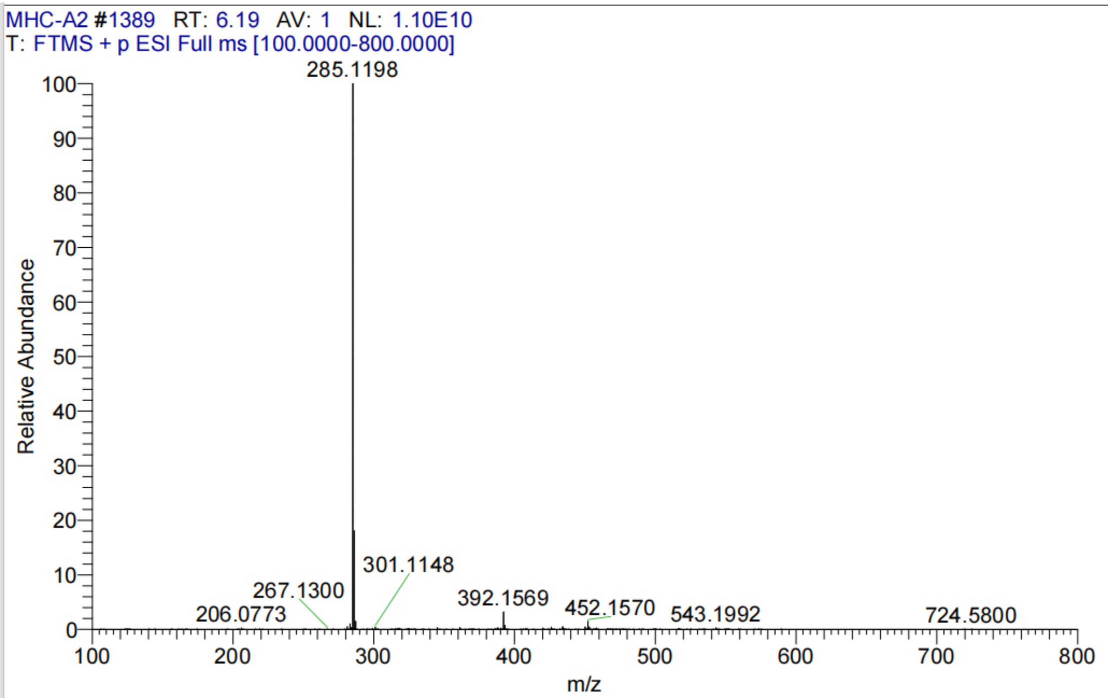

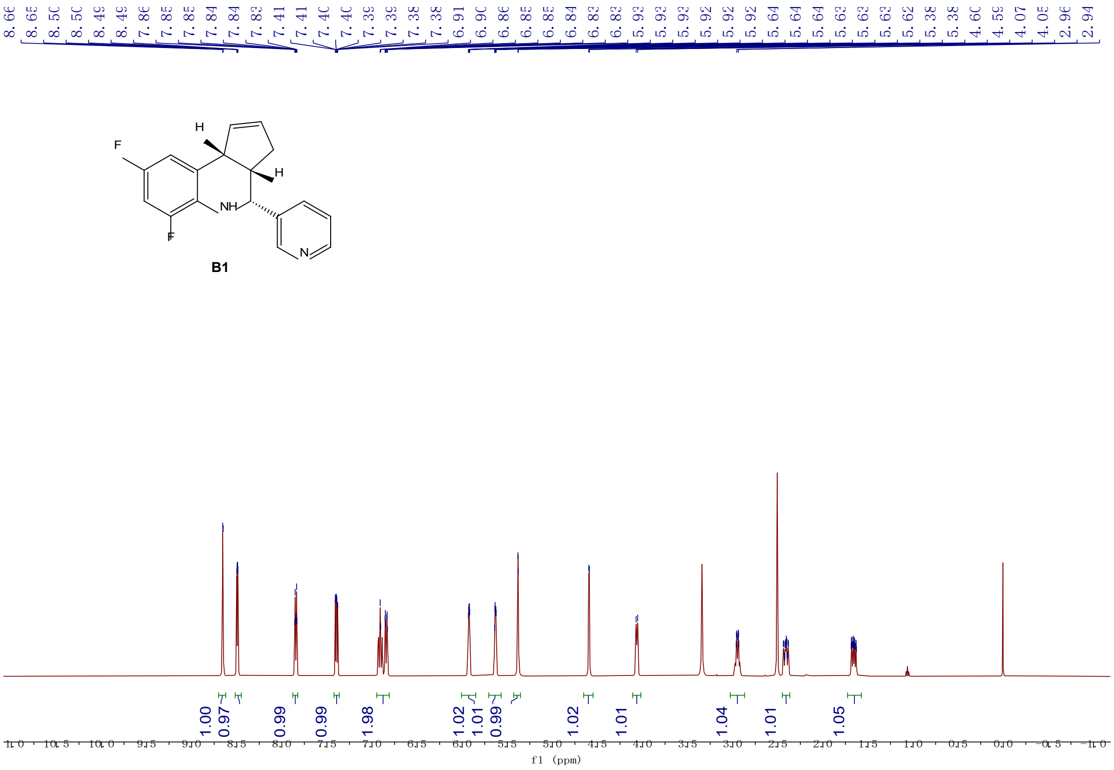

**[1]H NMR (500 MHz, DMSO-$d_6$)** δ 8.65 (d, $J$ = 2.3 Hz, 1H), 8.49 (dd, $J$ = 4.8, 1.7 Hz, 1H), 7.85 (dt, $J$ = 7.9, 2.0 Hz, 1H), 7.39 (ddd, $J$ = 7.8, 4.8, 0.8 Hz, 1H), 6.95 – 6.81 (m, 2H), 6.01 – 5.85 (m, 1H), 5.63 (dq, $J$ = 4.3, 1.4 Hz, 1H), 5.38 (d, $J$ = 2.0 Hz, 1H), 4.59 (d, $J$ = 3.3 Hz, 1H), 4.06 (d, $J$ = 8.6 Hz, 1H), 2.95 (ddd, $J$ = 9.7, 3.3, 1.5 Hz, 1H), 2.41 (ddt, $J$ = 14.2, 7.4, 2.4 Hz, 1H), 1.66 (dddd, $J$ = 16.2, 8.7, 2.7, 1.5 Hz, 1H).

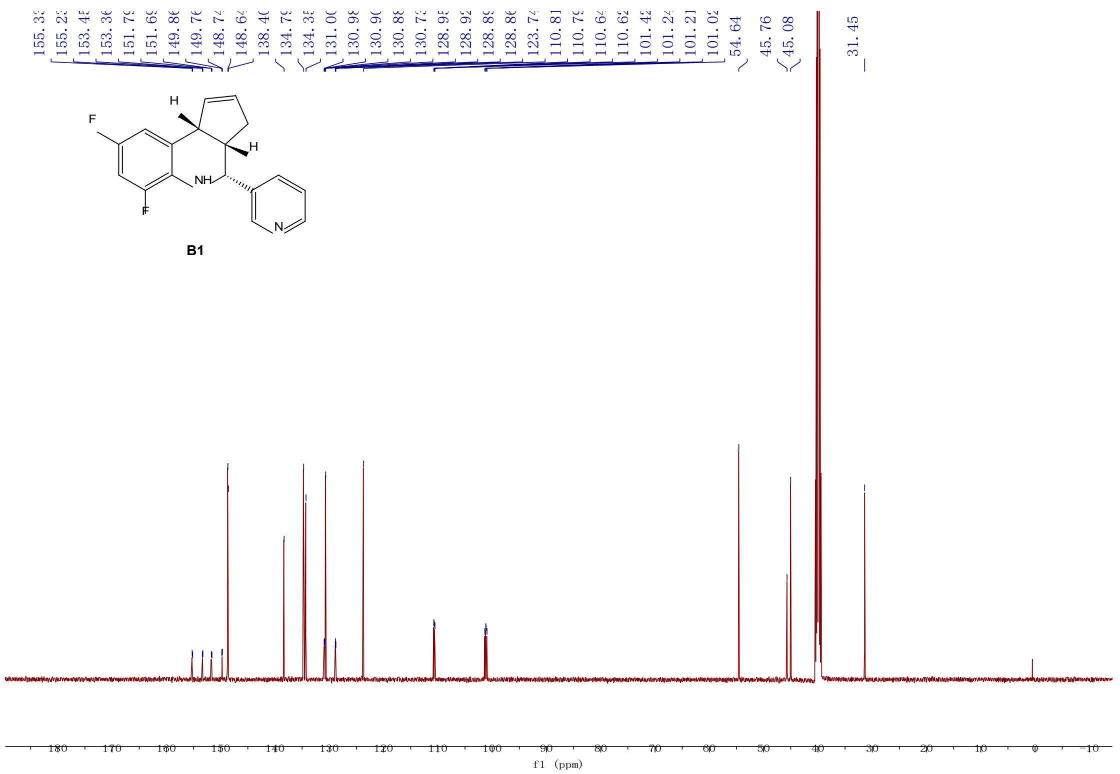

**[13]C NMR (126 MHz, DMSO-$d_6$)** δ154.34 (dd, J = 235.9, 11.8 Hz), 150.77 (dd, J = 242.6, 12.6 Hz), 148.74, 148.64, 138.40, 134.79, 134.35, 130.94 (dd, J = 12.9, 2.8 Hz), 130.73, 128.91 (dd, J = 7.7, 4.1 Hz), 123.74, 110.71 (dd, J = 21.2, 3.3 Hz), 101.22 (dd, J = 26.9, 23.0 Hz), 54.64, 45.76, 45.08, 31.45.

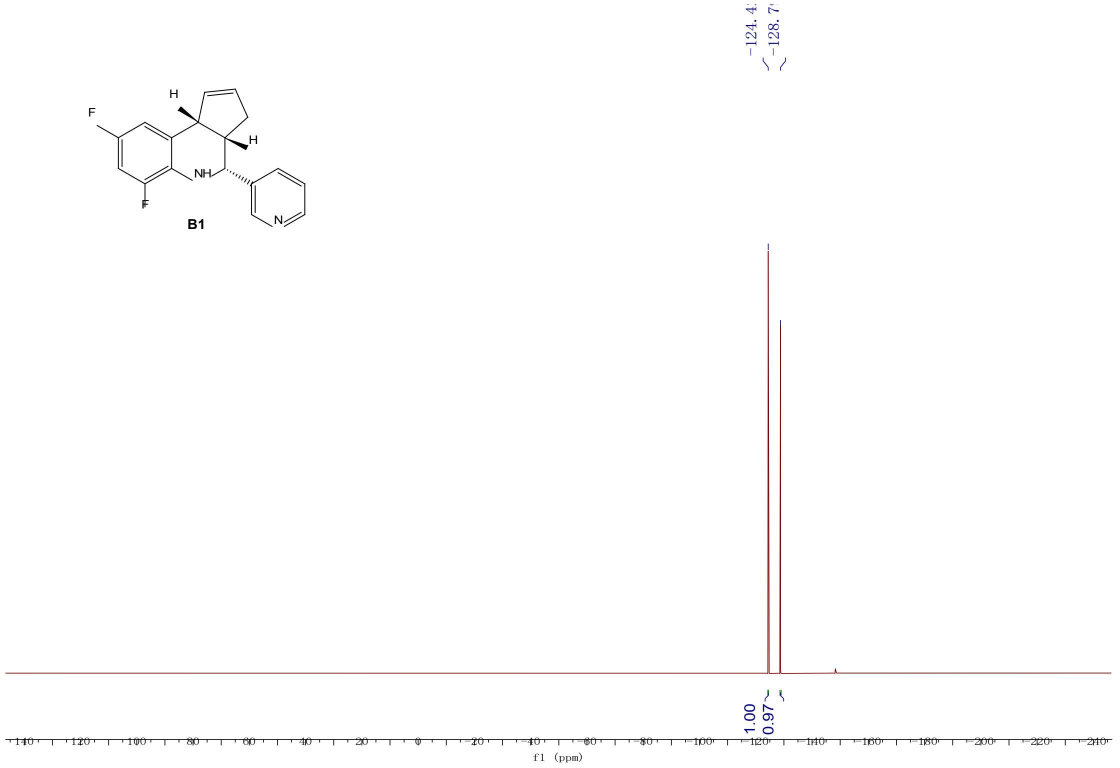

**[19]F NMR (471 MHz, DMSO-*d6*)** δ -124.42, -128.79.

5. X-Ray Structural Data for **B1**.

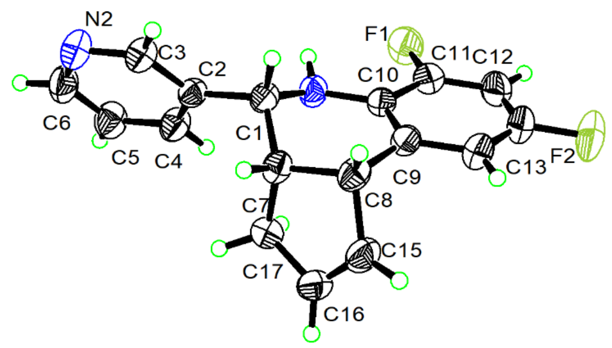

6. Crystal data and structure refinement for **B1.**

| | |
|---|---|
| Identification code | B1 |
| Empirical formula | $C_{17}H_{14}F_2N_2$ |
| Formula weight | 284.30 |
| Temperature/K | 301.00 |
| Crystal system | orthorhombic |
| Space group | $P2_12_12_1$ |
| a/Å | 5.5521(3) |
| b/Å | 13.8384(8) |
| c/Å | 17.9093(10) |
| α/° | 90 |
| β/° | 90 |
| γ/° | 90 |
| Volume/Å$^3$ | 1376.01(13) |
| Z | 4 |
| $\rho_{calc}$g/cm$^3$ | 1.372 |
| μ/mm$^{-1}$ | 0.837 |
| F(000) | 592.0 |
| Crystal size/mm$^3$ | 0.15 × 0.12 × 0.1 |
| Radiation | CuKα (λ = 1.54178) |
| 2Θ range for data collection/° | 8.074 to 137.028 |
| Index ranges | $-6 \leq h \leq 6$, $-16 \leq k \leq 16$, $-21 \leq l \leq 21$ |
| Reflections collected | 25250 |
| Independent reflections | 2535 [$R_{int}$ = 0.0357, $R_{sigma}$ = 0.0207] |
| Data/restraints/parameters | 2535/0/195 |
| Goodness-of-fit on F$^2$ | 1.048 |
| Final R indexes [I>=2σ (I)] | $R_1$ = 0.0256, $wR_2$ = 0.0686 |
| Final R indexes [all data] | $R_1$ = 0.0263, $wR_2$ = 0.0695 |
| Largest diff. peak/hole / e Å$^{-3}$ | 0.14/-0.11 |
| Flack parameter | 0.05(3) |

7. Fractional Atomic Coordinates (×$10^4$) and Equivalent Isotropic Displacement Parameters ($Å^2$×$10^3$) for **B1**. $U_{eq}$ is defined as 1/3 of the trace of the orthogonalised $U_{IJ}$ tensor.

| **Atom** | ***x*** | ***y*** | ***z*** | **U(eq)** |
|---|---|---|---|---|
| $F_1$ | 10420(2) | 1306.5(8) | 2654.9(6) | 58.3(3) |
| $F_2$ | 7263(4) | 1286.6(10) | 223.1(6) | 89.6(5) |
| $N_1$ | 7102(3) | 2595.0(10) | 3136.6(7) | 39.1(3) |
| $N_2$ | 3833(3) | 3708.3(12) | 5440.5(8) | 51.3(4) |
| $C_2$ | 5327(3) | 3449.6(11) | 4191.8(8) | 35.1(3) |
| $C_{10}$ | 7026(3) | 2265.5(11) | 2399.4(8) | 36.9(3) |
| $C_1$ | 4887(3) | 3018.5(11) | 3427.6(8) | 34.9(3) |
| $C_7$ | 3937(3) | 3772.4(11) | 2877.1(8) | 36.6(3) |
| $C_9$ | 5333(3) | 2596.1(11) | 1877.5(8) | 39.5(4) |
| $C_8$ | 3469(3) | 3346.6(12) | 2093.6(8) | 40.5(4) |
| $C_{17}$ | 5539(3) | 4658.3(11) | 2727.8(9) | 43.3(4) |
| $C_3$ | 3649(3) | 3330.4(13) | 4753.4(8) | 41.8(4) |
| $C_{11}$ | 8764(3) | 1612.4(12) | 2150.6(9) | 44.3(4) |
| $C_4$ | 7365(3) | 3983.1(13) | 4361.4(9) | 44.7(4) |
| $C_6$ | 5782(4) | 4244.1(13) | 5574.0(10) | 49.3(4) |
| $C_{16}$ | 4762(4) | 4945.8(13) | 1957.3(9) | 49.1(4) |
| $C_{14}$ | 5435(4) | 2252.0(14) | 1144.3(9) | 52.4(4) |
| $C_{15}$ | 3644(4) | 4243.3(13) | 1610.9(9) | 49.0(4) |
| $C_5$ | 7587(4) | 4393.4(14) | 5062.7(10) | 50.7(4) |
| $C_{12}$ | 8906(4) | 1278.2(14) | 1431.4(10) | 54.9(5) |
| $C_{13}$ | 7196(4) | 1609.4(15) | 940.9(9) | 58.0(5) |

8. Anisotropic Displacement Parameters ($Å^2$×$10^3$) for **B1**. The Anisotropic displacement factor exponent takes the form:
$-2\pi^2[h^2a^{*2}U_{11}+2hka^*b^*U_{12}+\ldots]$.

| **Atom** | **$U_{11}$** | **$U_{22}$** | **$U_{33}$** | **$U_{23}$** | **$U_{13}$** | **$U_{12}$** |
|---|---|---|---|---|---|---|
| $F_1$ | 57.6(6) | 58.9(6) | 58.3(6) | -1.1(5) | -0.4(5) | 17.6(5) |
| $F_2$ | 135.8(13) | 95.6(10) | 37.6(6) | -21.9(6) | 11.9(7) | 10.5(11) |
| $N_1$ | 44.4(7) | 41.6(7) | 31.3(6) | -1.6(5) | -3.4(6) | 5.6(6) |
| $N_2$ | 52.2(9) | 65.9(10) | 35.7(7) | -5.0(6) | 8.3(6) | -0.2(8) |
| $C_2$ | 38.7(7) | 36.8(7) | 29.7(7) | 2.4(6) | -0.4(6) | 2.6(6) |
| $C_{10}$ | 43.6(8) | 33.8(7) | 33.4(7) | -0.5(6) | 2.9(6) | -5.9(6) |
| $C_1$ | 36.2(8) | 38.4(7) | 30.2(7) | 0.7(6) | 0.7(6) | -4.0(6) |
| $C_7$ | 34.8(7) | 43.7(8) | 31.4(7) | -0.1(6) | 2.0(6) | 3.7(7) |
| $C_9$ | 47.8(8) | 39.1(8) | 31.5(7) | 0.1(6) | 1.4(7) | -7.2(7) |
| $C_8$ | 40.5(8) | 48.4(9) | 32.6(7) | 2.5(7) | -4.6(6) | -2.2(7) |
| $C_{17}$ | 50.6(9) | 36.6(7) | 42.8(8) | 0.6(6) | 0.9(7) | 2.2(7) |

8. Anisotropic Displacement Parameters ($Å^2×10^3$) for **B1**. The Anisotropic displacement factor exponent takes the form:
$-2\pi^2[h^2a^{*2}U_{11}+2hka^*b^*U_{12}+\ldots]$.

| **Atom** | **$U_{11}$** | **$U_{22}$** | **$U_{33}$** | **$U_{23}$** | **$U_{13}$** | **$U_{12}$** |
|---|---|---|---|---|---|---|
| $C_3$ | 39.7(8) | 50.5(9) | 35.2(8) | 0.2(7) | 2.6(7) | -1.7(7) |
| $C_{11}$ | 50.0(9) | 40.5(8) | 42.4(8) | 0.3(7) | 4.4(8) | -1.2(7) |
| $C_4$ | 43.5(9) | 54.3(9) | 36.3(8) | -1.1(7) | 4.1(7) | -7.7(8) |
| $C_6$ | 57.9(11) | 56.1(10) | 33.8(8) | -8.3(7) | -1.4(8) | 3.4(8) |
| $C_{16}$ | 62.8(11) | 40.5(8) | 44.1(9) | 9.0(7) | 7.3(8) | 8.2(8) |
| $C_{14}$ | 71.7(12) | 53.5(10) | 32.1(8) | -1.4(7) | -1.6(8) | -3.5(9) |
| $C_{15}$ | 59.1(10) | 53.6(10) | 34.3(8) | 7.4(7) | -2.6(8) | 11.0(9) |
| $C_5$ | 51.6(10) | 58.3(10) | 42.3(9) | -7.4(8) | -6.0(8) | -9.1(8) |
| $C_{12}$ | 69.5(12) | 46.1(9) | 49.3(10) | -7.4(8) | 18.4(9) | 0.1(9) |
| $C_{13}$ | 85.9(14) | 55.5(10) | 32.7(8) | -9.6(8) | 12.4(9) | -5.1(11) |

417

9. Bond Lengths for **B1.**

| **Atom** | **Atom** | **Length/Å** | **Atom** | **Atom** | **Length/Å** |
|---|---|---|---|---|---|
| $F_1$ | $C_{11}$ | 1.356(2) | $C_7$ | $C_8$ | 1.544(2) |
| $F_2$ | $C_{13}$ | 1.3616(19) | $C_7$ | $C_{17}$ | 1.538(2) |
| $N_1$ | $C_{10}$ | 1.3974(19) | $C_9$ | $C_8$ | 1.517(2) |
| $N_1$ | $C_1$ | 1.459(2) | $C_9$ | $C_{14}$ | 1.398(2) |
| $N_2$ | $C_3$ | 1.341(2) | $C_8$ | $C_{15}$ | 1.516(2) |
| $N_2$ | $C_6$ | 1.333(3) | $C_{17}$ | $C_{16}$ | 1.499(2) |
| $C_2$ | $C_1$ | 1.5128(19) | $C_{11}$ | $C_{12}$ | 1.371(2) |
| $C_2$ | $C_3$ | 1.381(2) | $C_4$ | $C_5$ | 1.384(2) |
| $C_2$ | $C_4$ | 1.385(2) | $C_6$ | $C_5$ | 1.373(3) |
| $C_{10}$ | $C_9$ | 1.402(2) | $C_{16}$ | $C_{15}$ | 1.310(3) |
| $C_{10}$ | $C_{11}$ | 1.395(2) | $C_{14}$ | $C_{13}$ | 1.371(3) |
| $C_1$ | $C_7$ | 1.529(2) | $C_{12}$ | $C_{13}$ | 1.372(3) |

418

10. Bond Angles for **B1.**

| **Atom** | **Atom** | **Atom** | **Angle/°** | **Atom** | **Atom** | **Atom** | **Angle/°** |
|---|---|---|---|---|---|---|---|
| $C_{10}$ | $N_1$ | $C_1$ | 116.32(13) | $C_{15}$ | $C_8$ | $C_7$ | 101.26(13) |
| $C_6$ | $N_2$ | $C_3$ | 116.30(15) | $C_{15}$ | $C_8$ | $C_9$ | 111.80(14) |
| $C_3$ | $C_2$ | $C_1$ | 120.20(14) | $C_{16}$ | $C_{17}$ | $C_7$ | 101.83(14) |
| $C_3$ | $C_2$ | $C_4$ | 117.08(14) | $N_2$ | $C_3$ | $C_2$ | 124.79(16) |
| $C_4$ | $C_2$ | $C_1$ | 122.72(14) | $F_1$ | $C_{11}$ | $C_{10}$ | 117.27(14) |
| $N_1$ | $C_{10}$ | $C_9$ | 122.91(14) | $F_1$ | $C_{11}$ | $C_{12}$ | 118.78(16) |
| $C_{11}$ | $C_{10}$ | $N_1$ | 119.51(15) | $C_{12}$ | $C_{11}$ | $C_{10}$ | 123.95(17) |
| $C_{11}$ | $C_{10}$ | $C_9$ | 117.52(14) | $C_5$ | $C_4$ | $C_2$ | 119.37(15) |
| $N_1$ | $C_1$ | $C_2$ | 110.22(12) | $N_2$ | $C_6$ | $C_5$ | 123.82(15) |

10. Bond Angles for **B1.**

| Atom | Atom | Atom | Angle/° | Atom | Atom | Atom | Angle/° |
|---|---|---|---|---|---|---|---|
| $N_1$ | $C_1$ | $C_7$ | 109.54(12) | $C_{15}$ | $C_{16}$ | $C_{17}$ | 112.05(15) |
| $C_2$ | $C_1$ | $C_7$ | 111.71(13) | $C_{13}$ | $C_{14}$ | $C_9$ | 119.95(18) |
| $C_1$ | $C_7$ | $C_8$ | 112.56(13) | $C_{16}$ | $C_{15}$ | $C_8$ | 111.58(14) |
| $C_1$ | $C_7$ | $C_{17}$ | 117.15(13) | $C_6$ | $C_5$ | $C_4$ | 118.58(17) |
| $C_{17}$ | $C_7$ | $C_8$ | 104.09(12) | $C_{11}$ | $C_{12}$ | $C_{13}$ | 116.68(18) |
| $C_{10}$ | $C_9$ | $C_8$ | 120.73(14) | $F_2$ | $C_{13}$ | $C_{14}$ | 118.91(19) |
| $C_{14}$ | $C_9$ | $C_{10}$ | 119.21(16) | $F_2$ | $C_{13}$ | $C_{12}$ | 118.41(19) |
| $C_{14}$ | $C_9$ | $C_8$ | 120.02(16) | $C_{14}$ | $C_{13}$ | $C_{12}$ | 122.69(16) |
| $C_9$ | $C_8$ | $C_7$ | 112.22(13) | | | | |

419

11. Torsion Angles for **B1.**

| A | B | C | D | Angle/° | A | B | C | D | Angle/° |
|---|---|---|---|---|---|---|---|---|---|
| $F_1$ | $C_{11}$ | $C_{12}$ | $C_{13}$ | -179.85(16) | $C_7$ | $C_{17}$ | $C_{16}$ | $C_{15}$ | -18.5(2) |
| $N_1$ | $C_{10}$ | $C_9$ | $C_8$ | 0.2(2) | $C_9$ | $C_{10}$ | $C_{11}$ | $F_1$ | 179.01(14) |
| $N_1$ | $C_{10}$ | $C_9$ | $C_{14}$ | 177.90(15) | $C_9$ | $C_{10}$ | $C_{11}$ | $C_{12}$ | -0.1(3) |
| $N_1$ | $C_{10}$ | $C_{11}$ | $F_1$ | 1.9(2) | $C_9$ | $C_8$ | $C_{15}$ | $C_{16}$ | -101.28(18) |
| $N_1$ | $C_{10}$ | $C_{11}$ | $C_{12}$ | -177.16(17) | $C_9$ | $C_{14}$ | $C_{13}$ | $F_2$ | -179.80(18) |
| $N_1$ | $C_1$ | $C_7$ | $C_8$ | 57.75(16) | $C_9$ | $C_{14}$ | $C_{13}$ | $C_{12}$ | 0.0(3) |
| $N_1$ | $C_1$ | $C_7$ | $C_{17}$ | -62.88(16) | $C_8$ | $C_7$ | $C_{17}$ | $C_{16}$ | 28.60(15) |
| $N_2$ | $C_6$ | $C_5$ | $C_4$ | 1.4(3) | $C_8$ | $C_9$ | $C_{14}$ | $C_{13}$ | 176.79(17) |
| $C_2$ | $C_1$ | $C_7$ | $C_8$ | -179.84(13) | $C_{17}$ | $C_7$ | $C_8$ | $C_9$ | 90.93(15) |
| $C_2$ | $C_1$ | $C_7$ | $C_{17}$ | 59.53(17) | $C_{17}$ | $C_7$ | $C_8$ | $C_{15}$ | -28.44(15) |
| $C_2$ | $C_4$ | $C_5$ | $C_6$ | 1.0(3) | $C_{17}$ | $C_{16}$ | $C_{15}$ | $C_8$ | 0.0(2) |
| $C_{10}$ | $N_1$ | $C_1$ | $C_2$ | -173.10(13) | $C_3$ | $N_2$ | $C_6$ | $C_5$ | -2.2(3) |
| $C_{10}$ | $N_1$ | $C_1$ | $C_7$ | -49.80(17) | $C_3$ | $C_2$ | $C_1$ | $N_1$ | -138.48(15) |
| $C_{10}$ | $C_9$ | $C_8$ | $C_7$ | 8.3(2) | $C_3$ | $C_2$ | $C_1$ | $C_7$ | 99.50(16) |
| $C_{10}$ | $C_9$ | $C_8$ | $C_{15}$ | 121.29(16) | $C_3$ | $C_2$ | $C_4$ | $C_5$ | -2.2(2) |
| $C_{10}$ | $C_9$ | $C_{14}$ | $C_{13}$ | -0.9(3) | $C_{11}$ | $C_{10}$ | $C_9$ | $C_8$ | -176.76(14) |
| $C_{10}$ | $C_{11}$ | $C_{12}$ | $C_{13}$ | -0.8(3) | $C_{11}$ | $C_{10}$ | $C_9$ | $C_{14}$ | 0.9(2) |
| $C_1$ | $N_1$ | $C_{10}$ | $C_9$ | 21.7(2) | $C_{11}$ | $C_{12}$ | $C_{13}$ | $F_2$ | -179.38(18) |
| $C_1$ | $N_1$ | $C_{10}$ | $C_{11}$ | -161.35(14) | $C_{11}$ | $C_{12}$ | $C_{13}$ | $C_{14}$ | 0.8(3) |
| $C_1$ | $C_2$ | $C_3$ | $N_2$ | -177.79(15) | $C_4$ | $C_2$ | $C_1$ | $N_1$ | 42.36(19) |
| $C_1$ | $C_2$ | $C_4$ | $C_5$ | 176.94(16) | $C_4$ | $C_2$ | $C_1$ | $C_7$ | -79.66(19) |
| $C_1$ | $C_7$ | $C_8$ | $C_9$ | -36.95(18) | $C_4$ | $C_2$ | $C_3$ | $N_2$ | 1.4(3) |
| $C_1$ | $C_7$ | $C_8$ | $C_{15}$ | -156.31(14) | $C_6$ | $N_2$ | $C_3$ | $C_2$ | 0.8(3) |
| $C_1$ | $C_7$ | $C_{17}$ | $C_{16}$ | 153.59(14) | $C_{14}$ | $C_9$ | $C_8$ | $C_7$ | -169.35(15) |
| $C_7$ | $C_8$ | $C_{15}$ | $C_{16}$ | 18.4(2) | $C_{14}$ | $C_9$ | $C_8$ | $C_{15}$ | -56.4(2) |

420

12. Hydrogen Atom Coordinates (Å×$10^4$) and Isotropic Displacement Parameters (Å$^2$×$10^3$) for **B1**.

| **Atom** | ***x*** | ***y*** | ***z*** | **U(eq)** |
|---|---|---|---|---|
| $H_{1A}$ | 3681.18 | 2505.04 | 3476.07 | 42 |
| $H_{7}$ | 2394.15 | 4008.53 | 3070.44 | 44 |
| $H_{8}$ | 1844.91 | 3071.2 | 2066.18 | 49 |
| $H_{17A}$ | 5233.66 | 5169.45 | 3086.1 | 52 |
| $H_{17B}$ | 7234.48 | 4489.38 | 2740.61 | 52 |
| $H_{3}$ | 2292.42 | 2960.87 | 4647.43 | 50 |
| $H_{4}$ | 8573.75 | 4064.71 | 4007.3 | 54 |
| $H_{6}$ | 5926.91 | 4533.51 | 6040.44 | 59 |
| $H_{16}$ | 5041.54 | 5551.65 | 1749.91 | 59 |
| $H_{14}$ | 4308.15 | 2458.88 | 794.79 | 63 |
| $H_{15}$ | 3033.08 | 4289.16 | 1128.61 | 59 |
| $H_{5}$ | 8930.37 | 4762.37 | 5185.17 | 61 |
| $H_{12}$ | 10101.41 | 848.46 | 1283.23 | 66 |
| $H_{1}$ | 7740(50) | 2187(16) | 3418(12) | 60(6) |

421

Data for **B2**

1. Chemical Structural for **B2**

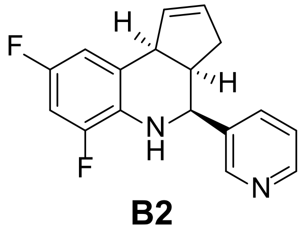

**B2**

2. HPLC trace of **B2**

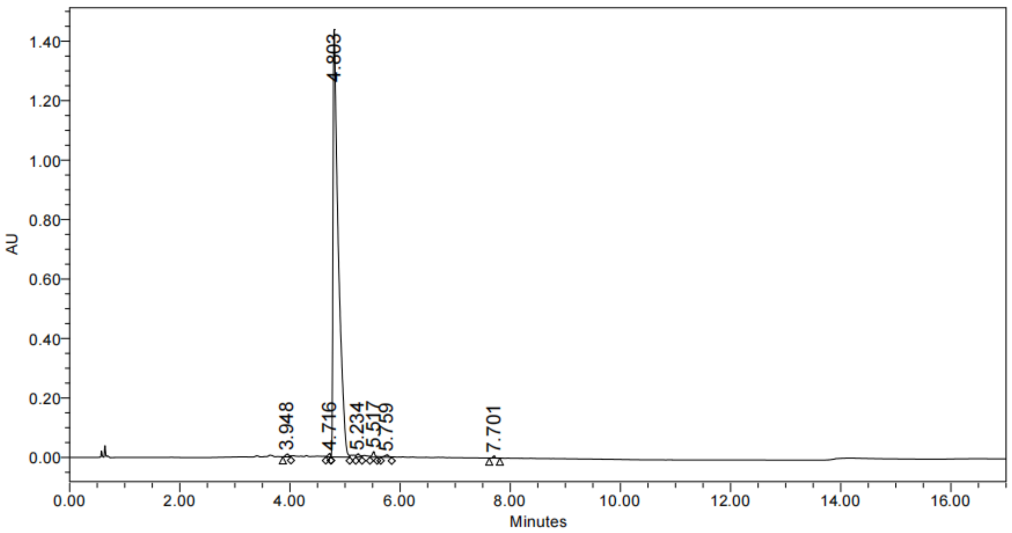

Peak Results

| | RT | Area | Height | % Area |
|---|---|---|---|---|
| 1 | 3.948 | 39410 | 8756 | 0.40 |
| 2 | 4.716 | 37264 | 10822 | 0.38 |
| 3 | 4.803 | 9526915 | 1438589 | 97.49 |
| 4 | 5.234 | 47650 | 10306 | 0.49 |
| 5 | 5.517 | 60932 | 17868 | 0.62 |
| 6 | 5.759 | 42423 | 7475 | 0.43 |
| 7 | 7.701 | 17673 | 6463 | 0.18 |

3. SFC analyst of **B2**

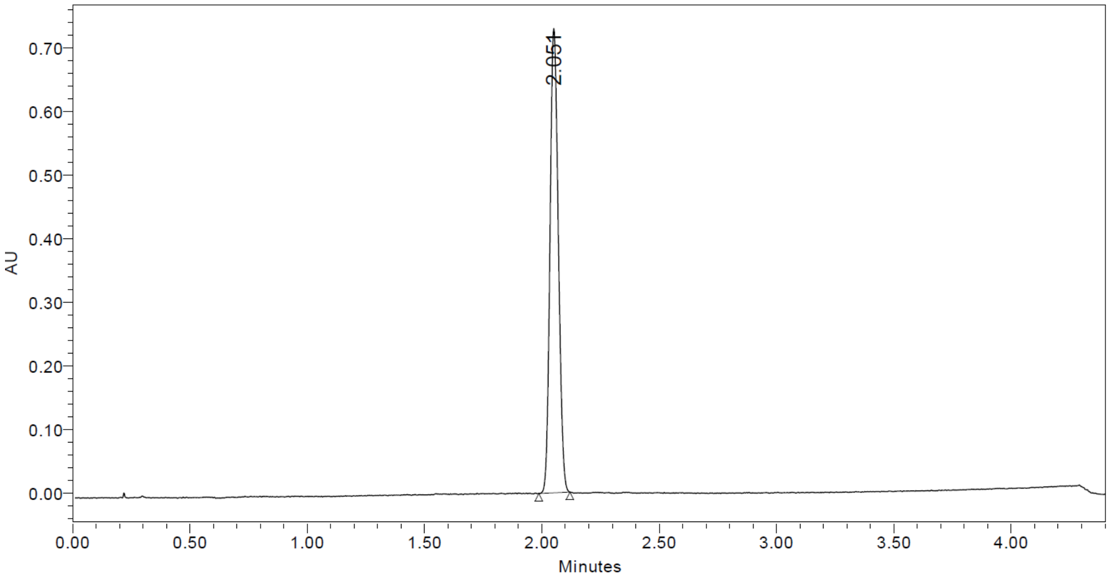

| | RT | Area | % Area |
|---|---|---|---|
| 1 | 2.051 | 1794689 | 100.00 |

4. HRMS of **B2**

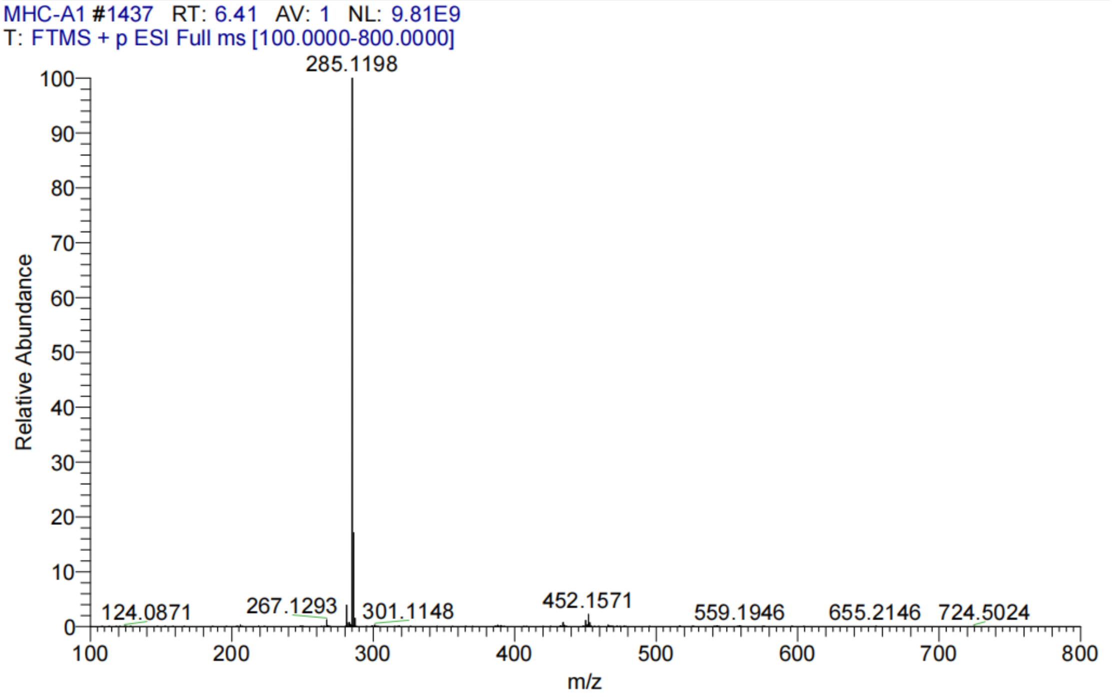

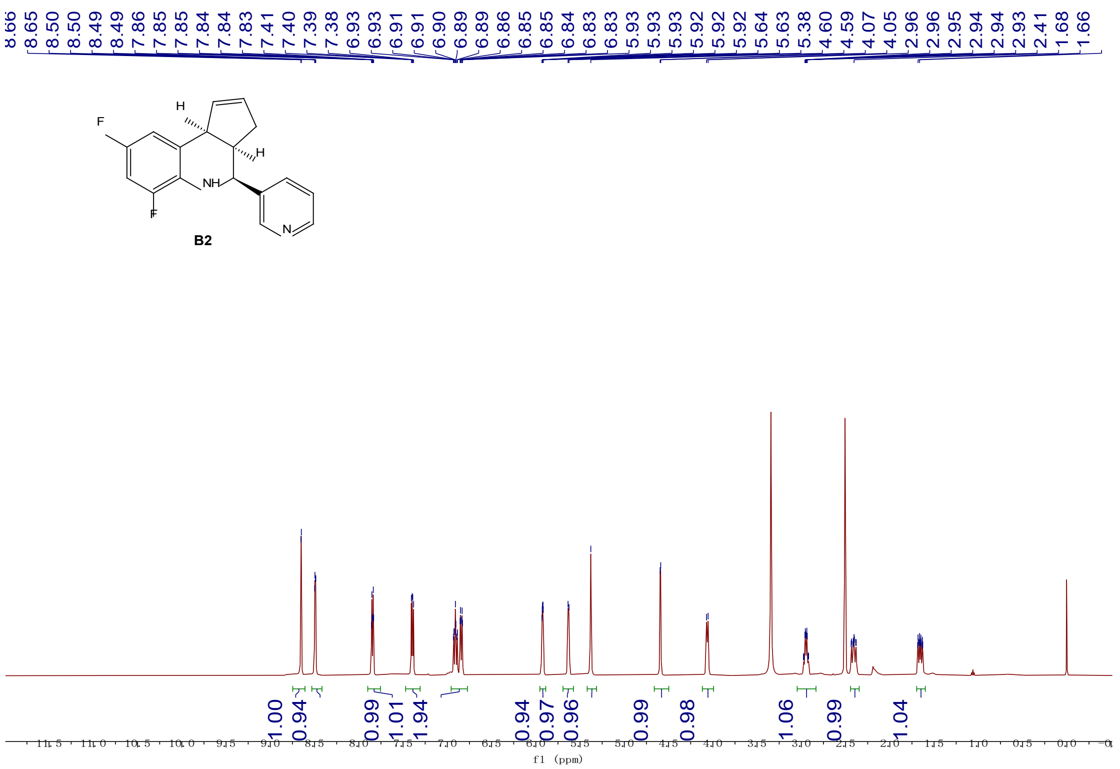

**[1]H NMR (500 MHz, DMSO-$d_6$)** δ 8.65 (d, *J* = 2.2 Hz, 1H), 8.49 (dd, *J* = 4.8, 1.6 Hz, 1H), 7.85 (dt, *J* = 7.9, 2.0 Hz, 1H), 7.39 (dd, *J* = 7.9, 4.8 Hz, 1H), 6.96 – 6.77 (m, 2H), 5.95 – 5.89 (m, 1H), 5.63 (d, *J* = 5.6 Hz, 1H), 5.38 (s, 1H), 4.59 (d, *J* = 3.3 Hz, 1H), 4.06 (d, *J* = 8.8 Hz, 1H), 2.95 (dddd, *J* = 9.9, 7.5, 5.9, 1.9 Hz, 1H), 2.44 – 2.35 (m, 1H), 1.66 (ddt, *J* = 16.1, 8.6, 2.2 Hz, 1H).

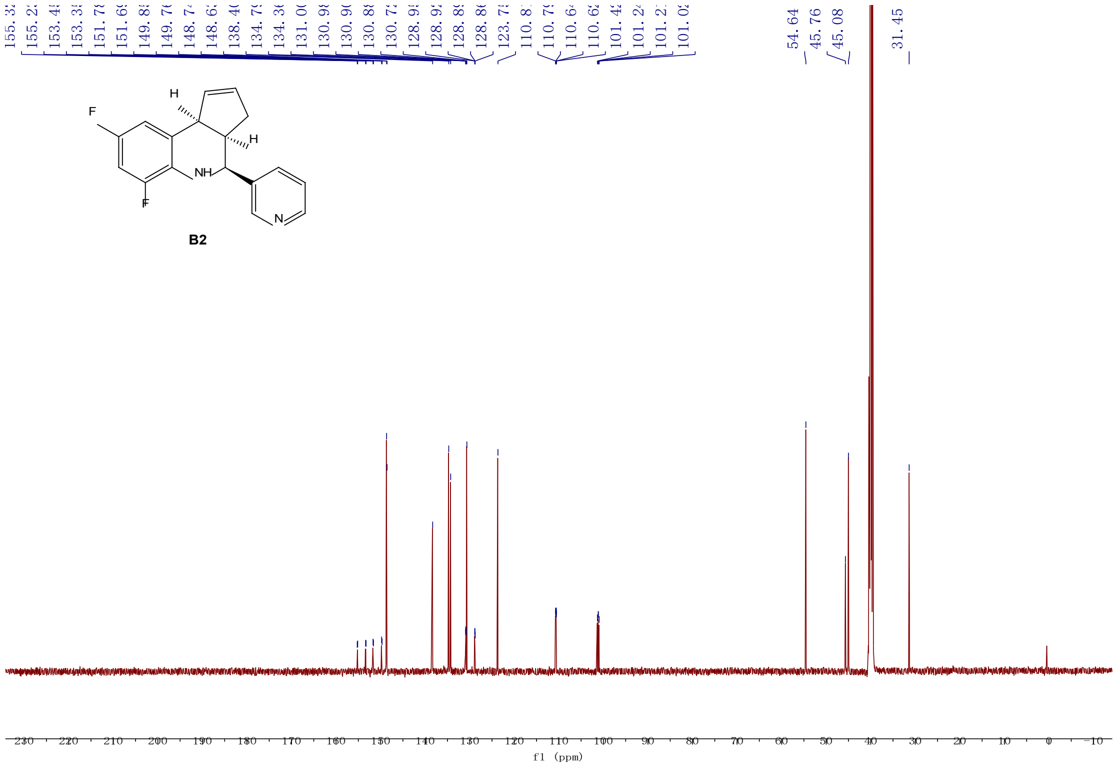

**[13]C NMR (126 MHz, DMSO-$d_6$)** δ 154.34 (dd, *J* = 236.0, 11.9 Hz), 150.77 (dd, *J* = 242.7, 12.4 Hz), 148.74, 148.63, 138.40, 134.79, 134.36, 130.94 (dd, *J* = 12.9, 2.7 Hz), 130.72, 128.91 (dd, *J*

= 7.8, 4.2 Hz), 123.75, 110.72 (dd, $J$ = 21.1, 3.3 Hz), 101.22 (dd, $J$ = 26.9, 23.0 Hz), 54.64, 45.76, 45.08, 31.45.

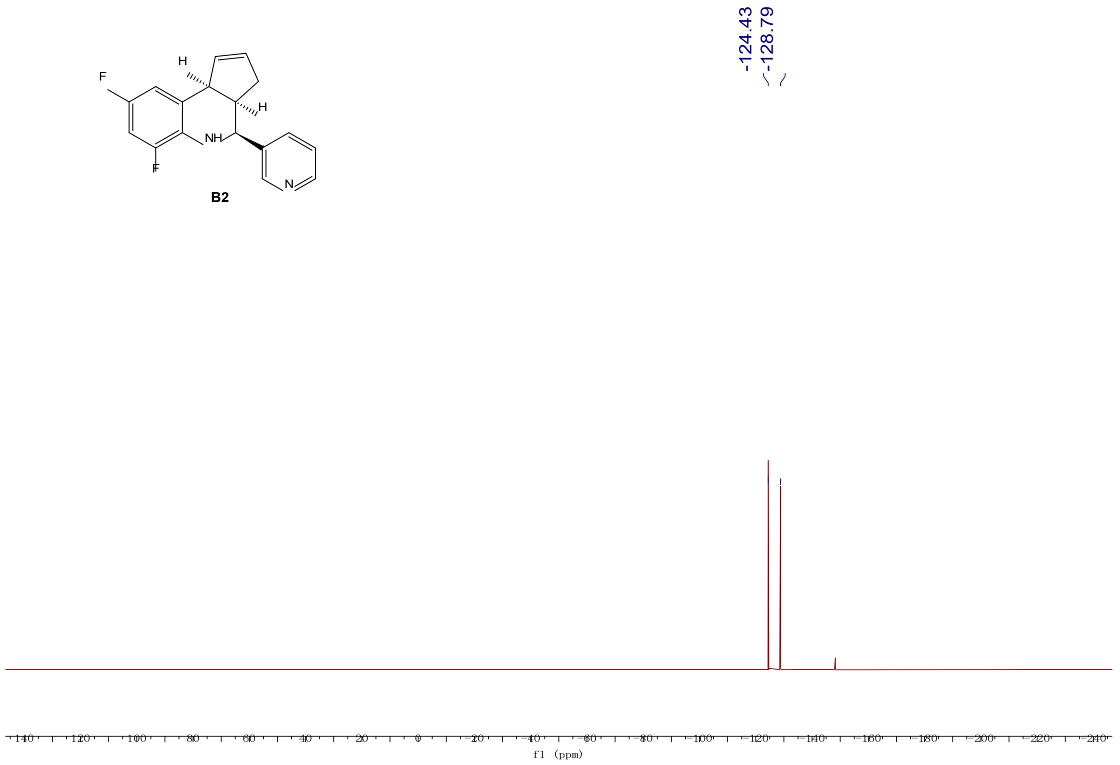

**$^{19}$F NMR (471 MHz, DMSO-$d_6$)** δ -124.43, -128.79.

5. X-Ray Structural Data for **B2**

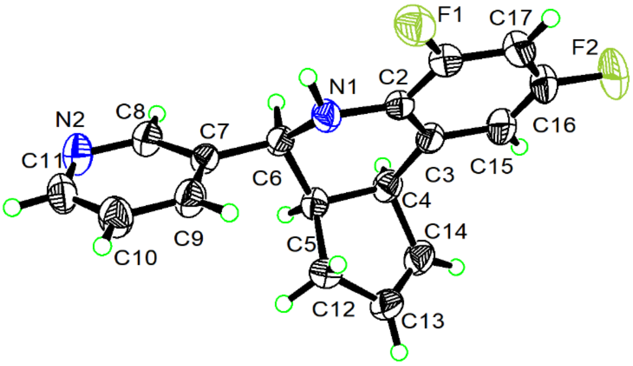

6. Crystal data and structure refinement for **B2.**

| | |
|---|---|
| Identification code | **B2** |
| Empirical formula | $C_{17}H_{14}F_2N_2$ |
| Formula weight | 284.30 |
| Temperature/K | 301.00 |
| Crystal system | orthorhombic |
| Space group | $P2_12_12_1$ |
| a/Å | 5.5511(2) |
| b/Å | 13.8305(4) |
| c/Å | 17.9177(5) |

| | |
|---|---|
| α/° | 90 |
| β/° | 90 |
| γ/° | 90 |
| Volume/Å$^3$ | 1375.62(7) |
| Z | 4 |
| $\rho_{calc}$g/cm$^3$ | 1.373 |
| μ/mm$^{-1}$ | 0.837 |
| F(000) | 592.0 |
| Crystal size/mm$^3$ | 0.15 × 0.12 × 0.1 |
| Radiation | CuKα (λ = 1.54178) |
| 2Θ range for data collection/° | 8.076 to 136.676 |
| Index ranges | -6 ≤ h ≤ 6, -16 ≤ k ≤ 16, -21 ≤ l ≤ 21 |
| Reflections collected | 25797 |
| Independent reflections | 2523 [$R_{int}$ = 0.0321, $R_{sigma}$ = 0.0199] |
| Data/restraints/parameters | 2523/0/190 |
| Goodness-of-fit on F$^2$ | 1.084 |
| Final R indexes [I>=2σ (I)] | $R_1$ = 0.0303, w$R_2$ = 0.0758 |
| Final R indexes [all data] | $R_1$ = 0.0308, w$R_2$ = 0.0764 |
| Largest diff. peak/hole / e Å$^{-3}$ | 0.12/-0.19 |
| Flack parameter | 0.05(3) |

7. Fractional Atomic Coordinates (×10$^4$) and Equivalent Isotropic Displacement Parameters (Å$^2$×10$^3$) for **B2**. $U_{eq}$ is defined as 1/3 of the trace of the orthogonalised $U_{IJ}$ tensor.

| **Atom** | ***x*** | ***y*** | ***z*** | **U(eq)** |
|---|---|---|---|---|
| $F_1$ | 10421(2) | 1306.0(8) | 7345.2(6) | 54.2(3) |
| $F_2$ | 7266(4) | 1286.8(11) | 9776.9(6) | 85.2(5) |
| $N_1$ | 7103(3) | 2593.5(10) | 6863.3(7) | 35.4(3) |
| $N_2$ | 3831(3) | 3708.6(13) | 4559.7(8) | 47.8(4) |
| $C_7$ | 5325(3) | 3448.0(12) | 5808.2(8) | 31.6(3) |
| $C_6$ | 4883(3) | 3017.6(12) | 6572.0(8) | 31.6(3) |
| $C_2$ | 7028(3) | 2265.3(11) | 7600.2(9) | 33.5(3) |
| $C_5$ | 3937(3) | 3772.2(12) | 7123.2(8) | 32.8(3) |
| $C_3$ | 5336(3) | 2596.1(12) | 8122.9(9) | 36.0(4) |
| $C_4$ | 3467(3) | 3345.6(13) | 7906.5(9) | 36.8(4) |
| $C_8$ | 3647(3) | 3330.1(14) | 5246.2(9) | 38.5(4) |
| $C_{12}$ | 5540(3) | 4658.1(12) | 7272.5(10) | 39.6(4) |
| $C_9$ | 7369(3) | 3983.4(14) | 5639.1(9) | 41.1(4) |
| $C_1$ | 8765(3) | 1612.4(13) | 7850.4(10) | 40.8(4) |
| $C_{11}$ | 5779(4) | 4245.1(14) | 4426.2(10) | 45.5(4) |
| $C_{14}$ | 3643(4) | 4244.2(14) | 8389.4(10) | 45.3(4) |

7. Fractional Atomic Coordinates ($\times 10^4$) and Equivalent Isotropic Displacement Parameters ($Å^2\times 10^3$) for **B2**. $U_{eq}$ is defined as 1/3 of the trace of the orthogonalised $U_{IJ}$ tensor.

| **Atom** | ***x*** | ***y*** | ***z*** | **U(eq)** |
|---|---|---|---|---|
| $C_{13}$ | 4761(4) | 4946.7(14) | 8042.0(10) | 45.7(4) |
| $C_{15}$ | 5437(4) | 2252.1(15) | 8856.2(10) | 48.7(5) |
| $C_{10}$ | 7587(4) | 4393.3(15) | 4937.8(10) | 46.8(4) |
| $C_{17}$ | 8910(4) | 1277.5(15) | 8568.0(11) | 51.2(5) |
| $C_{16}$ | 7198(5) | 1609.7(16) | 9058.6(10) | 54.4(5) |

8. Anisotropic Displacement Parameters ($Å^2\times 10^3$) for **B2**. The Anisotropic displacement factor exponent takes the form:
$-2\pi^2[h^2a^{*2}U_{11}+2hka^*b^*U_{12}+\ldots]$.

| **Atom** | $\mathbf{U_{11}}$ | $\mathbf{U_{22}}$ | $\mathbf{U_{33}}$ | $\mathbf{U_{23}}$ | $\mathbf{U_{13}}$ | $\mathbf{U_{12}}$ |
|---|---|---|---|---|---|---|
| $F_1$ | 54.0(6) | 53.8(6) | 54.7(7) | 1.0(5) | 0.8(5) | 17.0(5) |
| $F_2$ | 132.4(14) | 88.9(10) | 34.3(6) | 21.6(6) | -11.5(7) | 10.4(11) |
| $N_1$ | 41.2(7) | 36.5(7) | 28.5(7) | 1.5(5) | 3.0(6) | 5.4(6) |
| $N_2$ | 48.6(9) | 61.9(10) | 32.8(7) | 5.1(7) | -8.5(7) | 0.1(8) |
| $C_7$ | 35.5(7) | 32.7(7) | 26.6(7) | -2.4(6) | 0.4(6) | 2.1(6) |
| $C_6$ | 33.7(8) | 33.6(8) | 27.4(7) | -1.0(6) | -0.8(6) | -3.7(6) |
| $C_2$ | 40.5(8) | 29.4(7) | 30.7(7) | 0.3(6) | -2.7(6) | -5.4(6) |
| $C_5$ | 31.3(7) | 38.9(8) | 28.2(7) | -0.6(6) | -1.6(6) | 3.1(7) |
| $C_3$ | 45.0(9) | 34.6(8) | 28.4(7) | -0.1(6) | -1.5(7) | -6.8(7) |
| $C_4$ | 38.2(8) | 42.7(9) | 29.5(8) | -2.9(7) | 4.7(7) | -1.5(7) |
| $C_8$ | 37.3(8) | 45.5(9) | 32.6(8) | -0.7(7) | -2.2(7) | -1.9(8) |
| $C_{12}$ | 47.5(9) | 31.8(8) | 39.4(9) | -0.4(7) | -1.4(8) | 2.0(7) |
| $C_9$ | 39.6(9) | 50.3(10) | 33.5(8) | 0.9(7) | -4.1(7) | -7.2(8) |
| $C_1$ | 47.4(9) | 35.5(8) | 39.4(9) | -0.2(7) | -4.3(8) | -1.3(8) |
| $C_{11}$ | 53.1(11) | 52.1(11) | 31.2(8) | 7.9(8) | 1.3(8) | 3.2(9) |
| $C_{14}$ | 56.3(10) | 48.7(10) | 30.8(8) | -7.4(7) | 2.7(8) | 10.0(9) |
| $C_{13}$ | 60.0(11) | 36.1(9) | 40.8(9) | -8.8(8) | -7.2(9) | 8.3(8) |
| $C_{15}$ | 68.2(12) | 49.0(10) | 28.9(8) | 1.8(7) | 2.0(8) | -3.1(10) |
| $C_{10}$ | 47.6(10) | 53.9(11) | 38.8(9) | 6.9(8) | 5.6(8) | -9.0(9) |
| $C_{17}$ | 65.9(12) | 41.5(10) | 46.1(10) | 7.0(8) | -17.6(9) | 0.6(10) |
| $C_{16}$ | 81.9(14) | 50.9(10) | 30.4(9) | 9.6(8) | -12.2(10) | -5.7(11) |

9. Bond Lengths for **B2.**

| **Atom** | **Atom** | **Length/Å** | **Atom** | **Atom** | **Length/Å** |
|---|---|---|---|---|---|
| $F_1$ | $C_1$ | 1.358(2) | $C_5$ | $C_4$ | 1.545(2) |
| $F_2$ | $C_{16}$ | 1.363(2) | $C_5$ | $C_{12}$ | 1.538(2) |
| $N_1$ | $C_6$ | 1.461(2) | $C_3$ | $C_4$ | 1.517(3) |
| $N_1$ | $C_2$ | 1.397(2) | $C_3$ | $C_{15}$ | 1.398(2) |

9. Bond Lengths for **B2.**

| Atom | Atom | Length/Å | Atom | Atom | Length/Å |
|---|---|---|---|---|---|
| $N_2$ | $C_8$ | 1.341(2) | $C_4$ | $C_{14}$ | 1.517(2) |
| $N_2$ | $C_{11}$ | 1.333(3) | $C_{12}$ | $C_{13}$ | 1.499(2) |
| $C_7$ | $C_6$ | 1.512(2) | $C_9$ | $C_{10}$ | 1.384(2) |
| $C_7$ | $C_8$ | 1.381(2) | $C_1$ | $C_{17}$ | 1.369(3) |
| $C_7$ | $C_9$ | 1.388(2) | $C_{11}$ | $C_{10}$ | 1.375(3) |
| $C_6$ | $C_5$ | 1.530(2) | $C_{14}$ | $C_{13}$ | 1.310(3) |
| $C_2$ | $C_3$ | 1.403(2) | $C_{15}$ | $C_{16}$ | 1.370(3) |
| $C_2$ | $C_1$ | 1.395(2) | $C_{17}$ | $C_{16}$ | 1.374(3) |

451

10. Bond Angles for **B2.**

| Atom | Atom | Atom | Angle/° | Atom | Atom | Atom | Angle/° |
|---|---|---|---|---|---|---|---|
| $C_2$ | $N_1$ | $C_6$ | 116.27(13) | $C_3$ | $C_4$ | $C_{14}$ | 111.71(14) |
| $C_{11}$ | $N_2$ | $C_8$ | 116.34(15) | $C_{14}$ | $C_4$ | $C_5$ | 101.20(14) |
| $C_8$ | $C_7$ | $C_6$ | 120.25(15) | $N_2$ | $C_8$ | $C_7$ | 124.84(17) |
| $C_8$ | $C_7$ | $C_9$ | 117.07(14) | $C_{13}$ | $C_{12}$ | $C_5$ | 101.85(14) |
| $C_9$ | $C_7$ | $C_6$ | 122.67(14) | $C_{10}$ | $C_9$ | $C_7$ | 119.21(16) |
| $N_1$ | $C_6$ | $C_7$ | 110.15(13) | $F_1$ | $C_1$ | $C_2$ | 117.12(15) |
| $N_1$ | $C_6$ | $C_5$ | 109.44(12) | $F_1$ | $C_1$ | $C_{17}$ | 118.72(17) |
| $C_7$ | $C_6$ | $C_5$ | 111.80(13) | $C_{17}$ | $C_1$ | $C_2$ | 124.16(18) |
| $N_1$ | $C_2$ | $C_3$ | 123.04(15) | $N_2$ | $C_{11}$ | $C_{10}$ | 123.75(16) |
| $C_1$ | $C_2$ | $N_1$ | 119.55(15) | $C_{13}$ | $C_{14}$ | $C_4$ | 111.54(15) |
| $C_1$ | $C_2$ | $C_3$ | 117.34(15) | $C_{14}$ | $C_{13}$ | $C_{12}$ | 112.09(16) |
| $C_6$ | $C_5$ | $C_4$ | 112.58(13) | $C_{16}$ | $C_{15}$ | $C_3$ | 119.87(19) |
| $C_6$ | $C_5$ | $C_{12}$ | 117.21(13) | $C_{11}$ | $C_{10}$ | $C_9$ | 118.73(18) |
| $C_{12}$ | $C_5$ | $C_4$ | 104.12(13) | $C_1$ | $C_{17}$ | $C_{16}$ | 116.56(19) |
| $C_2$ | $C_3$ | $C_4$ | 120.66(14) | $F_2$ | $C_{16}$ | $C_{15}$ | 118.8(2) |
| $C_{15}$ | $C_3$ | $C_2$ | 119.30(17) | $F_2$ | $C_{16}$ | $C_{17}$ | 118.4(2) |
| $C_{15}$ | $C_3$ | $C_4$ | 120.01(16) | $C_{15}$ | $C_{16}$ | $C_{17}$ | 122.77(17) |
| $C_3$ | $C_4$ | $C_5$ | 112.20(13) | | | | |

452

11. Torsion Angles for **B2.**

| A | B | C | D | Angle/° | A | B | C | D | Angle/° |
|---|---|---|---|---|---|---|---|---|---|
| $F_1$ | $C_1$ | $C_{17}$ | $C_{16}$ | 179.85(17) | $C_5$ | $C_{12}$ | $C_{13}$ | $C_{14}$ | 18.6(2) |
| $N_1$ | $C_6$ | $C_5$ | $C_4$ | -57.86(17) | $C_3$ | $C_2$ | $C_1$ | $F_1$ | -178.99(15) |
| $N_1$ | $C_6$ | $C_5$ | $C_{12}$ | 62.87(17) | $C_3$ | $C_2$ | $C_1$ | $C_{17}$ | 0.2(3) |
| $N_1$ | $C_2$ | $C_3$ | $C_4$ | -0.1(2) | $C_3$ | $C_4$ | $C_{14}$ | $C_{13}$ | 101.25(19) |
| $N_1$ | $C_2$ | $C_3$ | $C_{15}$ | -177.94(16) | $C_3$ | $C_{15}$ | $C_{16}$ | $F_2$ | 179.79(19) |
| $N_1$ | $C_2$ | $C_1$ | $F_1$ | -1.9(2) | $C_3$ | $C_{15}$ | $C_{16}$ | $C_{17}$ | 0.0(3) |

11. Torsion Angles for **B2.**

| A | B | C | D | Angle/° | A | B | C | D | Angle/° |
|---|---|---|---|---|---|---|---|---|---|
| $N_1$ | $C_2$ | $C_1$ | $C_{17}$ | 177.22(18) | $C_4$ | $C_5$ | $C_{12}$ | $C_{13}$ | -28.66(16) |
| $N_2$ | $C_{11}$ | $C_{10}$ | $C_9$ | -1.5(3) | $C_4$ | $C_3$ | $C_{15}$ | $C_{16}$ | -176.91(18) |
| $C_7$ | $C_6$ | $C_5$ | $C_4$ | 179.83(13) | $C_4$ | $C_{14}$ | $C_{13}$ | $C_{12}$ | -0.1(2) |
| $C_7$ | $C_6$ | $C_5$ | $C_{12}$ | -59.44(18) | $C_8$ | $N_2$ | $C_{11}$ | $C_{10}$ | 2.3(3) |
| $C_7$ | $C_9$ | $C_{10}$ | $C_{11}$ | -0.9(3) | $C_8$ | $C_7$ | $C_6$ | $N_1$ | 138.59(16) |
| $C_6$ | $N_1$ | $C_2$ | $C_3$ | -21.7(2) | $C_8$ | $C_7$ | $C_6$ | $C_5$ | -99.51(17) |
| $C_6$ | $N_1$ | $C_2$ | $C_1$ | 161.39(15) | $C_8$ | $C_7$ | $C_9$ | $C_{10}$ | 2.1(3) |
| $C_6$ | $C_7$ | $C_8$ | $N_2$ | 177.71(16) | $C_{12}$ | $C_5$ | $C_4$ | $C_3$ | -90.78(16) |
| $C_6$ | $C_7$ | $C_9$ | $C_{10}$ | -176.86(17) | $C_{12}$ | $C_5$ | $C_4$ | $C_{14}$ | 28.43(16) |
| $C_6$ | $C_5$ | $C_4$ | $C_3$ | 37.20(19) | $C_9$ | $C_7$ | $C_6$ | $N_1$ | -42.4(2) |
| $C_6$ | $C_5$ | $C_4$ | $C_{14}$ | 156.41(14) | $C_9$ | $C_7$ | $C_6$ | $C_5$ | 79.5(2) |
| $C_6$ | $C_5$ | $C_{12}$ | $C_{13}$ | -153.73(14) | $C_9$ | $C_7$ | $C_8$ | $N_2$ | -1.3(3) |
| $C_2$ | $N_1$ | $C_6$ | $C_7$ | 173.06(13) | $C_1$ | $C_2$ | $C_3$ | $C_4$ | 176.84(15) |
| $C_2$ | $N_1$ | $C_6$ | $C_5$ | 49.76(17) | $C_1$ | $C_2$ | $C_3$ | $C_{15}$ | -1.0(2) |
| $C_2$ | $C_3$ | $C_4$ | $C_5$ | -8.5(2) | $C_1$ | $C_{17}$ | $C_{16}$ | $F_2$ | 179.40(19) |
| $C_2$ | $C_3$ | $C_4$ | $C_{14}$ | -121.36(17) | $C_1$ | $C_{17}$ | $C_{16}$ | $C_{15}$ | -0.8(3) |
| $C_2$ | $C_3$ | $C_{15}$ | $C_{16}$ | 0.9(3) | $C_{11}$ | $N_2$ | $C_8$ | $C_7$ | -0.9(3) |
| $C_2$ | $C_1$ | $C_{17}$ | $C_{16}$ | 0.7(3) | $C_{15}$ | $C_3$ | $C_4$ | $C_5$ | 169.30(16) |
| $C_5$ | $C_4$ | $C_{14}$ | $C_{13}$ | -18.3(2) | $C_{15}$ | $C_3$ | $C_4$ | $C_{14}$ | 56.5(2) |

12. Hydrogen Atom Coordinates ($Å\times10^4$) and Isotropic Displacement Parameters ($Å^2\times10^3$) for **B2.**

| **Atom** | ***x*** | ***y*** | ***z*** | **U(eq)** |
|---|---|---|---|---|
| $H_1$ | 7666.21 | 2157.67 | 6568.76 | 43 |
| $H_6$ | 3677.12 | 2503.97 | 6523.99 | 38 |
| $H_5$ | 2394.39 | 4009.15 | 6930.32 | 39 |
| $H_4$ | 1843.73 | 3069.52 | 7933.98 | 44 |
| $H_8$ | 2288.83 | 2960.96 | 5351.94 | 46 |
| $H_{12A}$ | 7235.16 | 4488.74 | 7260.26 | 48 |
| $H_{12B}$ | 5235.93 | 5169.29 | 6914.01 | 48 |
| $H_9$ | 8577.89 | 4065.49 | 5992.99 | 49 |
| $H_{11}$ | 5923.29 | 4535.77 | 3960.29 | 55 |
| $H_{14}$ | 3034.14 | 4290.46 | 8871.53 | 54 |
| $H_{13}$ | 5039.18 | 5553.41 | 8248.65 | 55 |
| $H_{15}$ | 4310.74 | 2459.03 | 9205.7 | 58 |
| $H_{10}$ | 8931.04 | 4761.91 | 4815 | 56 |
| $H_{17}$ | 10106.22 | 847.29 | 8715.69 | 61 |

#### 2. Supplementary Figures

##### Supplementary Figure 1

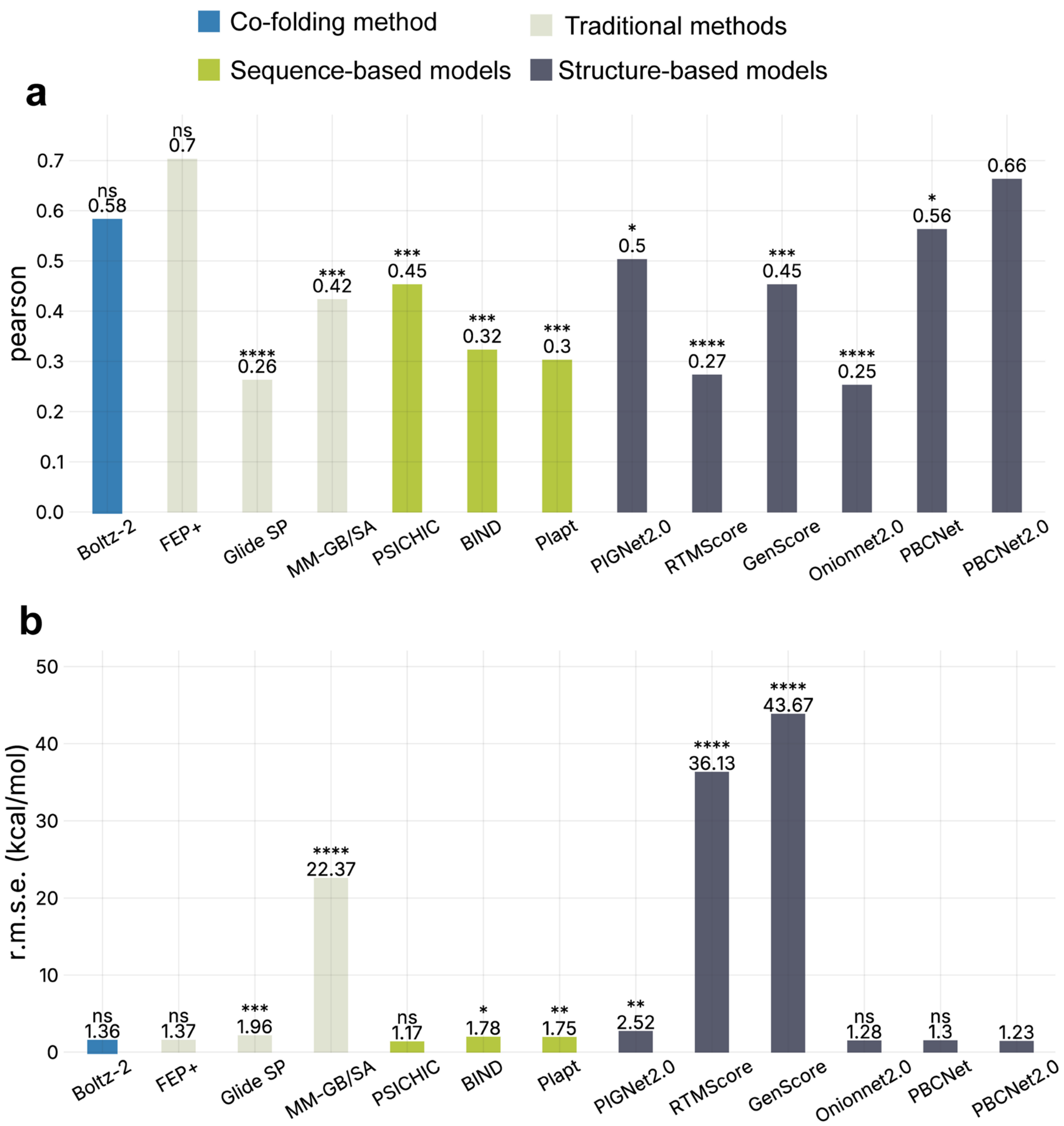

**Supplementary Figure 1.** The results of direct prediction one FEP set. The x-axis represents various binding affinity prediction methods, while the y-axis denotes the average of their performance (**a.** ranking ability, **b.** prediction ability) on the FEP set. The two-sided Wilcoxon signed-rank test was used to analyze whether there were significant differences between each model and the PBCNet2.0 predictions. *p ≤ 0.05, **p ≤ 0.01, ***p ≤ 0.001, ****p ≤ 0.0001, and ns (not significant) for p > 0.05. Different types of models are distinguished using bars of different colors.

**Supplementary Figure 2**

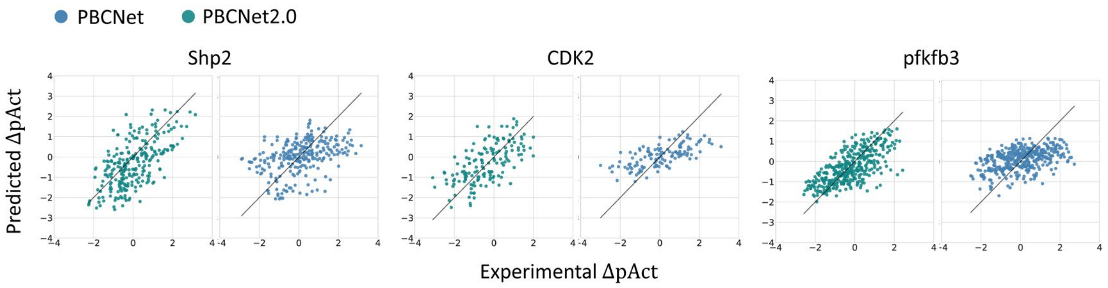

**Supplementary Figure 2.** Visualization of PBCNet2.0 and PBCNet prediction results. The x-axis represents experimental ΔpAct values, while the y-axis represents the predicted values. Green points represent PBCNet2.0 predictions, and blue points show PBCNet predictions.

**Supplementary Figure 3**

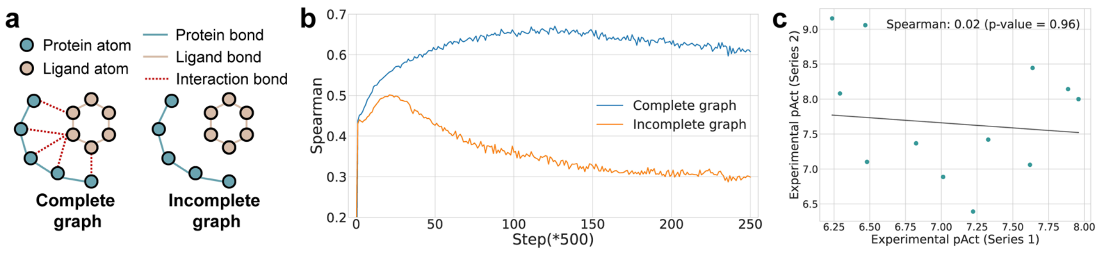

**Supplementary Figure 3. a. Illustration of Complete and Incomplete graphs.** An Incomplete graph is derived by removing interaction bonds between ligand and protein atoms from a Complete graph. **b. Performance dynamics of PBCNet2.0 on Complete and Incomplete graphs.** The x-axis represents the number of training steps and the y-axis indicates the ranking performance (Spearman) on the FEP set. The blue curve corresponds to the Complete graphs and the orange curve to the Incomplete graphs. **c. Visualization of activity correlation for a chemical series pair from the SAR-Diff Set.** Each scatter point represents the binding affinities of the compounds with the same R-group. P value indicates the significance of the Spearman coefficient in an exact permutation test.

**Supplementary Figure 4**

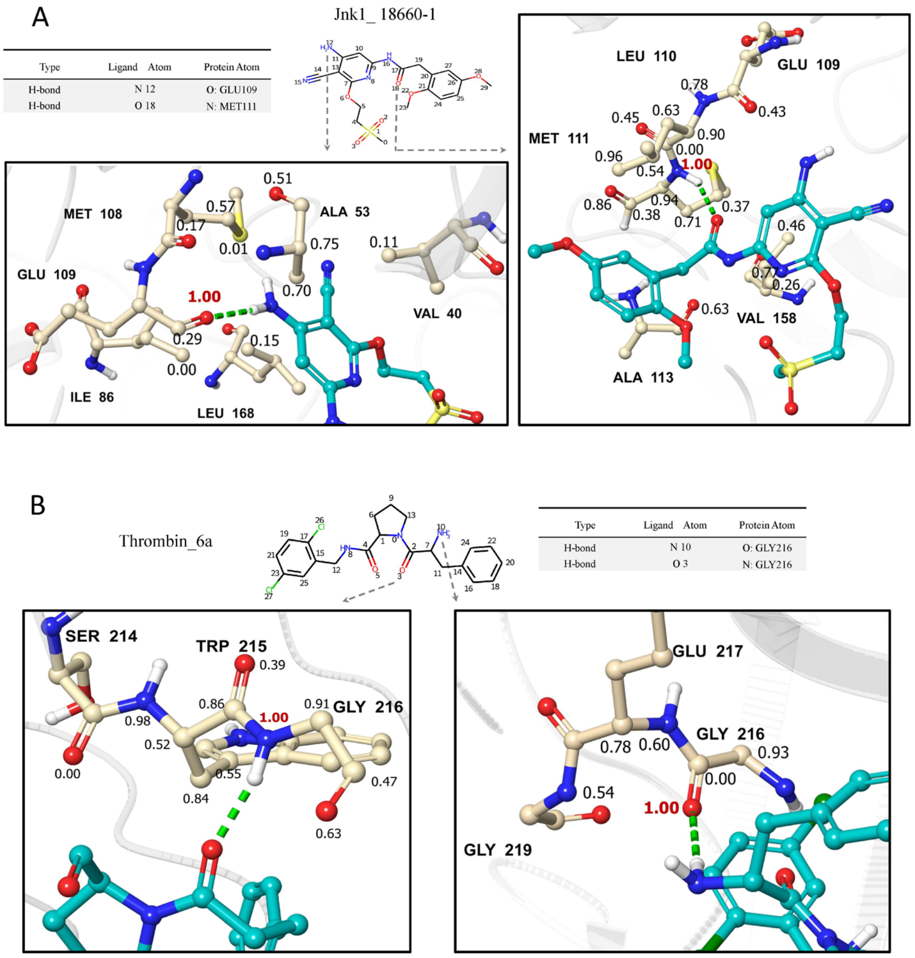

**Supplementary Figure 4.** Interpretability analysis results of PBCNet2.0 on two ligands: **A.** a JNK1 inhibitor 18660-1 and **B.** a Thrombin inhibitor 6a. The molecular structure, hydrogen-bond calculation results, 3-dimensional hydrogen-bonds visualization graphs, and weights visualization graphs are shown for comparison. In each visualization graph, the hydrogen bond is depicted as green dashed lines, and protein atoms within 5 Å of the nitrogen atom were annotated with weights, where red number indicates the weights associated with the oxygen atom involved in the hydrogen bond.

**Supplementary Figure 5**

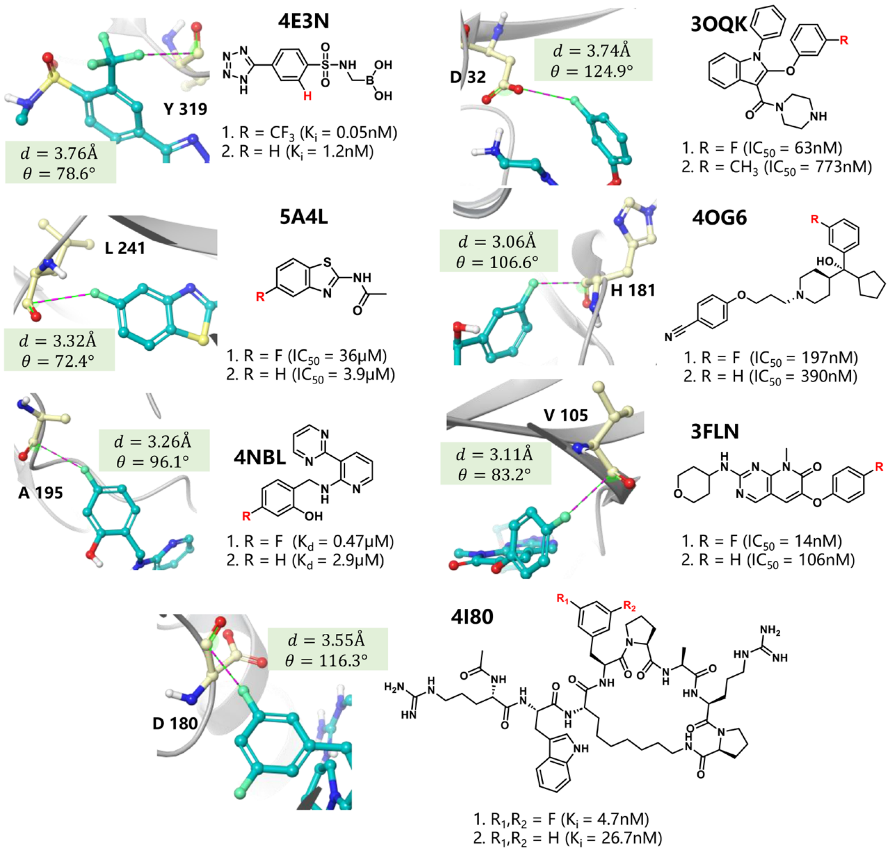

**Supplementary Figure 5.** Structures and binding affinities of protein-ligand complexes in the F-Opt test set. Structural details of seven additional protein-ligand complexes are shown, including interaction distances (d) and angles (θ) between protein residues and fluorinated ligands. Key interacting residues are labeled (e.g., Y319). For each ligand, two variants are shown: fluorinated (R = F) and non-fluorinated (R = H) forms, along with their respective binding affinities.

**Supplementary Figure 6**

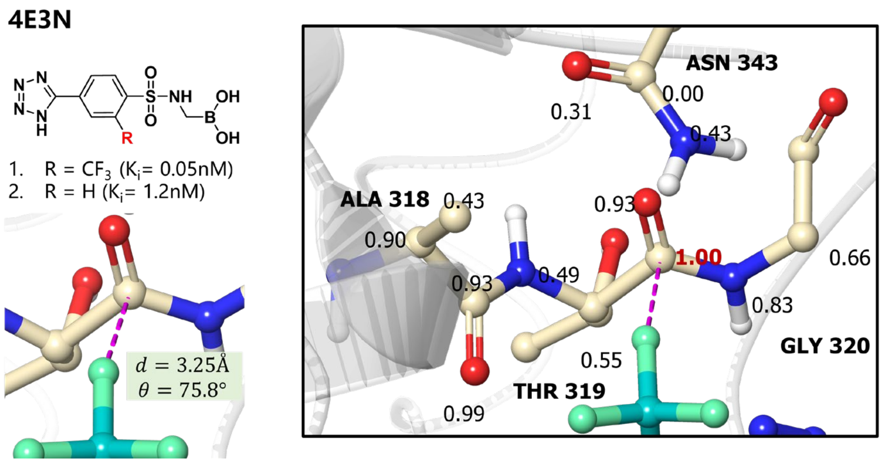

**Supplementary Figure 6.** Interpretability analysis of fluorine multipolar interactions in the 4E3N system. Left: Chemical structure of the ligand and its binding affinity data, showing enhanced potency of the fluorinated variant (R = CF3, Ki = 0.05 nM) compared to the non-fluorinated form (R = H, Ki = 1.2 nM). The fluorine multipolar interaction is indicated by a purple dashed line, with measured distance (d = 3.25 Å) and angle (θ = 75.8°). Right: Detailed view of the binding site showing weighted contributions of protein atoms within 5 Å of the fluorine atoms, where red numbers represent the weights of carbon atoms involved in the interactions. Key residues (ASN 343, ALA 318, THR 319, GLY 320) are labeled.

**Supplementary Figure 7**

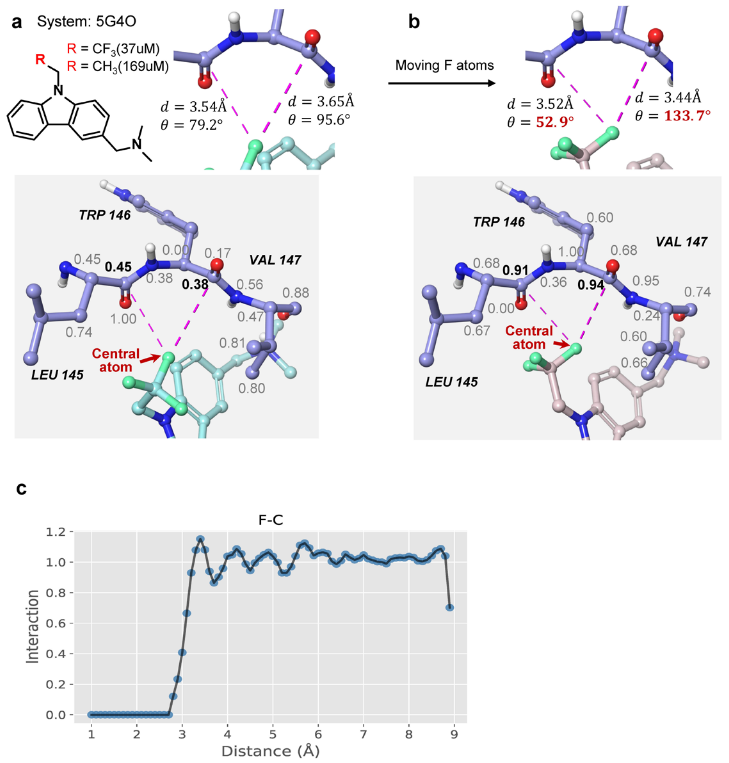

**Supplementary Figure 7. a. Interpretability analysis of the 5G4O system with PBCNet**. For a molecule with a trifluoromethyl R-group, two fluorine orthogonal multipolar interactions with the protein are identified. The binding mode and geometric constraints are illustrated, with fluorine orthogonal multipolar interactions represented by purple dashed lines. Protein atoms within 5 Å of the central fluorine atoms are annotated with weights, and bold numbers denote the weights of carbon atoms involved in the interactions. **b. Disruption of geometric constraints in fluorine orthogonal multipolar interaction**. The angles of fluorine orthogonal multipolar interactions were disrupted, changing one angle from 79.2° to 52.9° and another from 95.6° to 133.7° (both being unreasonable angles). Consequently, the weight distribution of surrounding atoms assigned by PBCNet was altered. **b. Visualization of Fluorine and Carbonyl Carbon Statistical Atom Pair Potential.** The x-axis represents the distance between pairs of atoms. y-axis is the custom defined interaction probability. A larger value indicates a higher likelihood of interaction between the atom pairs and vice versa, and when it is equal to 1 it indicates that it conforms to the background distribution, i.e., no interaction. We can find that this statistical atom pair potential does not emphasize the fluorine atoms and carbonyl carbon.

**Supplementary Figure 8**

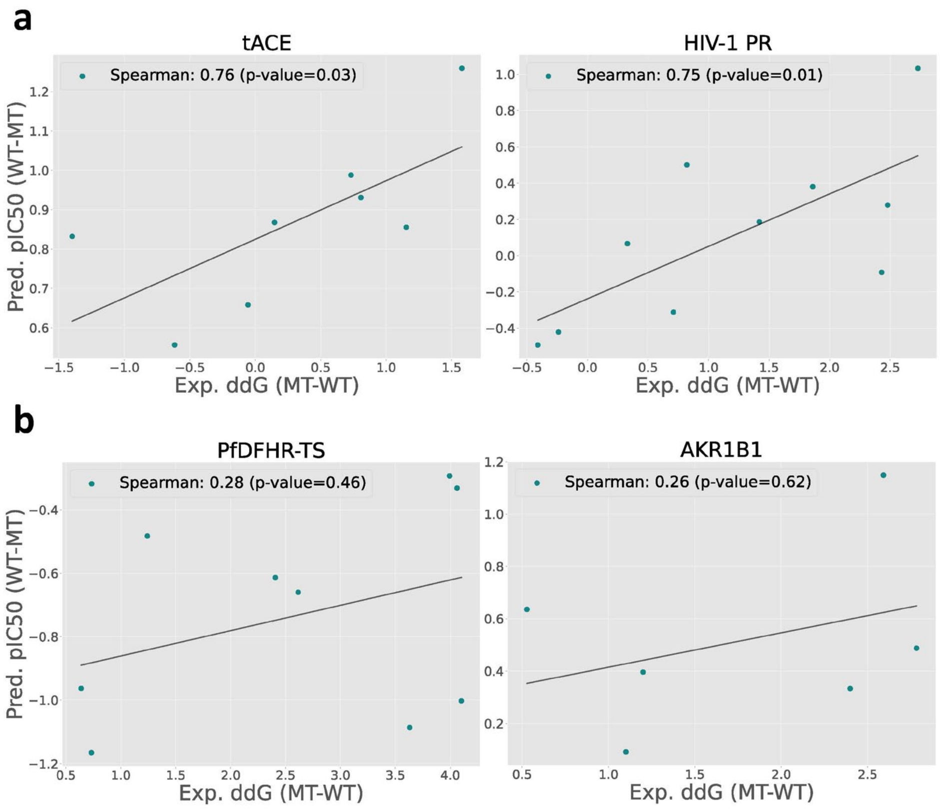

**Supplementary Figure 8. The visualization of prediction results of PBCNet2.0.** The x-axis represents experimental relative binding energies, while the y-axis shows predicted relative binding affinities. P value indicates the significance of the Spearman coefficient in an exact permutation test.

**Supplementary Figure 9**

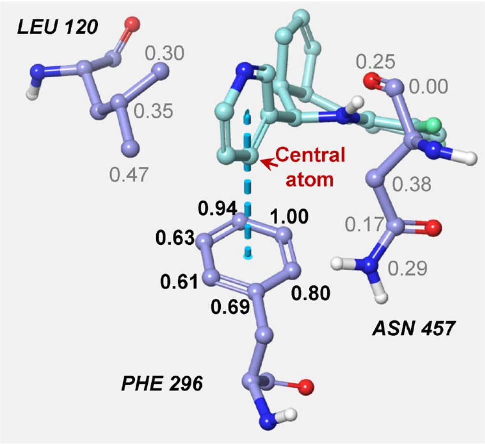

**Supplementary Figure 9. Interpretability analysis.** The $\pi-\pi$ stacking interaction is depicted as blued dashed lines, and protein atoms within 5 Å of the central carbon atom were annotated with weights, where the bold number indicates the weights associated with the carbon atom on the benzene ring of PHE296.

**Supplementary Figure 10**

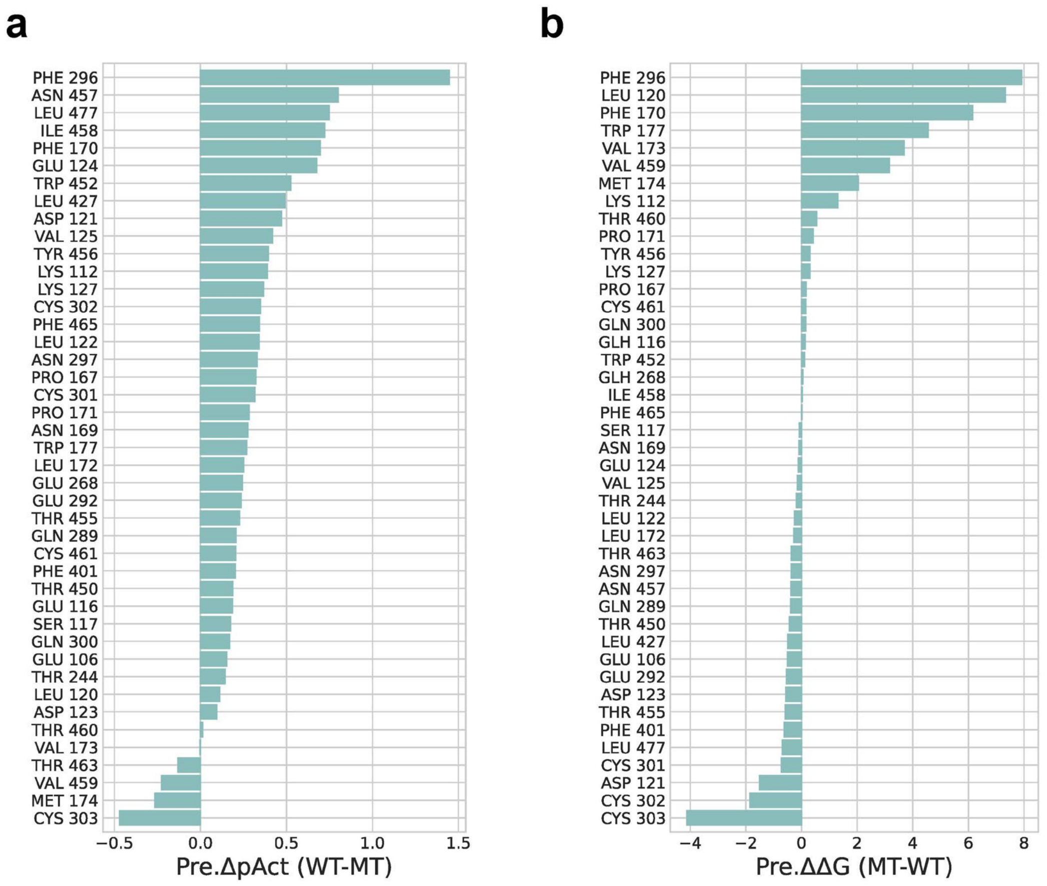

**Supplementary Figure 10. Mutation prediction results of PBCNet2.0 (a) and MM-GB/SA (b).** The x-axis represents the value of the decrease in binding affinity predicted by the model after the mutation, and the y-axis represents the mutated residue.

**Supplementary Figure 11**

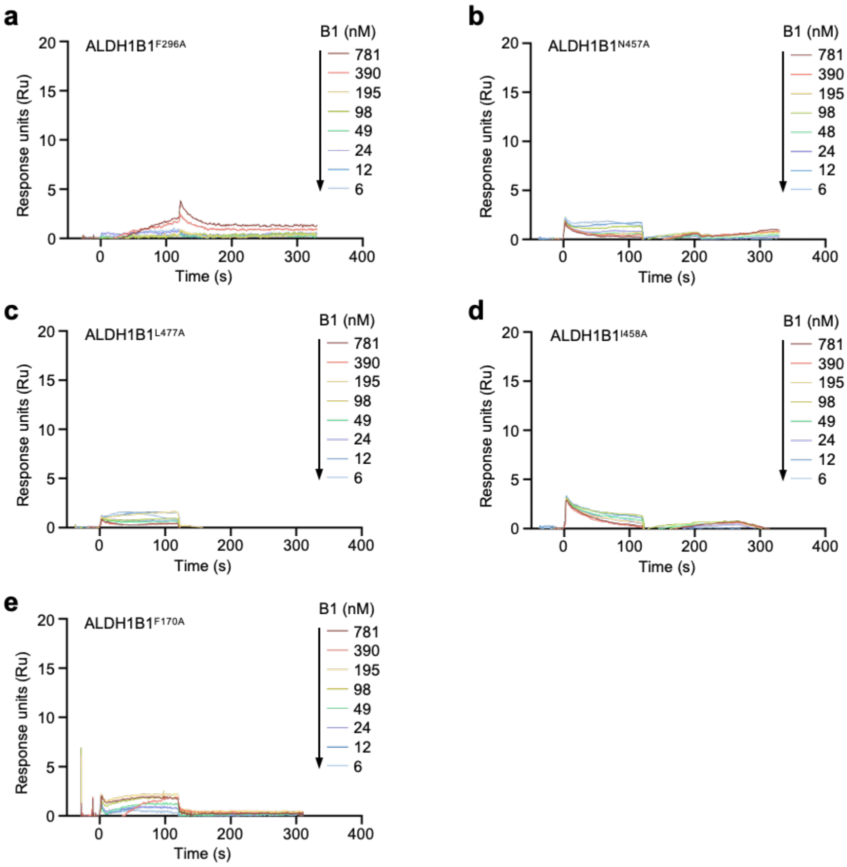

**Supplementary Figure 11 Surface plasmon resonance binding assay.** The binding kinetics measurement of B1 to various ALDH1B1 mutants were measured using SPR assay.

**Supplementary Figure 12**

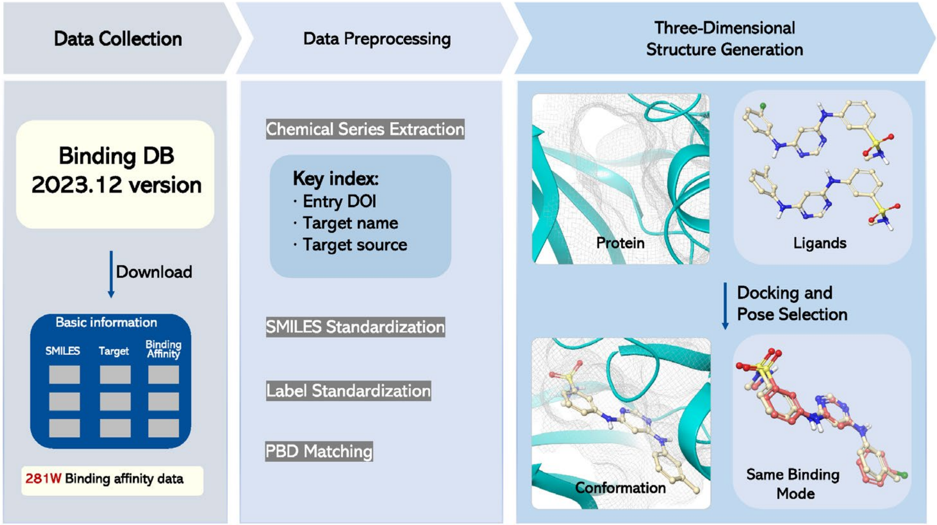

**Supplementary Figure 12.** Overview of the data preparation pipeline for PBCNet2.0 training dataset. The pipeline consists of three main stages: (1) Data Collection from BindingDB (2023.12 version) containing 2.81M binding affinity data; (2) Data Preprocessing, which involves chemical series extraction based on Entry DOI, target name, and source, standardization of SMILES structures and affinity labels, and matching with appropriate PDB structures; and (3) Three-Dimensional Structure Generation, where protein-ligand complexes are generated through Glide docking followed by pose selection based on maximum common substructure (MCS) alignment with co-crystallized ligands to ensure consistent binding modes within chemical series.

**Supplementary Figure 13**

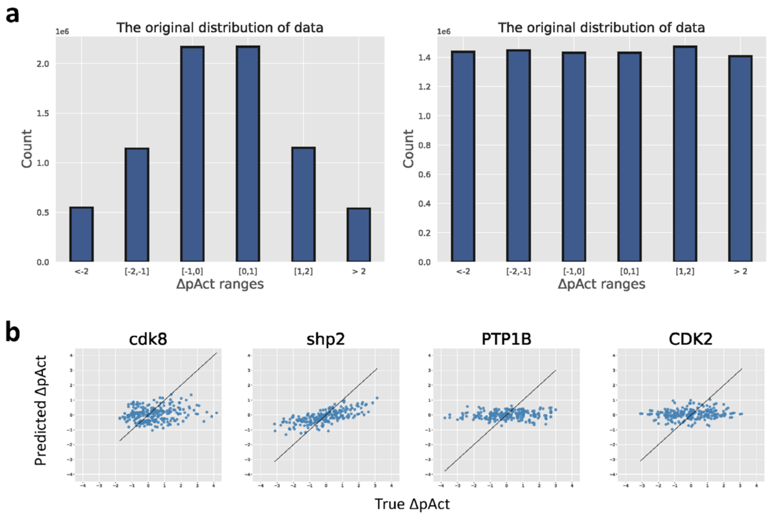

c

| Number | Training set of PBCNet | Training set of PBCNet2 |
|---|---|---|
| Chemical Series | 971 | 21722 |
| Target (Uniprot) | 960 | 1122 |
| Ligand | 20k | 280K |
| Protein-Ligand Complex | 30K | 387K |
| Ligand pairs (Training data) | 0.6M | 8.6M |

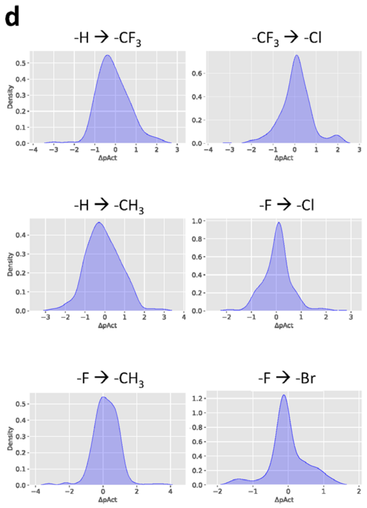

**Supplementary Figure 13.** Analysis of the training dataset. (a) Distribution of ΔpAct values in the original (left) and balanced (right) training datasets across different ranges. (b) Scatter plots showing the correlation between predicted and true ΔpAct values for four representative protein targets (cdk8, shp2, PTP1B, and CDK2). (c) Comparison table showing the number of chemical series, targets, ligands, protein-ligand complexes, and ligand pairs in the training sets of PBCNet and PBCNet2.0. (d) Distribution of activity changes (ΔpAct) for different types of R-group modifications, including -H→-CF3, -CF3→-Cl, -H→-CH3, -F→-Cl, -F→-CH3, and -F→-Br.

#### 3. Supplementary Tables

**Supplementary Table 1**

| Dataset | Target Name | Number of ligands | PDB ID |
|---|---|---|---|
| FEP1[3] | BACE | 36 | 4DJW |
| | CDK2 | 16 | 1H1Q |
| | JNK1 | 21 | 2GMX |
| | MCL1 | 42 | 4HW3 |
| | p38 | 34 | 3FLY |
| | PTP1B | 23 | 2QBS |
| | Thrombin | 11 | 2ZFF |
| | Tyk2 | 16 | 4GIH |
| FEP2[4] | CDK8 | 33 | 5HNB |
| | c-Met | 24 | 4R1Y |
| | Eg5 | 28 | 3L9H |
| | HIF-2α | 42 | 5TBM |
| | PFKFB3 | 40 | 6HVI |
| | SHP-2 | 26 | 5EHR |
| | SYK | 44 | 4PV0 |
| | TNKS2 | 27 | 4UI5 |

**Supplementary Table 1.** Protein targets in the FEP set. The first column indicates the dataset (FEP1 or FEP2) each target belongs to, the second column shows the target names, the third column presents the number of ligands for each target, and the last column displays the PDB ID of the target's crystal structure.

**Supplementary Table 2**

| Series | Target Name | Number of ligands | Uniprot ID | PDB ID | ChEMBL Assay ID |
|---|---|---|---|---|---|
| Series0_1 | TNNI3K | 9 | Q59H18 | 4YFI | 1674597 |
| Series0_2 | EHFR | 9 | P00533 | 5ZTO | 1687538 |
| Series1_1 | MAPK2 | 9 | P49137 | 2PZY | 456580 |
| Series1_2 | QPCT | 9 | Q16769 | 7CM0 | 1874814 |
| Series2_1 | Renin | 9 | P00797 | 2BKT | 321409 |
| Series2_2 | TNNI3K | 9 | Q59H18 | 4YFI | 1674597 |
| Series3_1 | WEE1 | 11 | P30291 | 1X8B | 379935 |
| Series3_2 | EPHX2 | 11 | P34913 | 4OD0 | 664904 |
| Series4_1 | PDE10A | 11 | Q9Y233 | 5SE0 | 1527711 |
| Series4_2 | EHMT1 | 11 | Q9H9B1 | 5VSD | 1705438 |
| Series5_1 | CATS | 10 | P25774 | 3IEJ | 591584 |
| Series5_2 | CHEK2 | 10 | O96017 | 4A9R | 1639625 |
| Series6_1 | F16P1 | 9 | P09467 | 5PZU | 702737 |
| Series6_2 | KIF11 | 9 | P52732 | 5ZO8 | 735125 |
| Series7_1 | FGFR1 | 9 | P11362 | 5EW8 | 1637322 |
| Series7_2 | FGFR1 | 9 | P11362 | 5EW8 | 1637327 |

**Supplementary Table 2.** Protein targets and associated data in the SAR-Diff set. Each series (Series0 to Series7) contains two targets that undergo identical R-group modifications but differ in their structure-activity relationships and scaffold structures. Column 1 shows the series designation, followed by the target name, number of ligands, UniProt ID, PDB ID for protein structure, and ChEMBL assay ID for activity data.

**Supplementary Table 3**

| Target Name | Uniprot ID | Mutation | PDB ID |
|---|---|---|---|
| EGFR | P00533 | G719C | 4WKQ |
| | | G719S | 2ITO |
| | | P753S | 4WKQ |
| | | T790M | 4WKQ |
| | | L828R+T790M | 4I22 |
| | | L858R | 2ITZ |
| | | WT | 4WKQ |
| AKR1B1 | P15121 | V47I | 2PDG |
| | | T113Y | 2PDG |
| | | F121P | 2PDG |
| | | L300P | 2PDG |
| | | S302R | 2PDG |
| | | C303D | 2PDG |
| | | WT | 2PDG |
| FldA | P0A3E0 | E57K | 2V5V |
| | | E57R | 2V5V |
| | | E57W | 2V5V |
| | | W57A | 1FLV |
| | | W57L | 1FLY |
| | | W57Y | 1FLY |
| | | Y94A | 1FLY |
| | | Y94F | 1FLY |
| | | Y94W | 1FLY |
| | | WT | 2V5V |
| HIV-1 PR | P04587 | L10F | 3GBG |
| | | M46I | 1HPV |
| | | I47V | 1HPV |
| | | G48V | 1HPV |
| | | G48V+L90M | 3BGB |
| | | I50V | 1HPV |
| | | V82A | 1HPV |
| | | V82I | 1HPV |

| | | | |
|---|---|---|---|
| | | V82F+I84V | 3BGB |
| | | I84V | 1HPV |
| | | WT | 1HPV |
| HSP90 | P02829 | K44R+K98N | 1A4H |
| | | E88G+N92L | 1A4H |
| | | L89V | 1A4H |
| | | L89V+L93I | 1A4H |
| | | N92L | 1A4H |
| | | L93I | 1A4H |
| | | K98N | 1A4H |
| | | V136M | 1A4H |
| | | WT | 1A4H |
| PfDFHR-TS | A7UD81 | A16C+S108T | 3UM8 |
| | | A16G | 3UM8 |
| | | A16S | 3UM8 |
| | | A16S+S108T | 3UM8 |
| | | A16T+S108T | 3UM8 |
| | | A16V | 3UM8 |
| | | N51I+C59R | 3UM8 |
| | | S108N | 3UM8 |
| | | S108T | 3UM8 |
| | | WT | 3UM8 |
| tACE | P12821 | E376D | 3BKL |
| | | V379S | 3BKL |
| | | V379S+V380T | 3BKL |
| | | V380T | 3BKL |
| | | F391Y | 3BKL |
| | | E403R | 3BKL |
| | | D453E | 3BKL |
| | | V518T | 3BKL |
| | | WT | 3BKL |
| LTS | P00469 | N229A | 1THY |
| | | N229C | 1THY |
| | | N229E | 1THY |
| | | N229G | 1THY |

| | | | |
|---|---|---|---|
| | | N229I | 1THY |
| | | N229L | 1THY |
| | | N229M | 1THY |
| | | N229Q | 1THY |
| | | N229S | 1THY |
| | | N229T | 1THY |
| | | WT | 1THY |

**Supplementary Table 3.** Protein targets and associated data in the mutation test set. The table lists target proteins, their UniProt IDs, and mutation types, where mutations are denoted by the original amino acid, position, and mutated amino acid (e.g., G719C indicates a mutation from Glycine to Cysteine at position 719). WT (Wild Type) indicates no mutation in the protein sequence. Solved crystal structures of mutant proteins in the PDB database were preferentially used for molecular docking. In cases where crystal structures of mutant proteins were not available, mutant protein structures from the MdrDB[5] dataset were used.

**Supplementary Table 4**

| Target Name | PDB ID | Activity before modification (IC50/Kd/Ki)/μM | Activity after modification (IC50/Kd/Ki)/μM | ΔpAct |
|---|---|---|---|---|
| p53-Y220C[6] | 5G4O | 169 | 37 | 0.66 |
| Thrombin[7] | 1OYT | 0.31 | 0.057 | 0.75 |
| p38[8] | 3FLN | 0.106 | 0.014 | 0.88 |
| DYRK1A[9] | 5A4L | 36 | 3.9 | 0.97 |
| Renin[10] | 3OQK | 0.773 | 0.063 | 1.10 |
| Procaspase-6[11] | 4NBL | 2.9 | 0.47 | 0.79 |
| gp120[11] | 4DKO | 3.7 | 0.76 | 0.69 |
| β-lactamase[11] | 4E3N | 0.0012 | 0.00005 | 1.38 |
| Menin-MLL[11] | 4OG6 | 0.39 | 0.197 | 0.30 |
| Macrocyclic menin-MLL[11] | 4I80 | 0.0267 | 0.0047 | 0.75 |

**Supplementary Table 4.** Protein targets and associated data in the F-Opt test set. The table presents 10 pairs of protein-ligand complexes where modifications from -H or -CH3 to -F or -CF3 led to improved binding affinities. Activity values ($IC_{50}$/Kd/Ki in μM) before and after modification are shown, along with their difference in negative logarithm form (ΔpK). These cases were selected based on documented fluorine orthogonal multipolar interactions and available co-crystal structures.

**Supplementary Table 5**

| Atom Feature | Description |
| --- | --- |
| Atom type | Range (0,128) |
| Degree | Range (0,11) |
| Formal charge | Range (-5,6) |
| Chirality | unspecified, tetrahedral CW, tetrahedral CCW, other |
| Hydrogens | Range (0,9) |
| Hybridization | sp, sp2, sp3, sp3d, sp3d2 |
| Implicit valence | Range (0,7) |
| Vdw radius | Range (0,5) |
| Aromaticity | Whether this atom is part of an aromatic system |

**Supplementary Table 5.** Atomic features

**Supplementary Table 6**

| Bond Feature | Description |
| --- | --- |
| Bond type | single, double, triple, aromatic, distance |
| Bond distance | 0 to 5 Å |

**Supplementary Table 6.** Bond features

8. Goldstein, D. M. *et al.* Discovery of 6-(2,4-Difluorophenoxy)-2-[3-hydroxy-1-(2-hydroxyethyl)propylamino]-8-methyl-8H-pyrido[2,3-d]pyrimidin-7-one (Pamapimod) and 6-(2,4-Difluorophenoxy)-8-methyl-2-(tetrahydro-2H-pyran-4-ylamino)pyrido[2,3-d]pyrimidin-7(8H)-one (R1487) as Orally Bioavailable and Highly Selective Inhibitors of p38α Mitogen-

Activated Protein Kinase. *J. Med. Chem.* **54**, 2255–2265 (2011).

9. Rothweiler, U. *et al.* Probing the ATP-Binding Pocket of Protein Kinase DYRK1A with Benzothiazole Fragment Molecules. *J. Med. Chem.* **59**, 9814–9824 (2016).

10. Discovery and optimization of a new class of potent and non-chiral indole-3-carboxamide-based renin inhibitors. *Bioorganic & Medicinal Chemistry Letters* **20**, 6268–6272 (2010).

11. Pollock, J. *et al.* Rational Design of Orthogonal Multipolar Interactions with Fluorine in Protein–Ligand Complexes. *J. Med. Chem.* **58**, 7465–7474 (2015).
